# iPSC-derived *NF1*-*CDKN2A*-PRC2 deficient neural crest mimics MPNST glial-to-neuro-mesenchymal transition and uncover new therapeutic opportunities

**DOI:** 10.1101/2025.08.13.670072

**Authors:** Itziar Uriarte-Arrazola, Míriam Magallón-Lorenz, Juana Fernández-Rodríguez, Jiajing Zhang, Emily Lee, Sara Ortega-Bertran, Edgar Creus-Bachiller, Judit Farrés-Casas, Kelli M. Wilson, Crystal McKnight, Katlin Recabo, Ignacio Blanco, Héctor Salvador, Cleofé Romagosa, Conxi Lázaro, Helena Mazuelas, Bernat Gel, Marc Ferrer, Meritxell Carrió, Eduard Serra

## Abstract

Neurofibromatosis Type 1 (NF1) predisposes to peripheral nerve tumor development. Commonly, the progression from a benign plexiform neurofibroma (PNF) towards a deadly malignant peripheral nerve sheath tumor (MPNST) involves a poorly understood glial-to-mesenchymal transition and the sequential loss of *NF1, CDKN2A*, and polycomb repressive complex 2 (PRC2). Using an iPSC-derived neural crest (NC) model, we reproduced this malignant transformation through gene editing. NF1-CDKN2A double-knockout (2KO) NCs retained glial differentiation capacity and formed neurofibroma-like tumors *in vivo*, requiring inactivation of p14ARF and p16INK4a. Additional PRC2 loss (3KO) disrupted pluripotency and induced mesenchymal stem cell-like features in iPSCs an NCs. 3KO NCs suffered a global chromatin reprograming that silenced *SOX10* preventing gliogenesis and activated neuro-mesenchymal programs. Gene signatures characterizing this glial-to-neuro-mesenchymal transition were recapitulated in human PNF-ANNUBP-MPNST tumors. 3KO NC spheres formed MPNST-like tumors *in vivo* upon nerve engraftment, genuinely mimicking an early-stage MPNST. We used the developed 3D NC spheroid models for the discovery of drugs targeting MPNSTs by performing a high-throughput screening of an epigenetic compound library. We found that poly(ADP-ribose) polymerase inhibitors (PARPi) exhibit selective efficacy in PRC2-deficient NC spheroids. We confirmed that Olaparib-Selumetinib combination in a MPNST PDX mouse model was well tolerated and significantly suppressed tumor growth.

## INTRODUCTION

Malignant peripheral nerve sheath tumors (MPNSTs) are aggressive soft tissue sarcomas arising from peripheral nerve cells. Nearly half of all cases are associated with Neurofibromatosis type 1 (NF1), a hereditary cancer syndrome that predisposes to the development of benign and malignant tumors.^1^ MPNSTs account for 3–10% of all soft tissue sarcomas, and NF1 patients face a 10–15% lifetime risk of developing them, making MPNSTs the leading cause of mortality in this collective.^2,3^ The 5-year survival rate is poor, particularly in the NF1 context.^2,4^ No effective therapies exist besides a timely surgery.^5^

In NF1, MPNST normally develops through an initial loss of several tumor suppressor genes (TSGs) in a stepwise manner.^6,7,8^ Benign plexiform neurofibromas (PNFs) arise in large nerves from neural crest (NC)–derived Schwann cell precursors (SCP) after *NF1* inactivation.^9^ The additional loss of the complete *CDKN2A* locus, characterizes the formation of a distinct pre-malignant nodular lesion termed atypical neurofibromatous neoplasm of uncertain biological potential (ANNUBP).^10,11^ Both lesions share a predominant glial component but with different cell microenvironment. The progression toward MPNST involves the loss of polycomb repressive complex 2 (PRC2), a chromatin remodeler complex driving transcriptional repression during development,^12^ through inactivation of its components *SUZ12* or *EED*.^13,8^ This transition is concurrent with a shift towards a mesenchymal identity of tumor cells. *TP53* is also frequently inactivated in MPNSTs, but at a lesser extent.^8,14^ After TSG loss, MPNSTs undertake an extensive genomic reorganization, including copy number alterations and structural variants.

Several *in vitro* and *in vivo* models have been developed for PNF, ANNUBP and MPNST, associated with NF1, to study their biology and identifying potential new therapies. However, only few models mimic the progression from benign to malignant NF1 tumors.^15,16,17^ High-throughput screening (HTS) platforms provide a powerful framework to evaluate drug responses, both individually and in combination. Previous HTS efforts in NF1-associated tumors have predominantly relied on two-dimensional (2D) culture models, including immortalized primary PNF-derived Schwann cells (SCs)^18,19^ and MPNST cell lines.^20^

In this work, we established a human iPSC-derived NC-based model platform with the sequential inactivation of *NF1, CDKN2A*, and *SUZ12*, to recapitulate the genetic events underlying PNF-MPNST progression. The resulting models captured the glial-to-mesenchymal transition characteristic of neurofibroma-to-MPNST transformation and triple-knock out NC spheroids formed early-stage MPNSTs upon nerve engraftment. Leveraging this platform, we performed a HTS of an epigenetics compound library, identifying PARP inhibitors as selectively active in PRC2-deficient models. Moreover, the PARPi–MEKi combination significantly suppressed tumor growth in a patient-derived MPNST xenograft model. Together, this 3D NC model platform provides a tractable system to study MPNST biology and discover new treatment possibilities.

## RESULTS

### *In vivo* neurofibroma-like tumor formation of *NF1*(−/−) *CDKN2A*(−/−) neurofibromaspheres requires the loss of p14ARF

We first targeted *CDKN2A* in an *NF1*-deficient iPSC line (1KO_D12) using CRISPR-Cas9.^21^ Since *CDKN2A* encodes both p14ARF (p14) and p16INK4a (p16) via alternative reading frames, we designed sgRNAs targeting exon 2: one selectively disrupting p16, and another inactivating both p14 and p16 (Figure 1A). We successfully generated two *NF1*(−/−) *CDKN2A* p16(−/−) (2KO (p16)), and three *NF1*(−/−) *CDKN2A* p14p16(−/−) (2KO (p14p16)), iPSC lines (Figure S1A, File S1). Clonal generation and expansion with p16 inactivated and p14 WT was less efficient than with both proteins inactivated, and one of the 2KO (p16) iPSC lines (2KO_A11) showed abnormal phenotype, slow proliferation, and loss of pluripotency (Figure S1B). We confirmed both, newly CRISPR generated mutations at *CDKN2A* and previous *NF1* pathogenic variants, by sanger sequencing (File S1) and western blot (Figure S1C, S1D). iPSC lines were characterized by pluripotency marker expression (OCT3/4) and their ability to differentiate into homogeneous populations of NC cells, as assessed by cytometry analysis (p75 and Hnk1) and immunocytochemistry (TFAP2 and SOX10) (Figure 1B). p16 is detected by western blot at the level of NC but not at iPSC. Thus, we characterized the impact of *CDKN2A* loss on NC physiology. We performed proliferation and ploidy assays, but no differences were observed compared to the parental 1KO cell line, only deficient in *NF1* (Figure S1E, S1F). Whole-genome sequencing (WGS) confirmed an isogenic diploid (2n) genome without structural or any other relevant alterations of the 2KO_F9 (p14p16) cell line used for subsequent analysis (Figure S1G).

**Figure 1.**
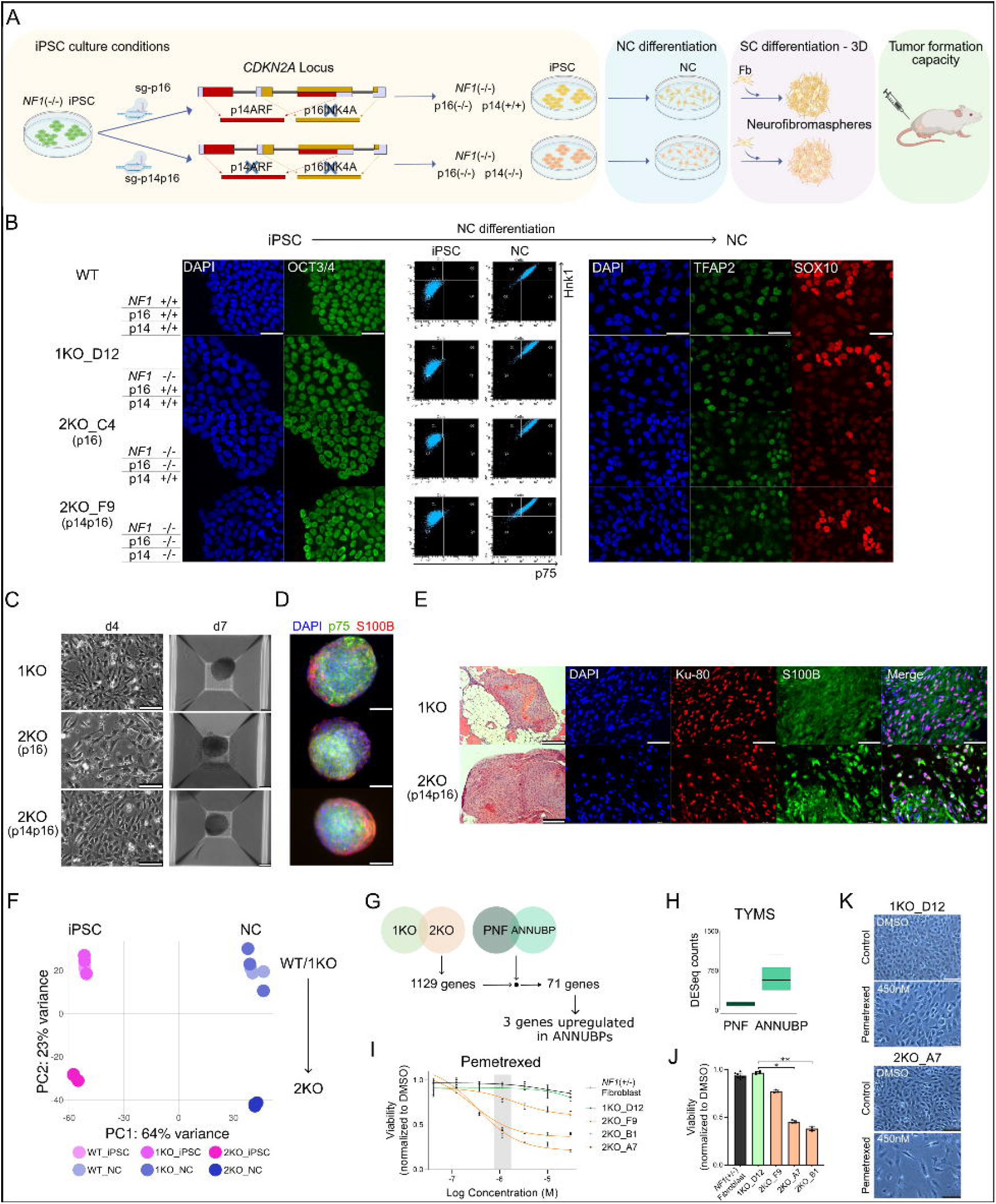
Generation of double knock out *NF1*(−/−) *CDKN2A*(−/−) iPSC lines. **A)** Schematic representation of the editing strategy used to knock out the *CDKN2A* locus in *NF1*(−/−) iPSCs, followed by neural crest (NC) formation and Schwann cell (SC) differentiation to generate neurofibromaspheres and assess tumorigenic potential. Sg: sgRNA; Fb: PNF-derived fibroblast. **B)** Left panel: OCT3/4 expression in iPSCs: wild type WT (*NF1* +/+); 1KO (*NF1* −/−); 2KOp16 (*NF1* −/− *CDKN2A* p16INK4a −/−); and 2KOp14p16 (*NF1* −/− *CDKN2A* −/−). Middle panel: Flow cytometry analysis of p75 and Hnk1 before (iPSC) and after (NC) NC differentiation. Right panel: TFAP2 and SOX10 expression at NC stage. Scale bar: 50 µm. **C)** Left: Micrographs of NC cells after 4 days (d4) of SC differentiation. Scale bar: 100 µm. Right: representative neurofibrompaspheres at day 7 (d7) of SC differentiation, containing also PNF-derived fibroblasts. Scale bar: 100 µm. **D)** Representative immunofluorescence images showing p75 (green) and S100B (red) expression in neurofibromasheres. Scale bar: 100 µm. **E)** Characterization of the tumors grown after engraftment of neurofiromaspheres in the sciatic nerve of nude mice. Left panel: H&E staining. Scale bar: 200 µm. Right panel: Immunofluorescence images for Ku-80 and S100B. Scale bar: 50 µm. **F)** Principal component analysis (PCA) of global gene expression of WT, 1KO and 2KO cell lines at iPSC and NC stage. One WT in triplicate and three 1KO and 2KO cell lines are shown. **G)** Schematic representation of upregulated genes shared by 2KO NCs and ANNUBPs. **H)** *TYMS* expression in PNFs and ANNUBPs. **I)** Cell viability dose–response curves for pemetrexed treatment after 96h in 1KO_D12 NC, 2KO NC, an *NF1*(+/−) fibroblast cell line, assessed by CellTiter-Glo assay. Mean ± SD of technical replicates (n ≥ 3). **J)** Cell viability after 1.11 µM pemetrexed treatment in cell lines from (I). Mean ± SD of technical replicates (n ≥ 3). Kruskal-Wallis multiple comparison test: *p < 0.05, **p < 0.01. **K)** Representative micrographs showing density and morphology changes between 1KO_D12 NC and 2KO_A7 NC cell lines and lack of senescence via X-Gal staining (blue) after 96h of 450 nM pemetrexed treatment. Scale bar: 100 µm.

We then tested the tumor formation capacity of 2KO (p16) and 2KO (p14p16) cell lines. We generated neurofibromaspheres by differentiating NCs towards cells of the SC lineage, and mixed them with primary neurofibroma fibroblasts allowing them to form spheres for up to 14 days (Figure 1C, 1D).^22^ Spheres were engrafted into the sciatic nerve of nude mice and analyzed after four months. Tumors developed in 10/15 injections with the 1KO line and in 9/15 with 2KO (p14p16). H&E and S100B staining confirmed neurofibroma-like morphology in both genotypes without evident histological differences. Ku80 staining confirmed the human origin of S100B positive cells (Figure 1E). In contrast, 2KO (p16) spheres displayed markedly reduced tumorigenicity, with outgrowths in only 3/15 injections that were significantly smaller than those from 1KO or 2KO (p14p16). These findings indicate that wild-type p14 in 2KO (p16) cells impairs proper neurofibroma formation.

### 2KO NC are sensitive to thymidylate synthase inhibition

Next, we interrogated the impact of *CDKN2A* loss on overall gene expression. We performed RNA-seq of biological triplicates of 1KO and 2KO (p14p16) at iPSC and NC stage, and performed a principal component analysis (PCA) of the global expression (Figure 1F). p14 and p16 loss had a clear impact on gene expression. We identified several differentially overexpressed genes in 2KO cells compared to 1KO, shared by iPSCs and NCs. 71 of these genes were also differentially expressed when comparing PNFs and ANNUBPs (Figure 1G, File S2). However, in ANNUBPs, 2KO cells may account for less than 50% of tumor cells,^7^ and only three of the 71 genes were found to be overexpressed in ANNUBPs, two of them being related to immune infiltrate. The third overexpressed gene was *TYMS*, encoding for thymidylate synthase (TS) (Figure 1H). The described synthetic lethality between loss of *CDKN2A* and TS inhibition^23^ prompted us to test the sensitivity of 2KO NC cells to TS inhibitors compared to 1KO NCs. We used the TS inhibitor Pemetrexed on 1KO and 2KO NC cultures (Figure 1I-1K) and observed a significantly higher sensitivity of 2KO NCs to *TYMS* inhibition (Figure 1I, 1J). Although senescence was ruled out as the cause of this sensitivity, we observed marked morphological changes in the cells, including increased size, aberrant nuclei, and enlarged cytoplasm in 2KO cells (Figure 1K).

### Inactivation of *NF1, CDKN2A* and PRC2 in iPSCs exhibit a biologically imposed order

Next step involved the inactivation of PRC2, through targeted editing of *SUZ12*, in 1KO, 2KO (p16) and 2KO (p14p16) iPSCs. We were able to assess PRC2 functional loss by directly interrogating the absence of the trimethylation of lysine 27 of histone 3 (H3K27me3), using a specific antibody (Figure 2A). We performed multiple transfections in each cell line (Figure 2B) successfully achieving *SUZ12* editing in three genotypically distinct iPSCs (Figure 2C). However, when trying to expand the different edited clones, we did not succeed for 1KO and 2KO (p16), that is, for those iPSCs with preserved p14ARF function (Figure S2A, S2B). However, we were able to expand clones from 2KO (p14p16) edited iPSCs and perpetuate them at the NC stage (see below), obtaining 3KO cell lines with no functional *NF1, CDKN2A* (p14p16) and *SUZ12* (Figure 2B, 2D). These results suggested that the order of TSG inactivation in *NF1*(−/−) iPSCs required a biologically imposed specific sequence: first the complete inactivation of the *CDKN2A* locus (involving p14 and p16) and then the loss of PRC2.

**Figure 2.**
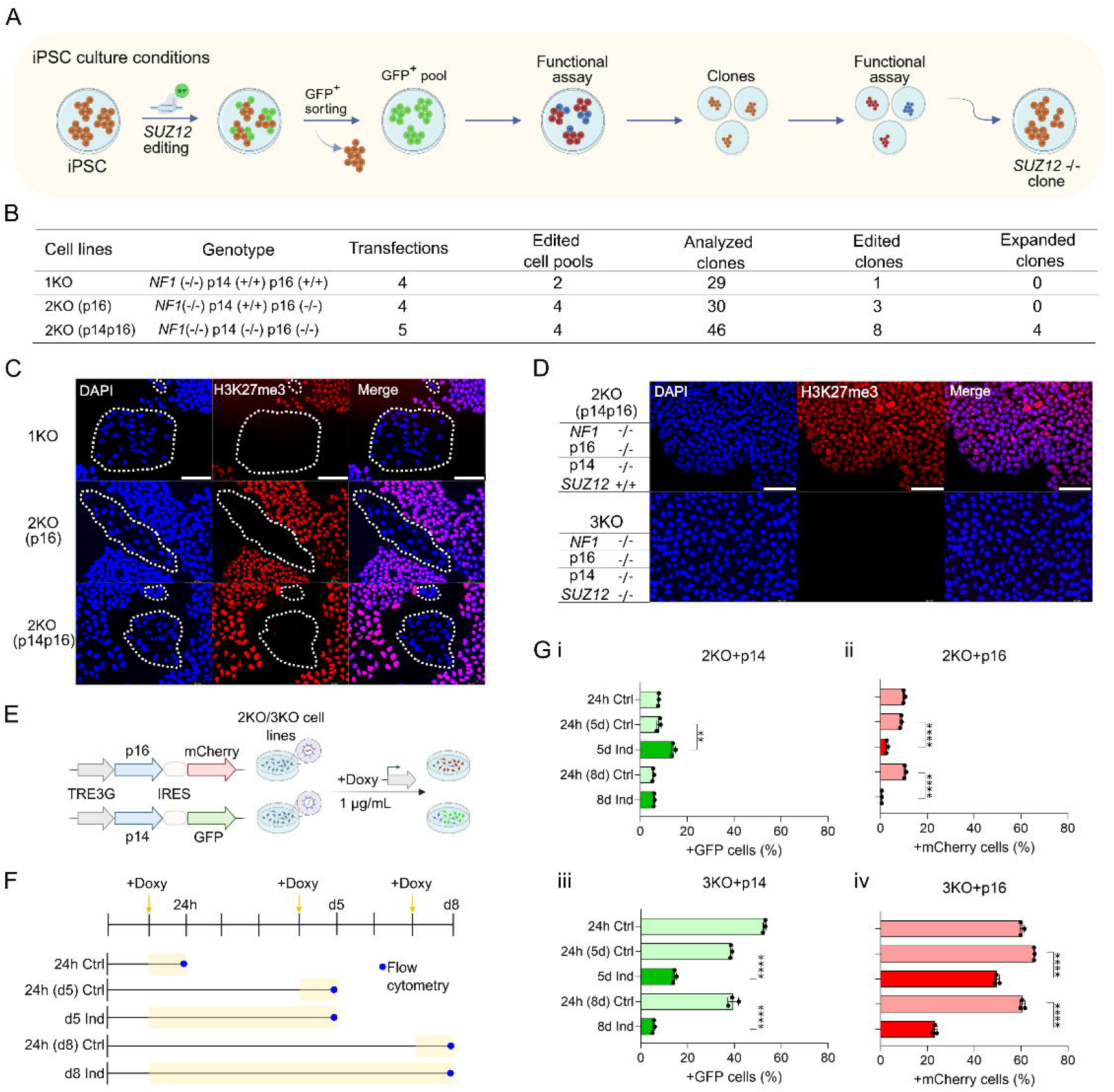
A biological constraint for the inactivation of *NF1, CDKN2A* and PRC2 in iPSCs. **A)** Schematic representation of the editing strategy used to knock out *SUZ12* in iPSCs and screening workflow for edited clones. **B)** Quantification of editing efficiency and clonal progression at each stage of the *SUZ12* editing workflow. **C)** Immunofluorescence detection of H3K27me3 in transfected 1KO and 2KO cell pools. *SUZ12*-edited cells are indicated by white dashed lines. Scale bar: 100 µm. **D)** Immunofluorescence of H3K27me3 in 2KO (p14p16) parental line and an expanded 3KO clonal line. Scale bar: 100 µm. **E)** Diagram of lentiviral doxycycline-inducible systems for p16 and p14 re-expression in 2KO and 3KO cell lines. **F)** Schematic representation of the experimental design for inducible expression of p14 and p16 in 2KO and 3KO cells. Transduced cells were treated with doxycycline either at day 5 (d5) or day 8 (d8) and expression levels of p16 (mCherry) and p14 (GFP) were assessed by flow cytometry. Control cells were treated with doxycycline for 24 hours prior to each measured time point (d1, d5 and d8). **G)** Flow cytometry analysis of GFP+ and mCherry+ cell populations at d5 and d8 post-doxycycline induction. (**i**) p14 reexpression in 2KO cell line, (**ii**) p16 re-expression in 2KO cell line, (**iii**) p14 re-expression 3KO cell line, (**iv**) p16 re-expression 3KO cell line. Data represent mean ± SD from three independent experiments. Unpaired two-tailed t tests: **, p < 0.01; ****, p < 0.0001.

To confirm these findings, we tested doxycycline-inducible re-expression of p14 or p16 in 2KO (p14p16) and 3KO NC cells using lentiviral bicistronic vectors co-expressing p14-GFP or p16-mCherry (Figure 2E). Fluorescence was quantified at baseline (24h) and after 5 and 8 days post-initial induction. Expression was also induced 24h before day 5 and 8 in independent cultures and measured at these days, to control for lentiviral silencing (Figure 2F). Fluorescent cell percentages differed between 2KO and 3KO lines (Figure 2G) consistent with variable lentiviral silencing. In 2KO (p14p16) NC cells, p14 re-expression did not affect viability, whereas p16 re-expression markedly reduced survival (Figure 2Gi-ii). As 2KO (p14) iPSCs were not generated, the suggested inability to derive this line could not be directly confirmed. In 3KO cells, re-expression of either p16 or particularly p14 severely compromised viability (Figure 2Giii-iv) aligning with the inability to expand PRC2-edited 3KO lines from 2KO (p16). Altogether these results support the existence of a biologically imposed order of inactivation of TSGs, as in the PNF-ANNUBP-MPNST progression, and support the previous finding that MPNSTs bear the complete inactivation of the *CDKN2A* locus, involving both p14ARF and p16INK4a proteins.^24^

### PRC2 loss in *NF1*(−/−) *CDKN2A*(−/−) iPSCs induces loss of pluripotency, mesenchymal identity acquisition, and can be perpetuated as neural crest

Inactivation of *SUZ12* in 2KO (p14p16) iPSCs resulted in clones that, under iPSC conditions, underwent spontaneous differentiation and progressively acquired a mesenchymal-like phenotype, eventually leading to proliferative arrest and senescence (Figure 3A–C). Flow cytometry revealed expression of mesenchymal stem cell (MSC) markers (CD73, CD13, and CD44) consistent with a shift toward an MSC-like identity (Figure 3D). Immunostaining confirmed OCT3/4 downregulation and SOX9 upregulation in 3KO cells compared to 2KO iPSCs (Figure 3E). 3KO cells were non-viable under standard iPSC culture conditions, nevertheless, we were able to perpetuate them under NC culture conditions and establish three independent non-perishable 3KO cell lines (Figure 3F, 3G, Figure S3A, File S1). Flow cytometry confirmed expression of NC markers HNK1 and p75 (Figure 3H). However, 3KO NCs also displayed elevated levels of MSC markers compared to their 1KO and 2KO counterparts (Figure 3H, Figure S3B). Proliferation rates and ploidy of 3KO NCs remained basically unchanged compared to 1KO and 2KO (Figure S3C, S3D). All 3KO NC cell lines also exhibited an unaltered 2n genome assessed by WGS analysis (Figure S3E). Notably, immunofluorescence for SOX10 and SOX9—key NC transcription factors (TFs)— revealed an inverse relationship across 1KO, 2KO, and 3KO NCs: while SOX10 expression gradually diminished and was absent in 3KO NCs, SOX9 reached a maximum protein expression in these cells (Figure 3I, Figure S3F). Moreover, 3KO NCs exhibited a drastic downregulation of glial genes (*SOX10, NGFR, PLP1*), and an upregulation of mesenchymal markers (*SOX9, TWIST1, PRRX1*) (Figure 3J) compared to 2KO NCs, as assessed by bulk-RNAseq analysis, suggesting a glial-to-mesenchymal shift.

**Figure 3.**
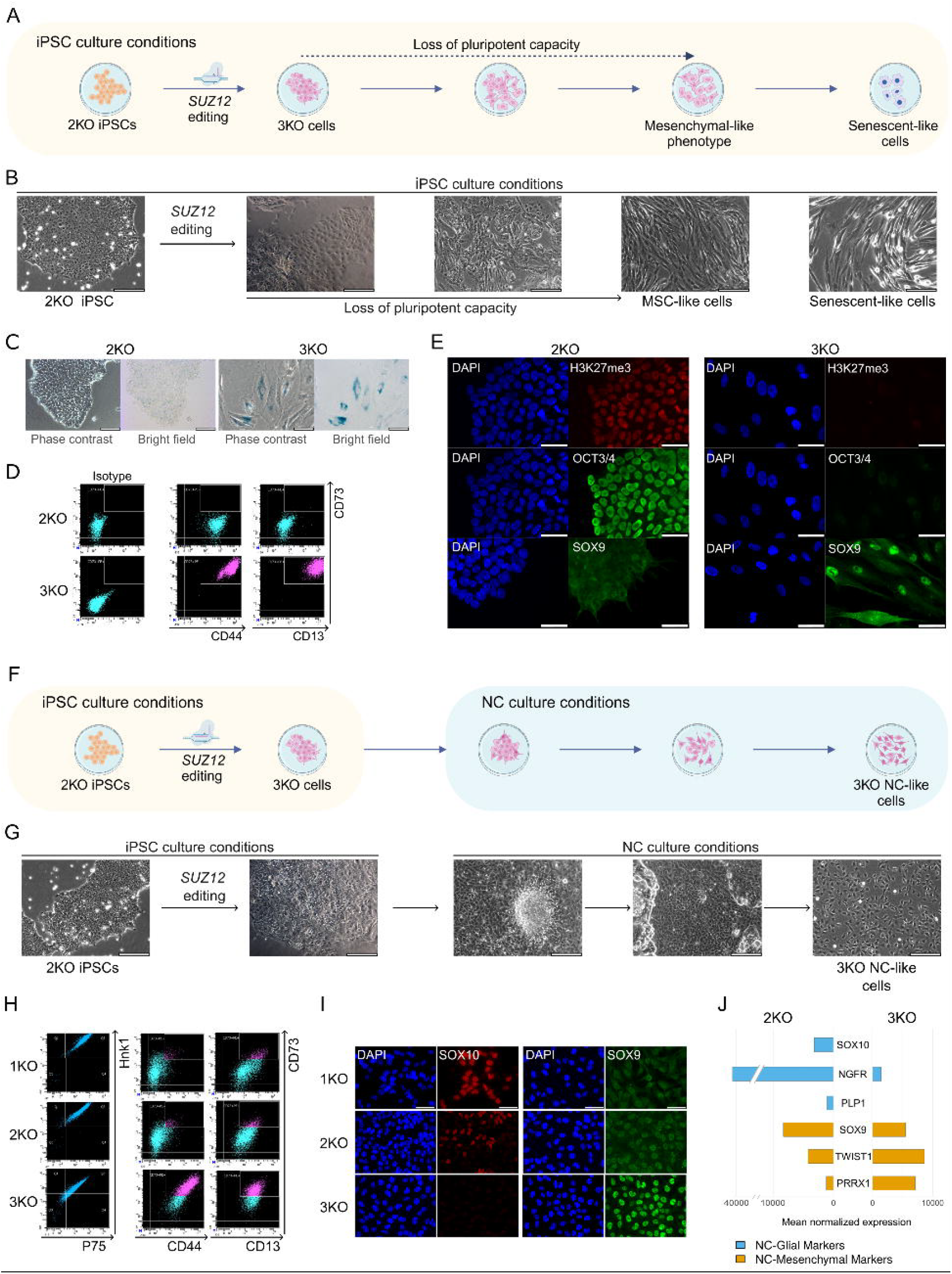
Loss of PRC2 function in 2KO iPSCs induces loss of pluripotency and transition toward a mesenchymal-like identity. **A)** Schematic representation of the acquired cell fate of 2KO cells after *SUZ12* editing in iPSC culture conditions. **B)** Representative micrographs illustrating loss of pluripotency, acquisition of an MSC-like phenotype, and senescence onset in 3KO *SUZ12*-edited cells. **C)** Representative micrographs of senescence analysis (X-Gal staining in blue) in 2KO and 3KO cell lines. Scale bar: 100 µm. **D)** Flow cytometry analysis of MSC markers CD13, CD44 and CD73. **E)** Immunofluorescence images illustrating pluripotency (OCT3/4), PRC2 function (H3K27me3) and mesenchymal identity (SOX9) in 2KO and 3KO cell lines. Scale bar: 50 µm. **F)** Schematic representation of the strategy used to perpetuate 3KO *SUZ12*-edited cells under NC culture conditions. **G)** Representative micrographs showing NC-like phenotype acquisition of 3KO cells under NC culture conditions. Scale bar: 200 µm. **H)** Flow cytometry for NC markers (p75 and Hnk1) and MSC markers (CD13, CD44 and CD73) in 1KO, 2KO and 3KO cell lines. **I)** Immunofluorescence of SOX10 and SOX9 expression in 1KO, 2KO and 3KO cell lines. Scale bar: 50 µm. **J)** Expression of selected NC-glial (*SOX10, NGFR, PLP1*) and NC-mesenchymal (*SOX9, TWIST1, PRRX1*) markers in 2KO (n=3) and 3KO (n=3) cell lines.

### 3KO NC spheroids generate human MPNST-like tumors upon nerve engraftment in nude mice

To functionally test these identity changes and the potential of 3KO NCs to form tumors *in vivo*, we performed a tumor formation assay by engrafting 3KO NC-like spheroids in contact with the sciatic nerve of nude mice, together with 1KO and 2KO neurofibromaspheres for comparative purposes. After four months, mice were sacrificed. We identified large, highly nucleated human tumors at the injection site of the 3KO NC-like spheroids, adjacent to the nerve (Figure 4A). These tumors exhibited numerous mitotic figures and a histology compatible with an MPNST by H&E staining, as assessed by an expert NF1 pathologist. We confirmed the human origin of these cells by Ku-80 staining (Figure 4A). Neurofibromaspheres derived from 1KO and 2KO NCs generated neurofibroma-like tumors as previously described^21^ (Figure 4B, Figure 1E). Immunostaining for H3K27me3, SOX10, S100B, vimentin, and Ki67 was performed (Figure 4B). 1KO and 2KO tumors retained H3K27me3 and expressed SOX10/S100B, whereas 3KO tumors lacked these markers, confirming PRC2 loss and loss of neurofibroma-like identity. All tumors were Vimentin and Ki67 positive, with slightly higher Ki67 in 3KO tumors, suggesting increased proliferation These findings demonstrated that 3KO NC spheres can generate early-stage MPNSTs, establishing a relevant model for studying malignant progression and drug testing.

**Figure 4.**
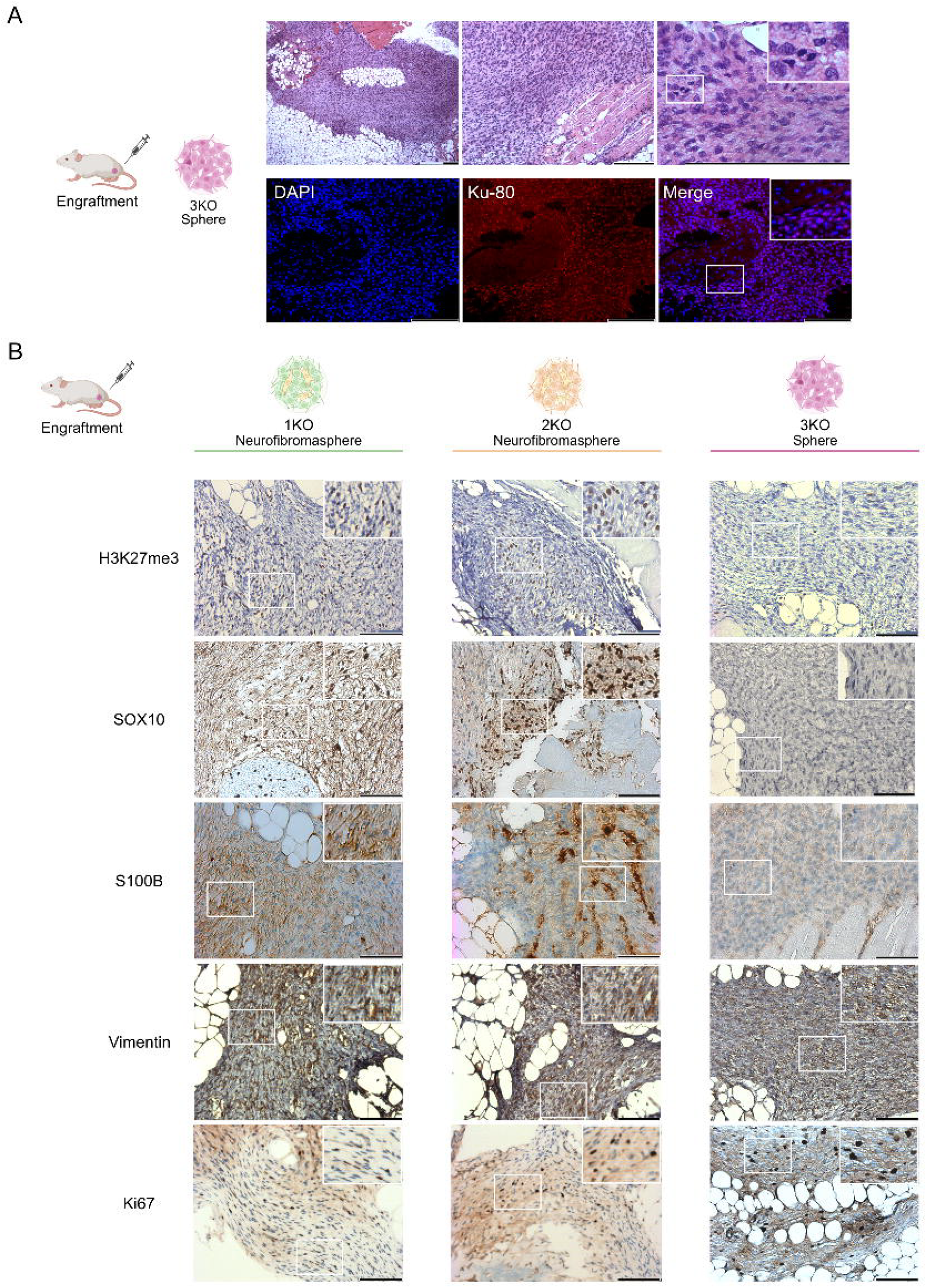
3KO NC spheroids generate human MPNST-like tumors *in vivo* mimicking early-stage MPNST development. **A)** Characterization of tumors arising from 3KO NC spheroid engraftment into the sciatic nerve of nude mice. Upper panel: representative hematoxylin and eosin (H&E) images showing high cellularity and mitotic activity. Scale bar: 200 µm. Lower panel: Ku-80 immunofluorescence with DAPI nuclear counterstaining, showing human origin. Scale bar: 200 µm. **B)** Immunohistochemical comparison of 1KO, 2KO, and 3KO tumor models showing H3K27me3, SOX10, S100B, Vimentin, and Ki67 expression. Scale bar:100 µm.

### MPNST cell identity is highly impacted by PRC2 inactivation and MSC-like features

3KO NC cells model early MPNST development (TSG inactivation, 2n genome) whereas established MPNST cell lines represent a late stage of disease (TSG inactivation, rearranged genome). To identify molecular features gained during this progression, we compared both *in vitro* models, focusing the analysis on how PRC2 loss and mesenchymal identity acquisition shape MPNST cell identity (Figure 5A).

**Figure 5.**
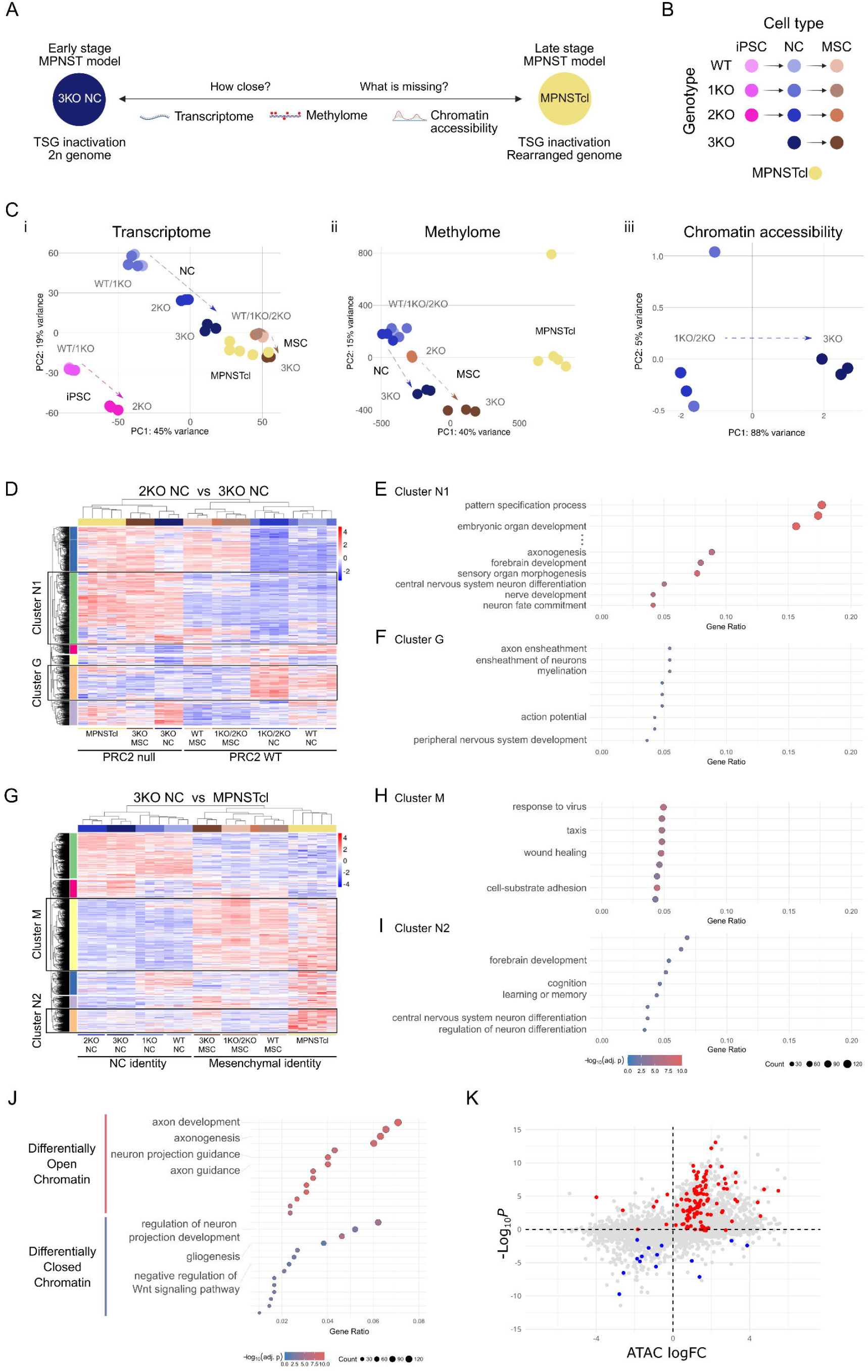
MPNST phenotype is markedly affected by PRC2 inactivation, the presence of neurogenesis and MSC-like traits and gliogenesis downregulation. **A)** Schematic overview of the transcriptomic and epigenomic comparison between 3KO NC and MPNST cell lines (MPNSTcl) *in vitro* models. **B)** Schematic representation showing the cell lines used for the omics analyses. **C)** Principal component analysis (PCA) of cell types in (B), comparing (**i**) transcriptome, (**ii**) methylome and (**iii**) chromatin accessibility. Arrows indicate the impact of genotype on the different cell type identities. **D)** Heatmap showing the expression patterns across all samples of genes differentially expressed between 2KO and 3KO NC. See also File S3. **E)** Biological processes identified by the enrichment analysis of genes in cluster N1 (see panel D). See also File S3. **F)** Biological processes identified by the enrichment analysis of genes in cluster G (see panel D). See also File S3. **G)** Heatmap showing the differential expression analysis of 3KO NC vs. MPNST cell lines in all samples. See also File S5. **H)** Biological processes identified by the enrichment analysis of genes in cluster M (see panel G). See also File S5. **I)** Biological processes identified by the enrichment analysis of genes in cluster N2 (see panel G). See also File S5. **J)** Enrichment analysis of genes with differentially open (top) and closed (bottom) chromatin accessibility regions in PRC2-deficient NC cell lines compared to PRC2-WT NC. See also File S6. **K)** Correlation plot integrating differential gene expression and ATAC-seq data in PRC2-wild-type (1KO, 2KO) and PRC2-mutant (3KO) NC cell lines. Cluster N1 and N2 neurogenesis signature genes highlighted in red; Cluster G gliogenesis signature highlighted in blue.

For that, we first differentiated 3KO NC lines and their parental 2KO, 1KO and WT isogenic NC lines into MSCs (Figure 5B, Figure S4A). Differentiating 3KO NCs into MSCs led to early proliferative arrest and senescence (Figure S4B), preventing the generation of stable 3KO MSC lines. However, sufficient MSCs were obtained to enable multiple downstream analyses. We included all different genotypes and cell identities, from iPSCs to NC and MSCs, and five MPNST cell lines bearing the inactivation of *NF1, CDKN2A* and *SUZ12* (Figure 5B).

We performed transcriptome (RNA-seq) for all cell types; methylome (EPIC array) for NCs, MSCs and MPNST cell lines, and chromatin accessibility analysis (ATAC-seq) for NCs. PCAs evidenced a profound contribution of both PRC2 loss and MSC expression in defining the identity of MPNST cell lines, with a clear impact of both factors on transcriptome and methylome profiles (Figure 5Ci, 5Cii). In addition, the loss of PRC2 also had a profound effect on chromatin accessibility in NCs, as expected (Figure 5Ciii). We compared the expression of 2KO and 3KO NC lines, to assess for the impact of PRC2 loss at the NC stage. Differentially expressed genes were visualized in a heatmap alongside MPNST cell lines, NCs and MSC genotypes (Figure 5D, File S3). Unsupervised clustering separated samples into two groups based on PRC2 status, with a shared set of upregulated genes (Cluster N1) identified in 3KO NCs, 3KO MSCs, and MPNST cell lines. Gene enrichment analysis of cluster N1 revealed strong associations with neurogenesis-related processes (Figure 5E, Figure S5A, File S3). In addition, we identified cluster G, enriched in genes related to gliogenesis, that was downregulated in 3KO NCs compared to PRC2 WT NCs (Figure 5F, Figure S6A-B). We also compared 3KO NCs with established MPNST cell lines to identify features absent in 3KO NCs key for becoming full MPNST cells. Unsupervised clustering of differentially expressed genes separated cell lines with NC identity, from those with a mesenchymal identity (MSCs and MPNST cell lines), regardless of their genotype (Figure 5G, File S5). We identified a set of genes upregulated in cells with mesenchymal identity (Cluster M). Enrichment analysis of cluster M revealed multiple processes associated with mesenchymal identity, including 73 genes involved in wound healing (Figure 5H, File S5). Furthermore, we identified an additional group of genes upregulated only in MPNST cell lines (Cluster N2). Interestingly, enrichment analysis of this cluster identified biological processes that were also related to neurogenesis (Figure 5I, Figure S5B, File S5), indicating its importance for MPNST identity.

Global expression analysis highlighted a glial-to-neuro-mesenchymal transition from 2KO to 3KO NC cells, mimicking features of ANNUBP-MPNST progression. To investigate the role of chromatin remodeling in this transition, we analyzed ATAC-seq data. Enrichment analysis of differential chromatin accessibility between PRC2-null (3KO NC) and PRC2-WT (1KO and 2KO NC) supported the global opening of neurogenesis genes and the partial closing of gliogenesis genes (Figure 5J, File S6). To complete this view, we plotted together gliogenesis and neurogenesis genes from clusters G, N1 and N2 (Figure 5D-I), in a graph correlating gene expression and chromatin accessibility. Gliogenesis genes were downregulated, partly by epigenetic silencing, while neurogenesis-related genes were predominantly upregulated by an open chromatin conformation (Figure 5K).

### 3KO neural crest cells recapitulate a glial-to-mesenchymal transition: gliogenesis impairment and expression of neuro-mesenchymal genes

We aimed to better understand the mechanisms driving this cell identity switch and its physiological implications. We first evaluated the ability of 3KO NCs to differentiate towards SCs, testing gliogenesis capacity. SC differentiation was induced in 1KO, 2KO, and 3KO NCs for 21 days^25^ (Figure 6A). 1KO and 2KO differentiating SCs exhibited both SOX9 and SOX10 expression, along with the SC lineage markers S100B and p75. Contrarily, 3KO differentiating cells preserved SOX9 expression but were not able to express SOX10, a master TF of gliogenesis, and any other SC marker, uncovering a loss of glial differentiation capacity. To determine whether *SOX10* inhibition was a direct epigenetic change produced by PRC2 loss, we analyzed chromatin accessibility in *SOX10* promoter. We found a close chromatin conformation in 3KO NCs compared to the open conformation of 1KO and 2KO NCs (Figure 6Bi), explaining gliogenesis shut down. In contrast, *SOX9* TF promoter remained under an open chromatin conformation in 3KO NC cells (Figure 6Bii).

**Figure 6.**
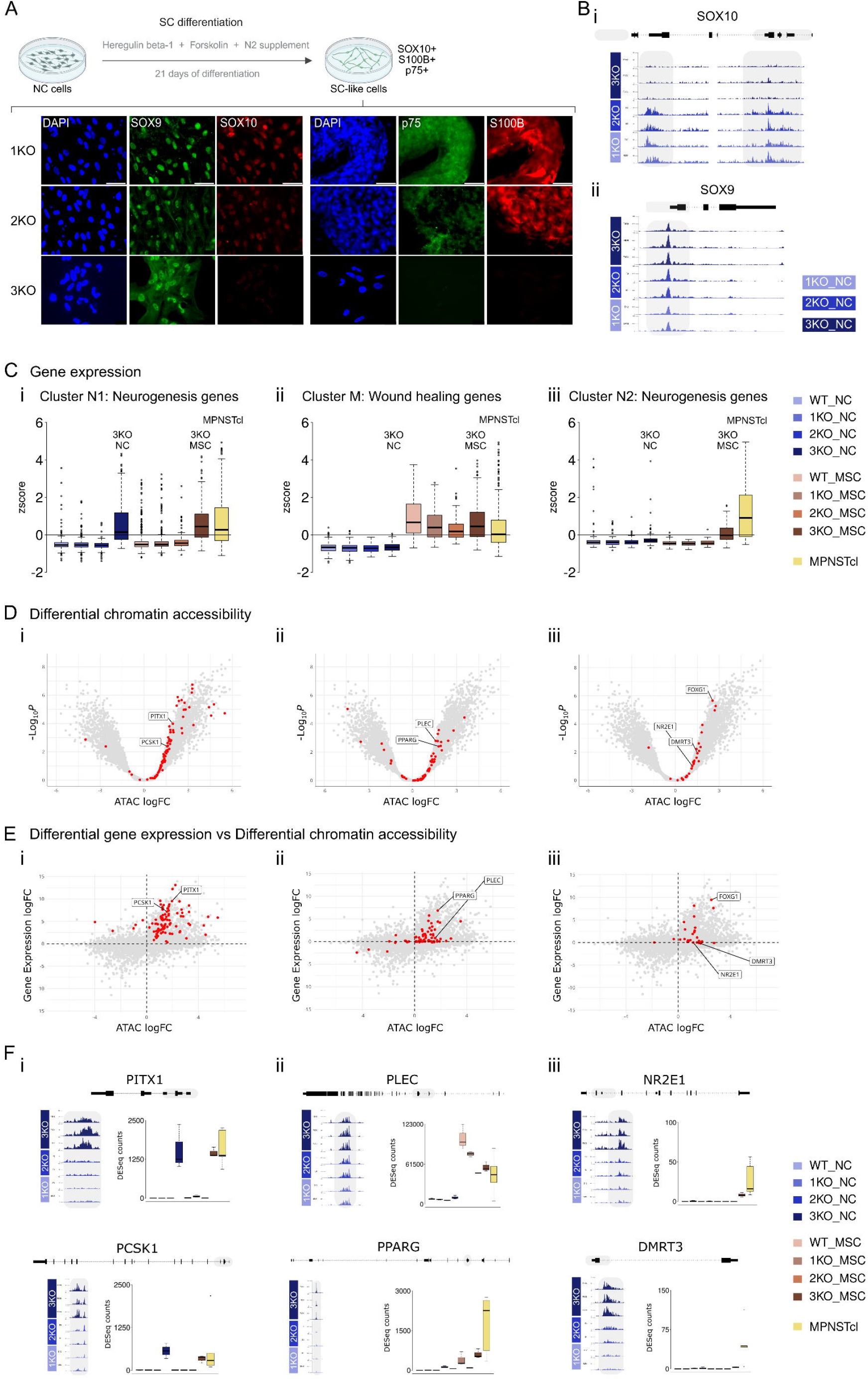
Loss of SC differentiation capacity in 3KO NC cell lines due to *SOX10* epigenetic closure. Neurogenesis genes: part of MPNST cell identity. **A)** NC-SC differentiation scheme and immunofluorescence images of SOX9, SOX10 and NC-SC markers (p75 and S100B) after 21 days of SC differentiation in 1KO, 2KO and 3KO cell lines. Scale bar: 50 µm. **B)** ATAC-seq chromatin accessibility plot in 1KO, 2KO and 3KO cell lines of (**i**) *SOX10* locus, showing epigenetic silencing in 3KO cell lines and (**ii**) *SOX9* locus, showing no changes between NC cell lines. **C)** Boxplot showing the expression z-score of (**i**) Cluster N1 neurogenesis gene signature (Figure 5D; 5E); (**ii**) Cluster N2 neurogenesis gene signature (Figure 5G; 5I); (**iii**) Cluster M wound healing gene signature (Figure 5G; 5H). **D)** Volcano-plot showing differential chromatin accessibility of PRC2 WT (1KO, 2KO) compared to PRC2-null (3KO) NC cell lines. (**i**) Cluster N1 neurogenesis genes highlighted in red; (**ii**) Cluster N2 neurogenesis genes highlighted in red; (**iii**) Cluster M wound healing genes highlighted in red. **E)** Correlation plot integrating differential gene expression and ATAC-seq data in PRC2 wild-type (1KO, 2KO) compared to PRC2-null (3KO) NC cell lines. (**i**) Cluster N1 neurogenesis genes highlighted in red; (**ii**) Cluster N2 neurogenesis genes highlighted in red; (**iii**) Cluster M wound healing genes highlighted in red. **F)** ATAC-seq chromatin accessibility plot in 1KO, 2KO and 3KO cell lines and normalized expression in all cell lines of (**i**) *PITX1* and *PCSK1* genes from Cluster N1 neurogenesis genes; (**i**) *NR2E1* and *DMRT3* genes from Cluster N2 neurogenesis genes; (**i**) *PLEC* and *PPARG* genes from Cluster M wound healing genes.

Furthermore, to obtain a mechanistic insight of the expression of neuro-mesenchymal gene signatures identified and their importance to conform MPNST cell identity, we performed a global analysis of these signatures together with their chromatin accessibility status. Neurogenesis genes of cluster N1 (Figure 5D, 5E) showed a consistent high expression in 3KO cell lines only—3KO NCs, 3KO MSCs and MPNST cell lines (Figure 6Ci). In 3KO NCs, their promoters exhibited a differentially open chromatin conformation compared to PRC2-WT NCs (Figure 6Di) and a positive correlation between chromatin opening and expression (Figure 6Ei). Thus, these neurogenesis genes were poised by PRC2. Wound healing-related genes of cluster M (Figure 5G, 5H) were only expressed in cells with mesenchymal identity, regardless of the genotype (Figure 6Cii). Accordingly, although chromatin was preferentially open in 3KO NC vs PRC2-WT NCs (Figure 6Dii), many of the genes did not change expression comparing all NC genotypes (Figure 6Eii), indicating that their expression in MPNSTS is dependent on mesenchymal-related factors. Finally, another group of neurogenesis genes within the N2 cluster (Figure 5G, 5I), were basically expressed only in MPNST cell lines (Figure 6Ciii). Again, most of these genes opened their chromatin upon PRC2 loss in 3KO NCs (Figure 6Diii) but many of them did not change the expression in 3KO NCs vs 2KO (Figure 6Eiii). All together indicated that in addition to PRC2 loss and mesenchymal-associated factors, other genes like those of the cluster N2 signature, required additional unknown factors to be expressed in MPNSTs, also reinforcing neurogenesis as part of the MPNST cell identity. Chromatin accessibility and expression of representative genes of the different signatures are shown in Figure 6F and Figure S7. Overall, results supported that NF1-associated 3KO MPNST identity is deeply influenced by losing glial differentiation capacity and the expression of developmental neuro-mesenchymal programs.

### Glial-to-neuro-mesenchymal signatures identified in 3KO NCs also differentiate benign from malignant NF1-associated PNS tumors

We then assessed the relevance of the different signatures identified in the iPSC-derived NC models, to the identity of the NF1-associated tumors and their benign-to-malignant progression, using global expression analysis. We first analyzed bulk RNA-seq data from primary PNF-derived SCs, PNFs, ANNUBPs, and MPNSTs, focusing on previously defined gliogenesis, neurogenesis genes and wound healing genes (Figure 7A). Gliogenesis genes from cluster G were predominantly expressed in SCs and benign tumors. Cluster N1 neurogenesis genes segregated into three groups: (i) genes (e.g., *NGF, VIM, ITGA2*) upregulated in benign tumors and SCs; (ii) genes (e.g., *FOXD1, GATA3, PCSK1*) enriched in MPNSTs, absent in SCs and with low expression in PNFs/ANNUBPs, likely due to the presence stromal cells; and (iii) genes (e.g., *NKX2-2, PITX1, FOXG1*), along with Cluster N2 genes (e.g., *NR2E1, DMRT3, FOXB1*), expressed only in MPNSTs. Selected wound healing genes from cluster M (File S5, in bold) were also predominantly expressed in MPNSTs. These transcriptional profiles reflect a glial-to-neuro-mesenchymal transition underlying NF1 tumor progression.

**Figure 7.**
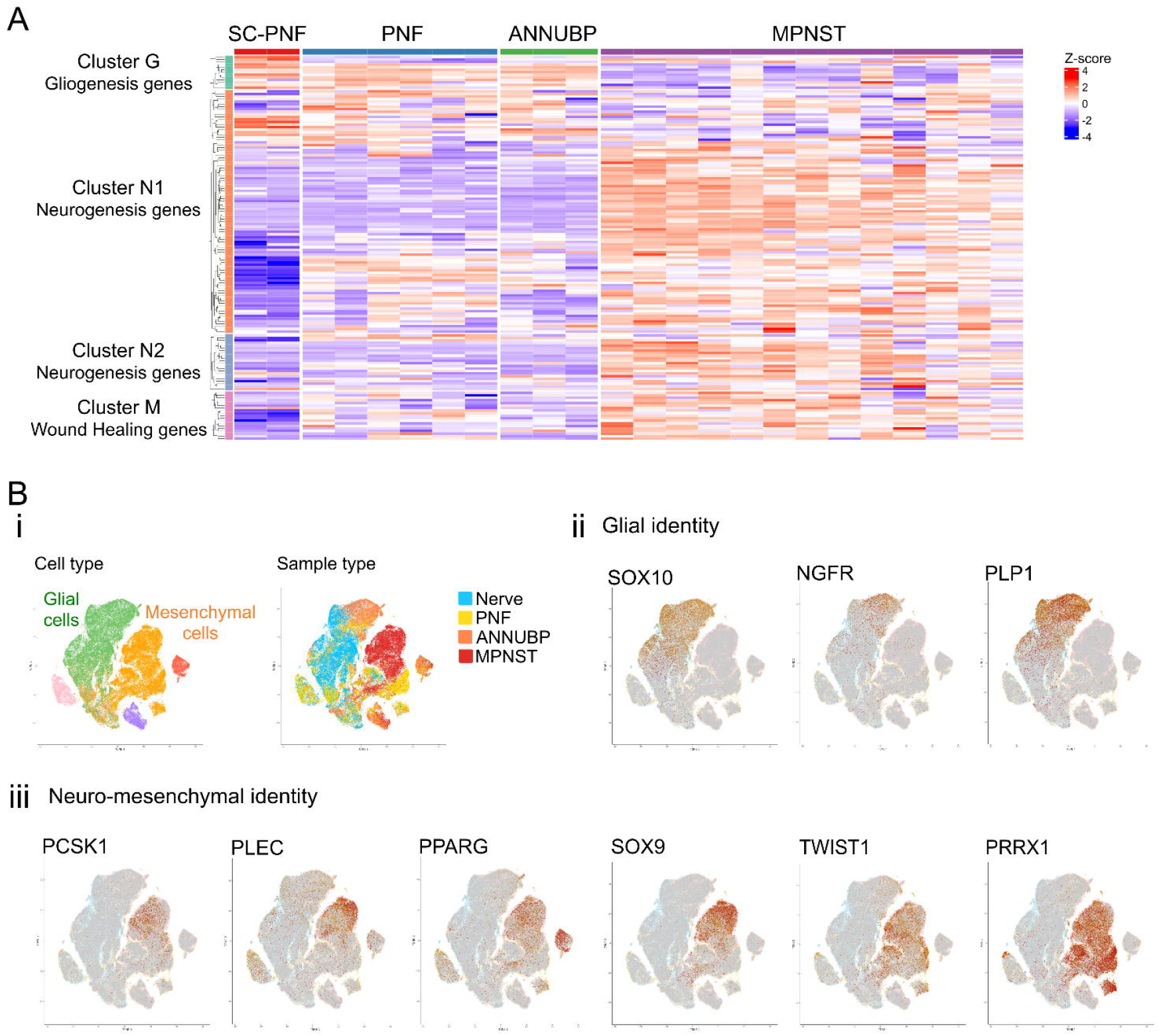
Glial to neuro-mesenchymal transition in human NF1-related tumors. **A)** Heatmap showing identified gliogenesis and neurogenesis gene signatures across human NF1-associated PNF-SCs, PNF, ANNUBP, and MPNST samples. **B)** TSNE plots representing single-cell RNA-seq data from nerve, PNF, ANNUBP and MPNST samples. (**i**) Representation of cell type composition and sample type distribution. (**ii**) Gliogenesis genes *SOX10, NGFR* and *PLP1* are exclusively expressed in de glial cell component. (**iii**) Neurogenesis gene (*PCSK1*) and mesenchymal genes (*PLEC, PPARG, SOX9, TWIST1* and *PRRX1*) are exclusively expressed in mesenchymal cells and mostly in MPNSTs.

Furthermore, we investigated this transition characterized by the identified signatures, using single cell data obtained from nerves, PNFs, ANNUBPs and MPNSTs, to better understand their relevance but according to the different cell types within tumors (Figure 7B). Expression of gliogenesis genes like *SOX10, NGFR* and *PLP1* identified in cluster G, was restricted to the glial component of all tissues and tumors analyzed, particularly expressed in ANNUBP samples (Figure 7Bii). Contrarily, expression of identified neuro-mesenchymal genes from clusters N1, N2 and M, like *PCSK1, PLEC* and *PPARG*, were restricted to the mesenchymal component of MPNSTs, like other MPNST markers like *SOX9, TWIST1* and *PRRX1* (Figure 7Biii), further supporting the importance of identified gene signatures for MPNST identity.

### High-throughput screening of epigenetic modulators identifies candidate compounds and synergistic combinations with MEK inhibition in 3KO NC spheroid 3D model

After showing that the NC-based models developed mimic many features of malignant NF1-associated PNS tumors and considering the global epigenetic remodeling produced by PRC2 loss, we next used these NC models to identify epigenetic modulators as potential therapies for MPNST treatment. We conducted a quantitative high-throughput screen (qHTS) using the NCATS Pharmacologically Active Chemical Toolbox (NPACT) Epigenetics library, comprising of 339 small-molecule modulators of epigenetic targets (Figure S8A-B) (File S7). We used the different NC models developed (1KO, 2KO, 3KO) grown as 3D spheroids and an additional 3KO cell line carrying a heterozygous *TP53* background (3KO+) because *TP53* is mutated in some MPNSTs (Figure S8B, Figure S9). We tested the compounds in dose-response across all 3D NC spheroid models and measured cell viability with CellTiter-Glo 3D (CTG) after 48h of treatment. Spheroids of NF1 patient-derived fibroblasts were used for assessing general toxicity (Figure S8B, Figure S10). We confirmed that these spheroid models generated robust assays, with Z⍰-factor values across all cell lines ranging between 0.75 and 0.82 (Table S2). In general, a large subset of compounds (250/339) showed no activity in 3KO cell lines (IC_50_≥13 µM) or showed poor or ambiguous dose-response curves (224/339). Of the 339 compounds tested, 15 showed broad cytotoxicity (**Figure 8A**, File S8).

**Figure 8.**
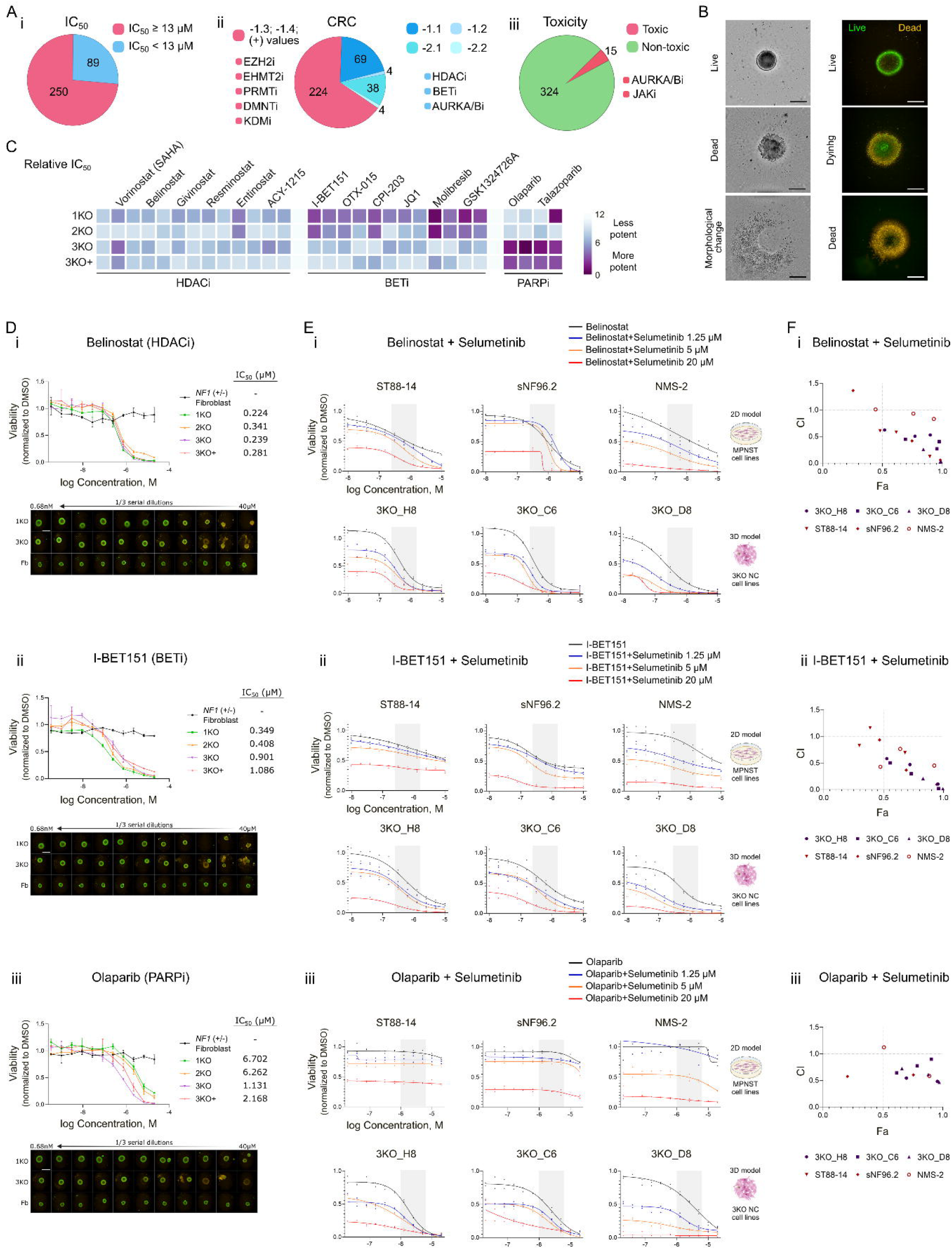
High-throughput screening and validation of epigenetic compound modulators, as single agents and in combination with Selumetinib (MEKi), in isogenic 3D KO NC spheroid models and MPNST cell lines. **A)** Sumary of qHTS results presented as pie charts. (**i**) Activity: classification based on IC_50_ and CRC values in 3KO spheroids. Active compounds (blue): CRC −1.2, −1.2, −2.1 and −2.2; and IC_50_<13 µM. Inactive compounds (red): CRC −1.3, −1.4 and positive values; and IC_50_≥13 µM. (**ii**) Toxicity: compounds with IC_50_ < 1 µM in control fibroblast spheroids or similar IC_50_ values across all cell lines (IC_50_ in isogenic spheroids > 5-fold IC_50_ in the fibroblast control spheroids). See also Table S3. **B)** Representative phase contrast (left) and Live/Dead fluorescence (right) micrographs of 3KO spheroids after 48h of treatment. Scale bar: 250 µm. **C)** Heatmap showing the relative IC_50_ of BETi, PARPi and HDACi across isogenic cell lines. Dark purple indicates higher potency (row data in File S9). **D)** Upper panel: Dose– response CellTiter-Glo (CTG) cell viability graphs of the four isogenic NC cell lines and control fibroblasts, grown in 3D, together with corresponding IC_50_ values. Data correspond to eleven dose–response concentrations; each point shows the mean ± SD from two independent replicates. Bottom panel: Live/dead micrographs for each dose in 1KO, 3KO, and control fibroblasts (Fb). Scale bar: 500 µm. **E)** Dose–response CTG cell viability plots of candidate compounds in combination with Selumetinib (MEKi) for each 3KO cell line and each MPNST cell line. Shaded areas indicate three concentrations above and below the IC_50_ of the compounds in 3KO_H8 used to calculate Combination index (CI). Data correspond to eleven dose-response concentrations; each point shows the mean ± SD from two independent replicates. **F)** CI plots of the shaded doses from panel (E), showing the fraction affected (Fa) versus CI for the 3KO (purple) and MPNST (red) cell lines. Fa represents the fraction of cell death induced by drug treatment, ranging from 0 (no cell death) to 1 (complete cell killing). CI values <0.9 indicate a synergistic interaction, values between 0.9 and 1.1 indicate an additive effect, and values >1.1 indicate antagonism.

Based on the initial qHTS curve classifications, we selected compounds displaying activity profiles (curve response class (CRC) values of –1.1, –1.2, –2.1, or –2.2) ^26^ together with high potency (IC_50_ < 13 µM) and spheroid morphology changes (**Figure 8B**), as well as a few generally cytotoxic compounds, as toxicity controls. We selected 77 compounds and performed 11-point dose–response curves to refine IC_50_ estimations (File S9) and also performed live-cell imaging (live/dead assays) to evaluate morphological changes not captured by CTG (Figure S11). Interestingly, although changes in the IC_50_ value were generally below three-fold, nearly all BET inhibitors (BETi) exhibited greater activity in 1KO (or 2KO) cells compared to 3KO and 3KO+ cells (**Figure 8C**). However, as for most compounds, HDAC inhibitor (HDACi) Belinostat and BETi I-BET151 exhibited comparable effects across all four isogenic lines (**Figure 8C, Figure 8D i–ii**). In contrast, selective loss of viability for PRC2-deficient spheroids was observed for PARP inhibitors (PARPi), with 3 out of 4 PARPi showing more than a three-fold change in IC_50_ in PRC2-mutant cells compared to PRC2 WT cells (**Figure 8C**). As an example, the effects of Olaparib are illustrated in **Figure 8Diii**. Additional effects were evident across several compounds (Figure S12).

Given the well-documented poor efficacy of monotherapies for MPNST treatment, particularly *in vivo*, we performed combination testing of using 48 selected epigenetic compounds with Selumetinib in a 6×6 matrix format in 3KO NC and control fibroblasts spheroids (File S10). As single agents, neither Selumetinib, a clinically approved treatment for PNFs, nor Ribociclib (a CDK4/6 inhibitor, CDKi) exhibited a robust dose-response activity in the 3KO NC model (Figure S13) consistent with observations reported for other MPNST models.^27–30^ However, combinations of Selumetinib and BETi, HDACi, AURKA/Bi, or PARPi, indicated slight synergistic effects, as measure by analyses using both DBSumNeg and Excess Highest Single Agent (HSA) (Figure S14 A-B, File S10), consistent with previous results for some of these combinations.^13,31,32^

We next performed an in-depth assessment of combination effects with a reduced number of compound combinations across multiple cell lines, including the three isogenic 3KO lines (3KO_H8, 3KO_C6 and 3KO_D8) and three established MPNST lines (STS88-14, sNF96.2, NMS-2), thus covering models representing both an early stage MPNST development (3KO NC spheroids) and late stage (2D MPNST cell lines) (Figure S15A). We prioritized non-toxic combinations: PARPi (Olaparib, Talazoparib), the least toxic HDACi (Entinostat, Belinostat, ACY-1215) and BETi (I-BET151, GSK1324726A, Alobresib, CPI-0610) (Figure S15B).

While both HDACi and BETi combinations with Selumetinib reduced viability across all models (**Figure 8E i-ii**, S17A–B), PARPi effects were restricted to 3KO NC spheroids, with minimal impact on MPNST cell lines (**Figure 8E iii**, S16A-C). HDACi combinations with Selumetinib were generally toxic, narrowing the potential therapeutic window (Figure S16C, S17A) and BETi combos showed some toxicity at higher concentrations (Figure S16C, S17B). Dose-response curves were highly consistent across replicates, underscoring the reproducibility of the assays. To further assess potential synergistic effects with Selumetinib, combination index (CI) and fraction affected (Fa) values were calculated using three concentrations above and below the IC_50_ of the compounds in 3KO_H8. For both I-BET151 and Belinostat, most CI values were below 0.9, indicating synergy in both models (**Figure 8F i–ii**). In contrast, nearly none of the combinations of Selumetinib with PARP inhibitors exhibited clear synergistic effects in MPNST cell lines, with most CI values exceeding 1.5 (Figure 8F iii, S16B) but indeed reached CI values below 0.6-0.9 in 3KO NC spheroids.

### Pilot *in vivo* study using NF1 MPNST PDOX identifies Olaparib-based combinations as a potential therapeutic strategy for MPNSTs

Given the observed selectivity of PARPi-Selumetinib co-treatments across the 3KO NC *in vitro* models, we evaluated the efficacy of Olaparib in combination with Selumetinib, Ribociclib, or I-BET151, in an MPNST PDOX model (NF1-18B) described recently.^31,33^

A pilot exploratory *in vivo* treatment consisted of groups of 5-8 mice (52 animals), mixing females and males, that after PDOX formation, were treated for 21 days with distinct compounds and combinations. We used Olaparib in double and triple combinations with Selumetinib (MEKi), and Ribociclib (CDKi) or I-BET151 (BETi), that were compared to vehicle treated control, Olaparib as single agent and the triple MEKi-CDKi-BETi combination as the combination that elicited the greatest response in the NF1-18B PDOX model so far (**Figure 9A**).^31^ Treatment conditions are detailed in Figure S18A. Body weight was monitored daily to asses toxicity throughout the 21-day treatment period, with no weight loss detected (**Figure 9B**). Relative tumor volume for each mouse was also recorded (**Figure 9C**). At the end of the treatment, mice were euthanized, and tumors were photographed (Figure S18B) and weighed (**Figure 9D**) to compare treatment efficacy.

**Figure 9.**
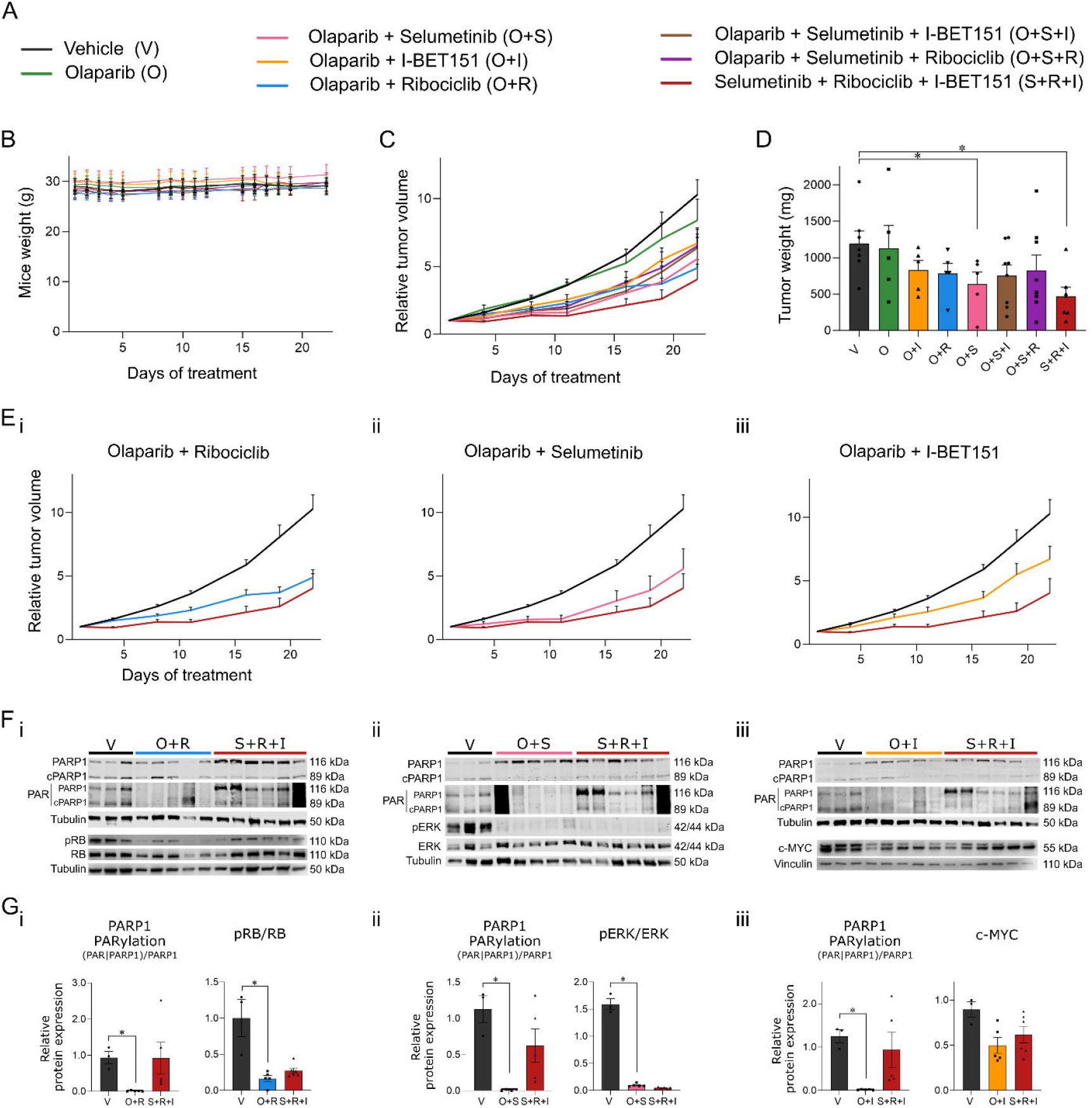
*In vivo* evaluation of Olaparib (PARPi) alone and in double or triple combinations with Selumetinib (MEKi) and Ribociclib (CDKi) or I-BET151 (BETi) in the NF1-associated NF1-18B MPNST PDOX mouse model. **A)** List of single, double, and triple treatment combinations used in the study, with each combination color-coded. **B)** Plot showing mice weight during the 3 weeks of treatment. **C)** Relative tumor volume growth of the NF1-18B PDOX mouse model in each treatment group over the course of the study. **D)** Tumor weight at the end of the experiment. Each black dot indicates one single value. Mann–Whitney test; *, p ≤ 0.05. **E)** Relative tumor volume growth for each double combination, vehicle control (negative control) and triple Selumetinib + Ribociclib + I-BET151 combination (positive control) throughout the experiment. (**i**) Olaparib + Ribociclib; (**ii**) Olaparib + Selumetinib; (**iii**) Olaparib + I-BET151. **F)** Western blot images of readouts of the molecular targets of Olaparib (PARP1 PARylation), Selumetinib (p-ERK/ERK), Ribociclib (p-RB/RB), and I-BET151 (c-MYC). (**i**) Olaparib + Ribociclib; (**ii**) Olaparib + Selumetinib; (**iii**) Olaparib + I-BET151. **G)** Quantification of molecular targets by Western blot for Olaparib (PARP1 PARylation), Selumetinib (p-ERK/ERK), Ribociclib (p-RB/RB and WEE1), and I-BET151 (MYC). (i) Olaparib + Ribociclib; (ii) Olaparib + Selumetinib; (**iii**) Olaparib + I-BET151. Mann– Whitney test; *, p ≤ 0.05.

Overall, single-agent Olaparib treatment did not change tumor volumes and growth kinetics compared to vehicle controls, indicating minimal anti-tumor activity. In contrast, all tested double combinations with Olaparib resulted in substantial reductions in relative tumor volume, as well as in tumor weight, compared with vehicle controls: Olaparib/I-BET151 achieved a 30.6% reduction, Olaparib/Ribociclib a 36.5% reduction, and Olaparib/Selumetinib a 61.6% reduction that was also statistically significant, suggesting an apparent therapeutic benefit (**Figure 9C-E**). Triple combinations of Olaparib, Selumetinib and a Ribociclib or I-BET151 did not enhance tumor reduction compared to double treatments (**Figure 9C**). Despite the limitations of this pilot *in vivo* study, Olaparib/Selumetinib combination yielded the most significant reduction in tumor weight (**Figure 9D**), close to the effect of the triple MEKi-CDKi-BETi combination, emerging as the most effective regimen from this work. However, as previously mentioned, both male and female mice were included in the study, and we identified different treatment responses according to mice sex (Figures S18C-E) which need to be explored in more detail.

Focusing on the favorable outcomes observed with the double-treatment regimens (**Figure 9E**) and comparing them with controls, we performed Western blot (WB) analyses in resected tumors at the end of the treatment, for each double combination to assess target engagement and downstream pathway effects (Rb, Myc and ERK) as described previously^31^ and PARP1 PARylation for PARPi, with loss of PARylation indicating target engagement.^34^ WB and their quantification (**Figure 9F and 9G**, respectively) confirmed the on-target activity of the different compounds within tumors, reaching statistical significance in all cases except for the reduction in c-MYC expression. In addition, WB analysis showed increased BIM levels in treated tumors, suggesting pro-apoptotic pathway activation, although without statistical significance (Figure S18F).

Taken together, these results indicate that our iPSC-derived NC models provide a valuable platform for drug screening with a good agreement with *in vivo* results and support the potential efficacy of of PARPi-MEKi or PARPi-CDKi, as a therapeutic option for MPNSTs.

## DISCUSSION

We studied the role of most commonly inactivated TSGs in the early development of MPNSTs using an iPSC-derived NC model system and sequentially inactivating *NF1, CDKN2A* and *SUZ12*, to mimic the benign-to-malignant transition during PNF-ANNUBP-MPNST progression.

It is now well established that the loss of *CDKN2A* in *NF1*(−/−) PNF cells is the trigger for ANNUBP formation,^6,7^ becoming a premalignant lesion difficult to manage in clinic due to the lack of tools for risk assessment. ^10,11,35^ In our NC models, 2KO NCs retain glial differentiation capacity and form neurofibroma-like lesions when engrafted in the sciatic nerve of nude mice. However, no histological atypia was detected in these neurofibromas, suggesting that a longer period of tumor development might be necessary. *CDKN2A* inactivation did not affect cell cycle or proliferation rate, but led to gene expression changes when compared to 1KO NCs. One of the differentially expressed genes was *TYMS* which is also differentially overexpressed in ANNUBPs compared to PNFs. TYMS inhibitors, like Pemetrexed, exist and have been used to inhibit proliferation in cells deficient in *CDKN2A*.^23^ Our results show the existence of a window of opportunity for *TYMS* inhibition in ANNUBPs, although further experiments are required to confirm this possibility.

*CDKN2A* encodes p14ARF and p16INK4a. Different genetically modified mouse (GEM) models have shown that the loss of *NF1* and *CDKN2A* is required for ANNUBP formation,^16,17^ some pointing to p14ARF as the key *CDKN2A* gene. Our results demonstrate that wild-type p14ARF function impeded the proper formation of tumors *in vivo*, but also support the necessity of losing both genes for ANNUBP development as it is seen in MPNST cell lines and human MPNST.^36,24^

PRC2 function helps maintain cell identity through transcriptional repression driven by closed chromatin ^37^ and has been involved in many types of cancer.^12^ Our work supports that the order of TSG inactivation in the PNF-ANNUBP-MPNST progression is biologically constrained. That would explain why neurofibromas in *NF1* patients carrying a constitutional microdeletion involving *NF1* and *SUZ12*, never exhibit LOH^38^: PRC2 loss would activate p14ARF/p16INK4a expression through chromatin accessibility, inducing senescence.^39,40^

We demonstrate that the NC models (1KO, 2KO, 3KO) developed here capture a benign-to-malignant transition, and a change in cell identity caused by PRC2 inactivation. PRC2 loss causes a global chromatin remodeling largely affecting genome methylation and transcription, in a programmed and reproducible manner, since the three independent 3KO NCs exhibit almost identical methylomes and transcriptomes. Mimicking the glial-to-mesenchymal transition observed from ANNUBP towards MPNST, 3KO NCs acquired a mesenchymal identity, consistent with previous results.^41^ Previous studies have shown the role of PRC2 in the acquisition of mesenchymal characteristics, by comparing MPNSTs with proficient and deficient PRC2, or re-expressing or deleting PRC2 components in MPNST or immortalized SC cell lines.^42,43,44^ Here we instead start from iPSCs to model stepwise TSG loss in NCs, a cell type close to the PNF/MPNST cell of origin, to investigate early tumorigenesis. We show that PRC2 loss changes cell identity by precluding gliogenesis capacity of 3KO NCs and activating the expression neuro-mesenchymal programs.

Gliogenesis is impeded by the epigenetic silencing of *SOX10*, a master regulator TF,^45^ upon closing chromatin accessibility. This supports previous findings in MPNSTs were *SOX10* was found downregulated and epigenetically silenced when compared to benign tumors, and downregulated after PRC2 inactivation in MPNST cell lines.^42,44,41,46^ On the other hand, absence of PRC2 triggers the expression of repressed neuro-mesenchymal gene programs by the opening of chromatin accessibility. 3KO NC express mesenchymal and stemness genes like *SOX9, TWIST1* and *PRRX1* and neurogenesis-associated transcriptional programs consistent with previous observations.^8,42,43, 47^ The combined analyses of expression and chromatin accessibility, comparing 2KO with 3KO NCs in one hand, and 3KO NCs with MPNST cell lines (and MSCs) on the other, allowed the identification and characterization of these neuro-mesenchymal signatures. We identified PRC2 poised neurogenesis genes that are immediately expressed after PRC2 loss, likely due to loss of REST–PRC2 repression,^48^ remaining active in MPNST cells. Other mesenchymal and neurogenesis genes may adopt an open chromatin conformation, or already have it, but 3KO NCs lack other necessary factors for their expression, some mesenchymal-associated present in MSCs and MPNST cell lines, and other unknown and only present in MPNSTs (perhaps associated to genomic alterations), driving the expression of additional neurogenesis genes. These identified gliogenesis, neurogenesis and mesenchymal signatures, reproduced the glial-to-neuro-mesenchymal transition when analyzed in human tumors representing the PNF-ANNUBP-MPNST malignant progression.

The reproduction of this cell identity transition during malignant progression, together with the capacity of 3KO NC spheres of generating histologically compatible early MPNST-like tumors in mice upon engraftment, reinforces the validity of these cells as a genuine MPNST model. This is the first *in vitro*/*in vivo* genetically engineered MPNST model incorporating loss of all three TSGs implicated in PNF-to-MPNST progression. Despite being diploid, the resulting tumors are hypercellular and proliferative, but lack the aggressiveness of MPNST PDOX models,^33^ consistent with an early developmental stage.

We then set up the 3KO NC MPNST spheroid model to be highly scalable, rapid, and reproducible, providing a robust platform for high-throughput drug screening (HTS). Because PRC2 loss has a great impact on chromatin, we tested a large library of compounds targeting epigenetic modifiers, identifying those inhibiting BET, HDAC and AURKA/B as active agents, which had already been identified as potential treatments for MPNST.^13,20,33,49–51^ However, most compounds inhibiting these targets were not selective for 3KO NCs. On the other hand, poly(ADP-ribose) polymerase inhibitors (PARPi) exhibited a preferential activity against PRC2-deficient spheroids, even though they were not as effective against MPNST cell lines in 2D. The combination Olaparib-selumetinib was synergistic in 3KO spheroid models and was well tolerated and significantly inhibited tumor growth *in vivo* in a patient-derived orthotopic xenograft MPNST mouse model. This result would have remained undetected if the HTS would have been done using MPNST cell lines in usual 2D cell growth assays. Further *in vivo* studies are needed, to better characterize the beneficial effects of this co-treatment, including optimizing MEKi and PARPi dosing regimens.

Several studies have explored potential combinations of PARPi with MEKi, CDKi, and BETi in different tumor types.^52–56^ In fact, PARPi have already been proposed as a therapeutic agent for MPNSTs^57^ although this line of research has been scarcely followed, only by Larsson and colleagues (2023).^58^ Here we uncover the combination of PARPi and Selumetinib as a therapeutic option for MPNST.

The NC spheroid system provides a representative model of PNF–ANNUBP–MPNST progression to investigate MPNST initiation mechanisms and malignant properties. It also serves as a robust platform for compound screening, bridging *in vitro* studies with *in vivo* validation and enabling the identification of novel combinatorial strategies for MPNST therapy.

Acknowledgments

We thank the IGTP core facilities and their staff for their contribution and technical support: Translational Genomics Core Facility; Cryobiology, High Performance Computing; Flow Cytometry (Gerard Requena, Joan Puñet Ortiz and Marco A. Fernández); IGTP-Hospital Universitari Germans Trias i Pujol (HUGTP) Biobank Facility (Laia Pérez-Roca, Christel Kisser Pioch, Reyes Rodríguez and Lidia Nieto) and Pathology Department (Dr. Pedro Luis González). We thank the Barcelona Stem Cell Bank, Regenerative Medicine Program, IDIBELL, Barcelona, for providing control iPSC line. We also thank the Compound Management team and the Bioprinting Lab at NCATS. We also thank the Endocrine Tumor research group led by Dr. Mireia Jordà (IGTP), and particularly Jennifer Marcos and Anna Rueda, for kindly providing lentiviral plasmids and assisting us with the transduction procedures. We are also grateful to the Fundación Proyecto Neurofibromatosis, the Asociación de Afectados de Neurofibromatosis (AANF), and the Catalan Neurofibromatosis Association (ACNefi) for their constant support. This work has mainly been supported by la Fundació La Marató de TV3 (51/C/2019) and Children’s Tumor Foundation, Drug Discovery Initiative (Grant ID: CTF-2023-05-001). The work has also been partially supported by the Spanish Ministry of Science and Innovation, Carlos III Health Institute (ISCIII) (PI20/00228, PI23/00422 and PI23/00583) Plan Estatal de I+D+I 2013–2016, co-financed by the FEDER program – a way to build Europe, by the Government of Catalonia and CERCA Program/Generalitat de Catalunya (2021 SGR 00967), and Fundación Proyecto Neurofibromatosis. Work at NCATS was funded by the NIH Intramural Research Program (IRP). The contributions of the NIH author(s) are made as part of their official duties as NIH federal employees are in compliance with agency policy requirements and are considered Works of the United States Government. However, the findings and conclusions presented in this paper are those of the author(s) and do not necessarily reflect the views of the NIH or the U.S. Department of Health and Human Services. Single cell data generated was supported by Department of Defense office of the Congressionally Directed Medical Research Programs (CDMRP) NFRP FY20 (NF200051). IU-A is supported by a PFIS fellowship from the ISCIII (FI21/00063).

## Materials and Methods

### Human iPSC lines

The following human induced pluripotent stem cell (hiPSC) lines were utilized in this study: FiPS Ctrl1-SV4F-7 used as a control *NF1*(+/+) cell line and named WT in this article, and *NF1*(−/−) edited FiPS Ctrl1-SV4F-7 previously generated in our lab^21^ and named 1KO_D12 in this article. These lines are banked and stored at the Spanish National Stem Cell Bank, Institute of Health Carlos III (BNLC-ISCIII). Their use was approved by the local ethics committee (Clinical Research Ethics Committee of Germans Trias i Pujol Hospital, Badalona, Spain, Project No PI-13-021).

The iPSC lines used were authenticated by PCR-based DNA fingerprinting analysis using the AmpFlSTR_IdentifilerTM PCR Amplification Kit panel from Applied Biosystems.

### hiPSC culture

Human iPSCs were cultured on Growth Factor Reduced Matrigel (Corning) diluted at a 1:20 ratio with mTeSR media (StemCell Technologies), and maintained at 37°C in a 5% CO_2_ atmosphere. Passage of cells, when necessary, was performed using Accutase (Thermo Fischer Scientific).

### CRISPR/Cas9 gene edition

We generated *NF1*(−/−) *CDKN2A* p14ARF(+/+)p16INK4a(−/−) iPSC lines (referred to as 2KO (p16)) and *NF1*(−/−) *CDKN2A* p14ARF(−/−)p16INK4Aa(−/−) iPSC lines (referred to as 2KO (p14p16)) through CRISPR/Cas9 gene editing of the *NF1*(−/−) edited FiPS Ctrl1-SV4F-7 (referred to as 1KO cell line) using the ArciTect ribonucleoprotein (RNP) system from StemCell Technologies. We next edited SUZ12 in the 2KO_F9 iPSC line, generating *NF1*(−/−) *CDKN2A* p14ARF(−/−)p16INK4a(−/−) SUZ12(−/−) cells (referred to as 3KO). We also edited the TP53 gene in 3KO NC cells (3KO_H8 cell line), generating a *NF1*(−/−) *CDKN2A* (−/−) *SUZ12* (−/−) TP53 (+/−) NC line (referred to as 3KO+).

Briefly, sgRNAs targeting exon 2 of the *CDKN2A* gene for 2KO (p16) and for 2KO (p14P16)), exon 1 and exon 10 of the *SUZ12* gene were introduced into hiPSCs (File S1). In the case of the *TP53* gene, sgRNAs targeting exon 5 were introduced into NCs (File S1). In all cases, sgRNAs were introduced using TransIT-X2 transfection reagent (Mirus), following manufacturer’s instructions. Single-cell clones were, amplified, and DNA was extracted for *CDKN2A* and *SUZ12* and *TP53* mutation analysis using DNA Sanger sequencing. For the sequencing of each allele, the Gateway Technology cloning method (Invitrogen) was employed using the pDONR 221 vector. Sequences were analyzed using CLC workbench 8 software (Qiagen). Primers used for sequencing can be found in File 1.

#### Other cell lines

Three MPNST cell lines were used: sNF96.2 (RRID: CVCL_K281), ST88-14 (RRID: CVCL_8916), and NMS-2 (RRID: CVCL_4662). The human NF1 patient-derived fibroblast cell line (Coriell #GM00622) was used as a control cell line.

ST8814, sNF96.2 and NF1 fibroblasts were maintained in DMEM + 10% FBS + 2mM GlutaMAX with and 500 U/mL penicillin/500 mg/mL streptomycin (Gibco). NMS-2 cell line was maintained in RPMI + 10% FBS + 2mM GlutaMAX with and 500 U/mL penicillin/500 mg/mL streptomycin (Gibco). All cell lines were incubated at 37°C with 5% CO_2_.

### iPSC differentiation toward neural crest (NC)

Neural crest (NC) differentiation was performed as reported before.^25^ The NC medium consisted of: DMEM:F12 (Gibco) 1:1; 5 mg/mL BSA (Sigma); 500 U/mL penicillin/500 mg/mL streptomycin (Gibco); 2 mM GlutaMAX (Gibco); 1x MEM non-essential amino acids (Gibco); 1x trace elements A (Corning); 1x trace elements B (Corning); 1x trace elements C (Corning); 2-mercaptoethanol (Gibco); 10 mg/mL transferrin (Sigma); 50 mg/mL sodium L-ascorbate (Sigma); 10 ng/mL Heregulin-b1 (PeproTech); 200 mg/mL LONG R3 IGFR (PeproTech); 8 ng/mL basic fibroblast growth factor 2 (PeproTech), 2 mM CHIR9902 (STEMCELL Technologies) and 20 mM SB432542 (STEMCELL

Technologies). NCs were maintained in this medium and split with Accutase (Thermo Fischer Scientific) when necessary.

### 3D spheroid generation

For engraftment experiments, neurofibromasphere generation of 1KO and 2KO hiPSC cells was performed as reported before.^21,22^ For 3KO cells, 1.2 x 10^6^ cells were seeded into AggreWell™ 800 plates pre-treated with Anti-Adherence Rinsing Solution (STEMCELL Technology) in NC media. 48 hours later, spheroids were collected and engrafted into the sciatic nerve of nude mice.

For HTS experiments, three thousand neural crest (NC) cells or two thousand control *NF1*(+/−) fibroblast cells were seeded per well in 384-well, round-bottom, ultra-low attachment (ULA) plates (Corning, cat. no. CLS3830), in either NC medium or fibroblast medium, respectively. Following seeding, plates were centrifuged at 300 g for 5 minutes to promote cell aggregation and incubated at 37°C in a humidified atmosphere with 5% CO_2_. Under these conditions, cells spontaneously formed spheroids within 24 hours. Spheroids were treated with compounds at 24 hours post-seeding, and downstream functional assays were performed at 48 hours.

### Animal models

All mouse experiments were conducted in accordance with relevant Spanish national laws (RD 1201-/2005) and international guidelines and regulations (European Union rules 2010/63/UE), about the protection of animals used for experimentation and other scientific purposes. The experimental protocols were approved by the IDIBELL Animal Ethic Experimentation Committee (CEEA-IDIBELL approval No 9111) in Spain. They complied with the International Association for Assessment and Accreditation of Laboratory Animal Care procedures (AAALAC). Six-week-old male and female nude mice (Harlan) were used and maintained under sterile air flow isolator conditions, artificial light-dark cycle of 12 hours, regulated temperature of 22°C, relative humidity of 55%, and fed with autoclaved standard food and acidified water *ad libitum*.

### Engraftment experiments

For 1KO and 2KO cell lines, neurofibromaspheres were engrafted following previously reported protocols.^21,22^

For 3KO cell lines, spheroids containing approximately 2 million cells were resuspended in 1:2 diluted Growth Factor Reduced Matrigel (Corning) in a total volume of 70µl of NC medium and injected into the exposed sciatic nerve of nude mice using a 25 G syringe. We injected 1KO, 2KO_C4 and 2KO_F9 cell lines and the three 3KO cell lines (3KO_C6, 3KO_H8 and 3KO_D8). Tumor growth was monitored by manual palpation, and after 4 months, mice were euthanized and tumors fixed in formalin for paraffin embedding.

### Western Blot

Cells were washed twice with chilled PBS and lysed with RIPA buffer (50 mM Tris-HCl (pH 7.4), 150 mM NaCl (Merck), 1mM EDTA (Sigma), 0.5% Igepal CA-630 (Sigma)) supplemented with 3mM DTT (Thermo Fisher), 1mM PMSF (Sigma), 1mM sodium orthovanadate (Sigma), 5mM NaF (Sigma), 10 µg/ml leupeptin (Sigma), 5µg/ml aprotinin (Sigma) and 1xPhosSTOP (Roche).

Protein was extracted from vehicle control tumors (n=3), tumors treated with the double combinations (n=5) and triple control tumors (n=6) using RIPA buffer supplemented with protease inhibitor (complete Tablets, Roche). Briefly, 15–20 mg of tissue was homogenized in RIPA using TissueLyser II (Qiagen) and centrifuged at 16,000 × g at 4°C for 10 minutes.

Proteins were quantified with the Pierce BCA Protein Assay Kit (ThermoScientific). Lysates were boiled with 1X Laemmli buffer (50% glycerol (v/v) (Sigma), 10% SDS (m/v) (Merck), 0.05% bromophenol blue (m/v) (Sigma), 25% Tris HCl 1M pH 6.8 (v/v) (Millipore), 5% beta-mercaptoethanol (v/v) (Sigma) in distilled water) and 20 µg of protein was used for Western blot detection using either chemiluminescence or fluorescence. For chemiluminescence detection, protein was separated on 12% SDS-acrylamide gels (Bio-Rad) and transferred to nitrocellulose membranes. Membranes were blocked with BSA, incubated with primary antibodies (Table S1) overnight at 4°C and detected with SuperSignal West Femto chemiluminescent substrate kits (Thermo Fisher Scientific). Quantification was done using Image Lab, normalizing proteins to tubulin or vinculin. For fluorescence detection, protein was subjected SDS-PAGE and transferred onto PVDF membranes (1 hour 350 mA at 4°C; or 18 hours 90 mA at 4°C, for neurofibromin). Membranes were blocked with Odyssey Blocking Buffer (TBS, LI-COR) and incubated with primary antibodies (Table S1) at 4°C overnight; and with mouse anti-αtubulin (Sigma) 1 hour at room temperature. Membranes were then incubated with IRDye 680RD anti-Rabbit and IRDye 800CW anti-Mouse secondary antibodies (1:10,000 each, LI-COR) for 1 hour at room temperature and scanned with the Odyssey CLx (LI-COR) using the Image Studio Lite (LI-COR).

### Flow cytometry

Cells were dissociated with Accutase and resuspended in 0.1% BSA in PBS. For NC linage markers, cells were incubated for 30 min on ice with unconjugated p75 primary antibody and detected with Alexa Fluor 568-conjugated secondary antibodies, following incubation for 30 min on ice with unconjugated primary antibody Hnk1 and detected with Alexa Fluor 488-conjugated secondary antibodies. For MSC markers, cells were incubated with conjugated APC-CD73, PE-CD13 and PE-CD44 for 15min. Cells were analyzed by flow cytometry using BD FASCCanto II and analyzed using BD FACSDiva 6.2 software. See Table S1 for antibody details.

### Immunocytochemical analysis

Cells were fixed in 4% paraformaldehyde (PFA) (Chem Cruz) in PBS for 15 minutes, permeabilized with 0.1%Triton-X100 in PBS for 10 minutes, blocked in 10% FBS in PBS for 15 minutes, and incubated with primary antibodies (OCT3/4, TFAP2, SOX10, SOX9, S100B, p75 and H3K27me3) overnight at 4°C. Secondary antibodies were Alexa Fluor 488 and Alexa Fluor 568 (Thermo Fisher Scientific). Nuclei were stained with DAPI (Stem Cell Technologies, 1:1000). Slides were mounted with Vectashield (Vector laboratories), and coverslips were secured with a polish nail. See Table S1 for antibody details.

Spheroids were fixed in 4% PFA (Chem Cruz) in PBS for 30 minutes at room temperature. The staining protocol was the same as for cells.

Fluorescent and phase contrast images from cells and spheroids were captured using the DMI 6000B microscope (Leica) and LAS X software (Leica).

### Quantitative High-Throughput Screen (qHTS)

A targeted chemical screen was performed using the NCATS Pharmacologically Active Chemical Toolbox (NPACT) Epigenetics qHTS Library containing 339 small molecules (File S7) in 3D spheroid cultures of NC-derived cells (1KO, 2KO, 3KO, and 3KO+ lines) and control fibroblast spheroids (GM00622). Compounds were prepared as DMSO solutions and pre-plated in two 1536-well source plates at threefold serial dilution.

At 24 hours cell post-seeding, compounds were transferred to spheroid assay plates using a 384-pin tool dispenser, and cells were incubated with compounds for 48 hours at 37°C in a humidified 5% CO_2_ incubator. Cell viability was assessed using the CellTiter-Glo® 3D Cell Viability Assay (Promega) following manufacturer’s instructions.

### Curve Response Class (CRC)

Compound activity was determined based on their curve response class (CRC), in which normalized data is fitted to 4-parameter dose-response curves using a custom grid-based algorithm to generate a curve response class score for each compound dose-response.^59^ CRC values of −1.1. −1.2, −2.1, are considered the highest quality hits; CRC values of −1.3, −1.4, −2.3, −2.2, −2.4, and −3 are considered inconclusive hits; and positive CRC values are considered inactive compounds.^26^

### *In vitro* validation in additional 3KO cell lines and MPNST cell lines

384-well plates were pre-spotted at NCATS with candidate compounds at the following starting concentrations: PARP inhibitor Olaparib (20⍰µM) and Talazoparib (1⍰µM); HDAC inhibitors Entinostat (10⍰µM), Belinostat (10⍰µM), and ACY-1215 (20⍰µM); and BET inhibitors I-BET151 (10l1µM), GSK1324726A (2⍰µM), Alobresib (2⍰µM), and CPI-0610 (20⍰µM). Each compound was pre-spotted as 8-point, 1:2 serial dilution, alone or in combination with Selumetinib at 20⍰µM, 5⍰µM, or 1.25⍰µM (n=2).

Three different 3KO cell lines (3KO_H8, 3KO_C6, and 3KO_D8) and the control fibroblast line GM00622 were cultured as spheroids (3D). MPNST cell lines (sNF96.2, ST88-14, and NMS-2) were maintained in 2D. Compounds were manually resuspended and pipet dispensed (10⍰µL/well) in the spheroid plates. The 3D viability assay was performed as previously described. For 2D cultures, viability was assessed using the CellTiter-Glo® Luminescent Cell Viability Assay (Promega) following manufacturer’s instructions. Luminescence was recorded using GloMax Explorer plate reader (Promega). Synergy was determined using CompuSyn software (RRID: SCR_022931), based on Chou–Talalay calculations.^60^

### Drugs and Maximum tolerated dose test

Drugs for in vivo experiments were purchased from MedChemExpress: Olaparib (Cat. #HY-10162), Selumetinib (Cat. #HY-50706), I-BET151 (Cat. #HY-13235), and Ribociclib (Cat. #HY-15777). We adjusted the dosages and the administration schemes for Olaparib and combinations, using athymic nude six-week-old male and female mice, in which no tumor was engrafted. Three mice were used for each triple combination condition (Olaparib + Selumetinib + Ribociclib; Olaparib + Selumetinib + I-BET151). The doses of Selumetinib, Ribociclib, and I-BET151 were selected based on previous studies.^31^ Mice were treated for three weeks and body weight was daily measured to analyze potential toxicity. The optimal doses of each compound, at which no toxicity was observed, were determined to be: Olaparib 40 mg/kg, Selumetinib 45 mg/kg, I-BET151 22 mg/kg, and Ribociclib 85 mg/kg.

### *In vivo* drug testing on the MPNST PDOX model

The NF1-18B PDOX model established in Conxi Lazaro’s Lab was used.^31,33^ Tumors were expanded in 5-week-old athymic nude mice. When tumors reached 1,000 to 1,500 mm^3^, they were grafted into the sciatic nerve of new 5-week-old mice. Each group consisted of 5 to 8 mice. Once tumors reached 300 to 500 mm^3^, mice were randomized into treatment groups and treated for 3 weeks. In the absence of observed toxicity, doses of several compounds were escalated after the first two weeks, resulting in the following regimen: Olaparib 40 mg/kg orally once daily for the first 2 weeks, then 45 mg/kg once daily; Selumetinib 45 mg/kg orally once daily for the first 2 weeks, then 55 mg/kg once daily; I-BET151 22 mg/kg intraperitoneally three days per week; and Ribociclib 85 mg/kg orally once daily. Mice were sacrificed 4 to 5 hours after the last dose, and tumors were removed. Tumors were measured using a caliper twice a week, and the volume was calculated using the formula v = (w^2^L/2), in which L is the longest diameter and w is the width. Statistical analyses used the Mann– Whitney test, with a significance level at 0.05.

### RNA-seq and analysis

RNAseq library was prepared at BGI (Shenzhen, China) using DNBseq standard protocols. Data was aligned with Salmon v1.8.0 (RRID:SCR_017036)^61^ against the UCSC refMrna and hg38 genome. Transcript-level estimates were imported into R (R v4.3.0 Bioconductor v3.17) and summarized to the gene level using tximport (RRID:SCR_016752). Genes with less than 5 counts in more than 2 samples were filtered out. A principal component analysis (PCA) plot was generated. Differentially expressed genes (DEG) between different cell types were identified using DESeq2 (RRID:SCR_015687)^62^ with the Wald test and apglm (RRID:SCR_015687). Genes with an adjusted p-value < 0.05 were considered differentially expressed. Heatmaps were created using the pheatmap package. Finally, we used the clusterProfiler R package^63^ for DEG enrichment analysis.

### ATACseq

ATACseq was produced at Diagenode (Ougrée, Belgium). In brief, ATAC-seq reads were mapped to the GRCh38 human reference genome using BWA-MEM ^64^ and deduplicated Picard MarkDuplicates. We then called peaks with macs2^65^ with standard parameters and used deeptools bamCoverage to generate bigwig files for plotting. For differential accessibility, we called consensus peaks with macs2, converted the narrowPeak files into SAF with awk and used featureCounts to create a counts table. Consensus peak counts were imported into R. edgeR^66^ was used for normalization and differential analysis between genotypes (1KO and 2KO vs 3KO). Peaks were assigned to genes and a score was calculated for each peak based on p value, log FC and logCPM. The peak with the greatest score was selected to represent each gene in volcano plots and in plots correlating accessibility and expression. Functional enrichment was performed on GO terms with clusterProfiler^63^ with genes with a p value below 0.05 and a log fold change over +/−1 independently for differentially closed and differentially open genes.

### Data Visualization

ATAC-seq coverage plots were created from bigwig files with karyoploteR.^67^ Heatmaps were plotted with ComplexHeatmap.^68^ Boxplots of neurogenesis genes were plotted using R after standardizing the expression data to mean=0 and sd=1. Schemes and diagrams were created with Biorender. Low throughput molecular data was analyzed and plotted with GraphPad Prism (version 7.00).

### Additional material and methods

The following additional sections of materials and methos are available as supplementary:

DNA extraction; RNA extraction; MPNST cell lines and PNF-derived Schwann cells RNAseq data; Plexiform neurofibromas (PNFs), ANNUBPS, MPNSTs and nerve samples; Neural Crest (NC) differentiation towards Mesenchymal Stem Cells (MSC); Neural Crest (NC) differentiation towards Schwann cells (SCs); Immunohistochemical analysis of formalin-fixed paraffin-embedded (FFPE) samples; Fluorescent immunohistochemistry of FFPE samples; Senescence assay; Ploidy assay; Proliferation assay; Lentiviral vectors; Viral production and viral transduction; Pemetrexed treatment; Extended qHTS; Live/dead cell viability assay; Compound Combination Matrix Screening; DNA methylation; Single-cell RNA-Seq and analysis; WGS.

## Author contributions

Conceptualization, E.S., M.C., I.U.-A., M.M.-L. and B.G.; methodology, I.U.-A., M.C., J.F.-R., J.Z. and E.L.; software and formal analysis, M.M.-L. and B.G.; investigation, I.U.-A., J.F.-R., M.M.-L., J.Z., E.L., B.G., H.M., E.C.-B, J.F.-C., S.O.-B., K.M.W., C.M., K.R., C.R., M.C. and E.S.; resources, C.L., M.F., E.S.; writing – original draft, I.U.-A., M.C., and E.S.; writing – review & editing, M.M.-L., B.G., I.U.-A, J.Z., J.F.-C, M.F., M.C., and E.S.; visualization, I.U.-A, M.M.-L., B.G., M.C., and E.S.; supervision, B.G., M.F., M.C., and E.S.; project administration, M.C. and E.S.; funding acquisition, M.F. and E.S.

## Declaration of generative AI and AI-assisted technologies in the writing process

During the preparation of this work, authors used ChatGPT to rephrase or shorten the text. After using this tool, authors reviewed and edited the content as needed and take full responsibility for the content of the publication.

## Data and code availability

Omic data generated or used in this work has been deposited and is available from different sources:

**RNA-seq data:** iPSC WT, 1KO,^21^ deposited in EGA (https://ega-archive.org/), accession number EGA:EGAS00001005907; iPSC 2KO, 3KO; NC WT, 1KO, 2KO, 3KO; MSC WT, 1KO, 2KO, 3KO; MPNST cell lines (this publication), deposited in NF Data Portal (https://doi.org/10.7303/syn68876148); PNF-SC cultures,^21^ deposited at EGA (https://ega-archive.org/), accession number EGA: EGAS00001005907.

**Methylome data:** MSC WT, 1KO, 2KO, 3KO; NC WT, 1KO, 2KO, 3KO (this publication), MPNST cell lines,^36^ deposited in NF Data Portal (https://doi.org/10.7303/syn68876148).

**ATAC-seq data:** NC WT, 1KO, 2KO, 3KO (this publication), deposited in NF Data Portal (https://doi.org/10.7303/syn68876148)

**scRNA-seq data:** nerves, PNFs, ANNUBPs, MPNSTs (Gel et al. manuscript in preparation), deposited in NF Data Portal: DOI: https://doi.org/10.7303/syn52146919

**WGS:** iPSC WT, 1KO, 2KO, 3KO (this publication), deposited in NF Data Portal (https://doi.org/10.7303/syn68876148)

## Supplemental information index

**Document S1**. Figure S1-S18; Table S1-S3; Supplementary materials

**File S1**. Excel file containing the sgRNA sequences, the list of mutations in gene-edited cell lines and the sequencing and cloning primers. Related to Figure 1 and S3.

**File S2**. Excel file containing the list of genes in common between differentially overexpressed genes in 2KO (n=3) vs 1KO (n=3) and differentially expressed genes in ANNUBPs (n=3) vs. PNFs (n=3). Related to Figure 1.

**File S3**. Excel file containing the list of differentially expressed genes and enrichment of biological processes between 3KO and 2KO NC cell lines. Related to Figure 5 and Figure S5.

**File S4**. Excel file containing the list of differentially expressed genes and enrichment of biological processes between 2KO and 3KO NC cell lines. Related to Figure S6.

**File S5**. Excel file containing the list of differentially expressed genes and enrichment of biological processes between 3KO NC and MPNST cell lines. Related to Figure 5 and Figure S5.

**File S6**. Excel file containing the list of differentially open/closed genes and enrichment of biological processes between wild-type PRC2 and mutated PRC2 NC cell lines. Related to Figure 5.

**File S7**. Excel file containing NCATS Pharmacologically Active Chemical Toolbox (NPACT) epigenetics library.

**File S8**. Excel file containing qHTS results in 3D cell lines: isogenic cell lines (1KO, 2KO, 3KO, 3KO+) and fibroblast NF1 (+/−) controls. Related to Figure 8.

**File S9**. Excel file containing the secondary screening results for 77 compounds (n=2) in all 3D cell lines: isogenic cell lines (1KO, 2KO, 3KO, 3KO+) and fibroblast NF1 (+/−) controls. Related to Figure 8.

**File S10**. Excel file containing all the combination screening results. Related to Figure S13.

