## Supplementary_Information for "iPSC-derived *NF1*-*CDKN2A*-PRC2 deficient neural crest mimics MPNST glial-to-neuro-mesenchymal transition and uncover new therapeutic opportunities"

Figure S1

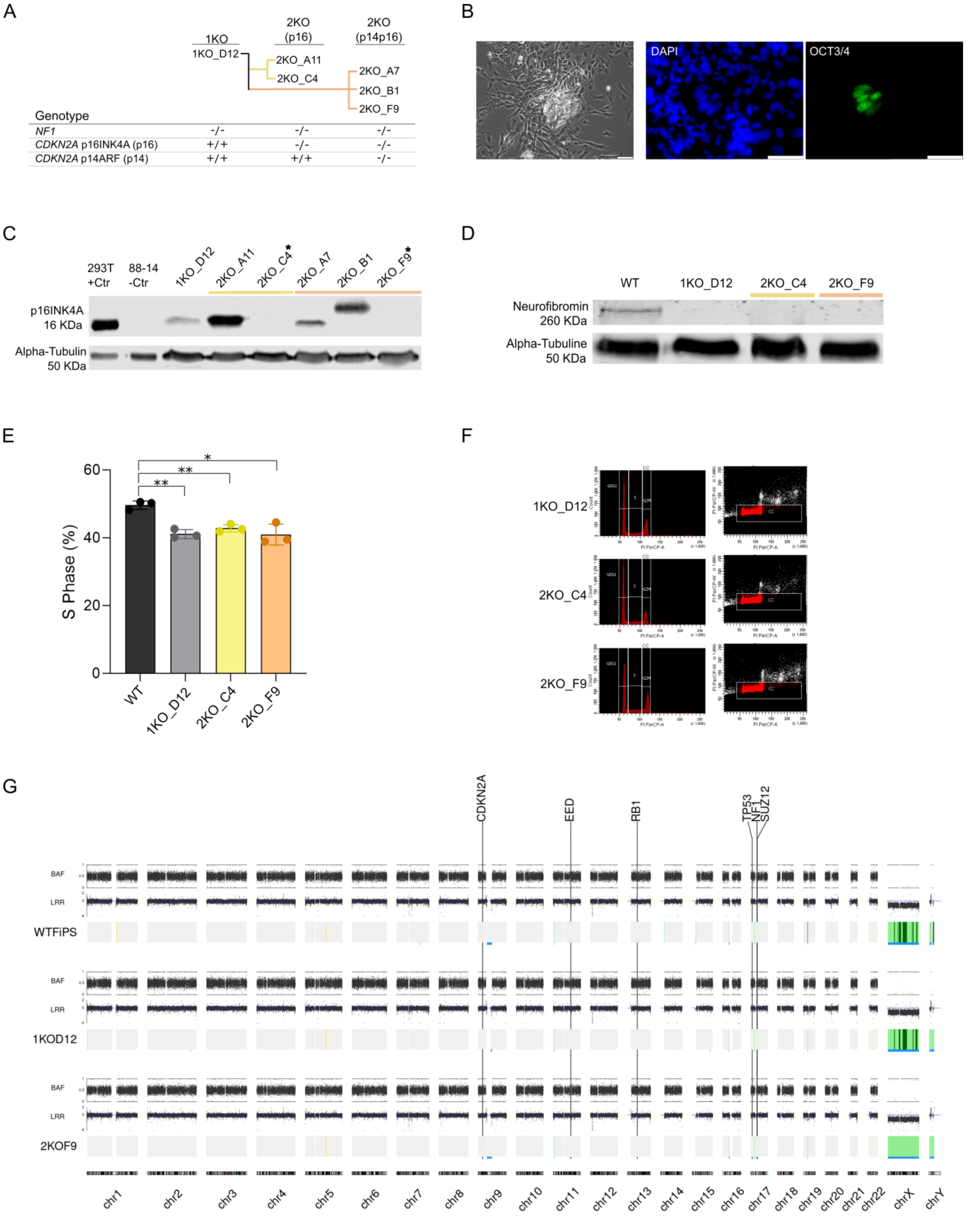
A
1KO
2KO (p16)
2KO (p14p16)
1KO_D12
2KO_A11
2KO_C4
2KO_A7
2KO_B1
2KO_F9
Genotype
NF1 -/- -/- -/-
CDKN2A p16INK4A (p16) +/+ -/- -/-
CDKN2A p14ARF (p14) +/+ +/+ -/-
B
DAPI
OCT3/4
C
293T +Ctr
88-14 -Ctr
1KO_D12
2KO_A11
2KO_C4*
2KO_A7
2KO_B1
2KO_F9*
p16INK4A 16 KDa
Alpha-Tubulin 50 KDa
D
WT
1KO_D12
2KO_C4
2KO_F9
Neurofibromin 260 KDa
Alpha-Tubuline 50 KDa
E
S Phase (%)
60
40
20
0
*
**
**
WT
1KO_D12
2KO_C4
2KO_F9
F
1KO_D12
2KO_C4
2KO_F9
G
CDKN2A
EED
RB1
TP53
NF1
SUZ12
BAF
LRR
WTFiPS
1KOD12
2KOF9
chr1
chr2
chr3
chr4
chr5
chr6
chr7
chr8
chr9
chr10
chr11
chr12
chr13
chr14
chr15
chr16
chr17
chr18
chr19
chr20
chr21
chr22
chrX
chrY

Figure S1

**Figure S1. Characterization of *CDKN2A* knock out *NF1*(-/-) iPSC lines, related to Figure 1. A)** Dendrogram of edited iPSC lines, showing their genotypes. See also File S1. **B)** Left: Representative micrograph illustrating spontaneous differentiation observed in the edited 2KO_A11 iPSC clone. Scale bar: 100 µm. Right: Immunofluorescence image demonstrating lack of OCT3/4 expression in 2KO_A11 iPSC clone. Scale bar: 50 µm. **C)** Western blot analysis for p16INK4a in edited cell lines at NC stage. HEK 293T cells were used as positive control; and 88-14 MPNST cell line as negative control. (*) Represent candidate clones selected for further analyses. **D)** Western blot analysis for neurofibromin in *NF1* (+/+) WT and *NF1* (-/-) 1KO and 2KO iPSC cell lines. **E)** Proliferation capacity of WT, 1KO and 2KO NC lines assessed by Click-iT EdU flow cytometry assay. Bars represent mean values ± SD from three independent experiments. Unpaired t tests: **, $p < 0.01$; *, $p < 0.05$. **F)** Flow cytometry analysis of ploidy in 1KO and 2KO NC cell lines assessed by propidium iodide staining. All three lines exhibit diploid genomes. Viable cells are shown in red. **G)** Genomic structure of WT iPSC (WTFIPS), 1KO_D12 iPSC and 2KO_F9 iPSC cell lines analyzed by whole genome sequencing. B-allele frequency (BAF) and log R ratio (LRR) plots indicate a diploid (2n) genome and absence of copy number alterations.

Figure S2

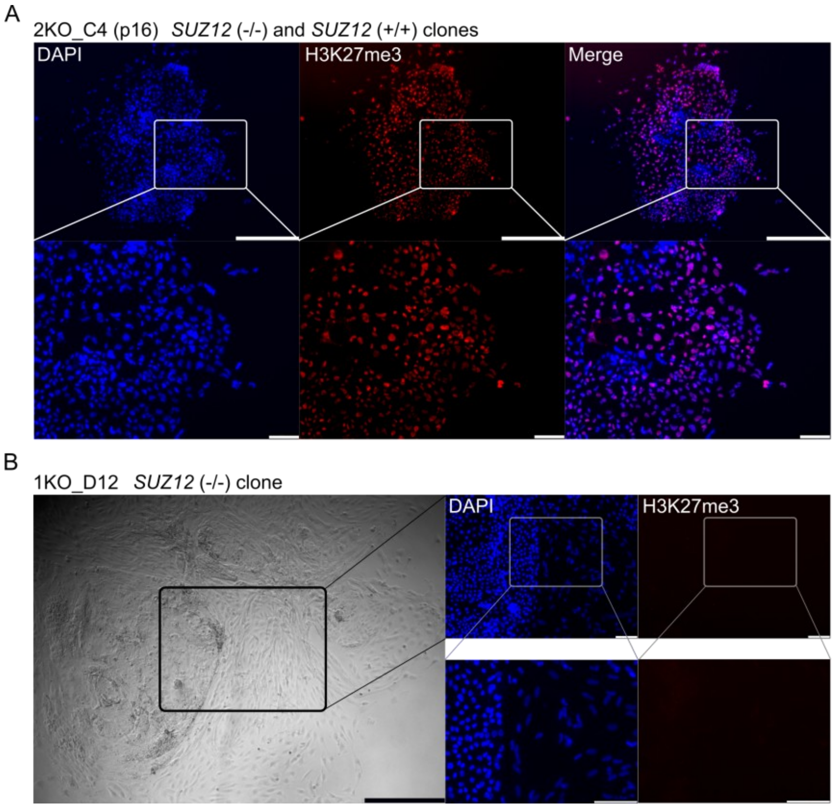


**Figure S2. H3K27me3 expression in *SUZ12*-edited cells, related to Figure 3. A)** Immunofluorescence detection of H3K27me3 in a 2KO (p16)-*SUZ12* mixed colony showing *SUZ12* WT (in red) and *SUZ12* knock out cells. Upper panel scale bar: 500 µm. Lower panel scale bar: 100 µm. **B)** Example of loss of pluripotency in 1KO iPSCs after *SUZ12* editing. Left panel: Phase-contrast image showing an iPSC colony surrounded by differentiated cells. Scale bar: 500 µm. Right panel: Immunofluorescence staining for H3K27me3 showing negative staining in differentiated areas confirming *SUZ12* edition. Scale bar: 100 µm.

Figure S3

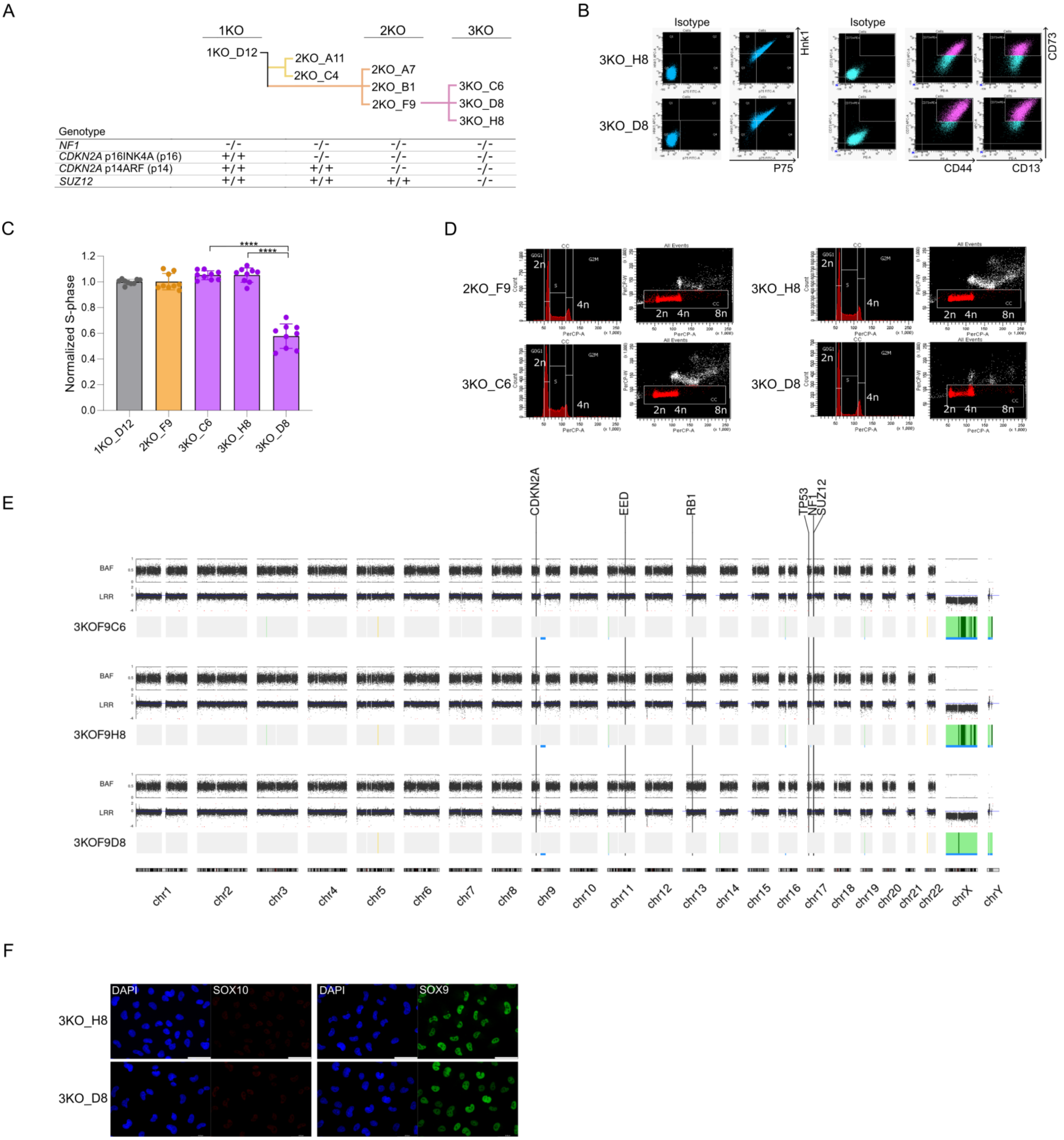
A
1KO
2KO
3KO
1KO_D12
2KO_A11
2KO_C4
2KO_A7
2KO_B1
2KO_F9
3KO_C6
3KO_D8
3KO_H8
Genotype
NF1 -/- -/- -/- -/-
CDKN2A p16INK4A (p16) +/+ -/- -/- -/-
CDKN2A p14ARF (p14) +/+ +/+ -/- -/-
SUZ12 +/+ +/+ +/+ -/-
B
Isotype
Isotype
3KO_H8
3KO_D8
Hnk1
P75
CD73
CD44
CD13
C
Normalized S-phase
****
****
1KO_D12
2KO_F9
3KO_C6
3KO_H8
3KO_D8
D
2KO_F9
3KO_C6
3KO_H8
3KO_D8
2n
4n
8n
E
CDKN2A
EED
RB1
TP53
NF1
SUZ12
BAF
LRR
3KOF9C6
3KOF9H8
3KOF9D8
chr1
chr2
chr3
chr4
chr5
chr6
chr7
chr8
chr9
chr10
chr11
chr12
chr13
chr14
chr15
chr16
chr17
chr18
chr19
chr20
chr21
chr22
chrX
chrY
F
DAPI
SOX10
DAPI
SOX9
3KO_H8
3KO_D8

Figure S3

**Figure S3. Characterization of *SUZ12*-edited iPSC-derived NC cell lines, related to Figure 3. A)** Schematic editing tree illustrating the stepwise generation of isogenic cell lines. The table below details the genotype of each cell line for *NF1*, *CDKN2A*, and *SUZ12*. See also File S1. **B)** Flow cytometry analysis of NC markers (p75 and Hink1) and MSC markers (CD73, CD44 and CD13) in 3KO_H8 and 3KO_D8 NC lines. **C)** Proliferation capacity of 1KO, 2KO and 3KO NC lines assessed by Click-iT EdU flow cytometry assay. 3KO_D8 exhibited a significantly reduced proliferative capacity. Bars represent mean values ± SD from three independent experiments. Unpaired t tests: ****, $p < 0.0001$. **D)** Flow cytometry analysis of ploidy in 2KO and 3KO NC cell lines assessed by propidium iodide staining. All lines exhibit a diploid genome. Viable cells are shown in red. **E)** Genomic structure of 3KO cell lines analyzed by whole genome sequencing. B-allele frequency (BAF) and log R ratio (LRR) plots indicate a diploid (2n) genome and absence of copy number alterations**. F)** Immunofluorescence images for SOX10 (red) and SOX9 (green) in 3KO NC lines. Scale bar: 50 µm.

Figure S4

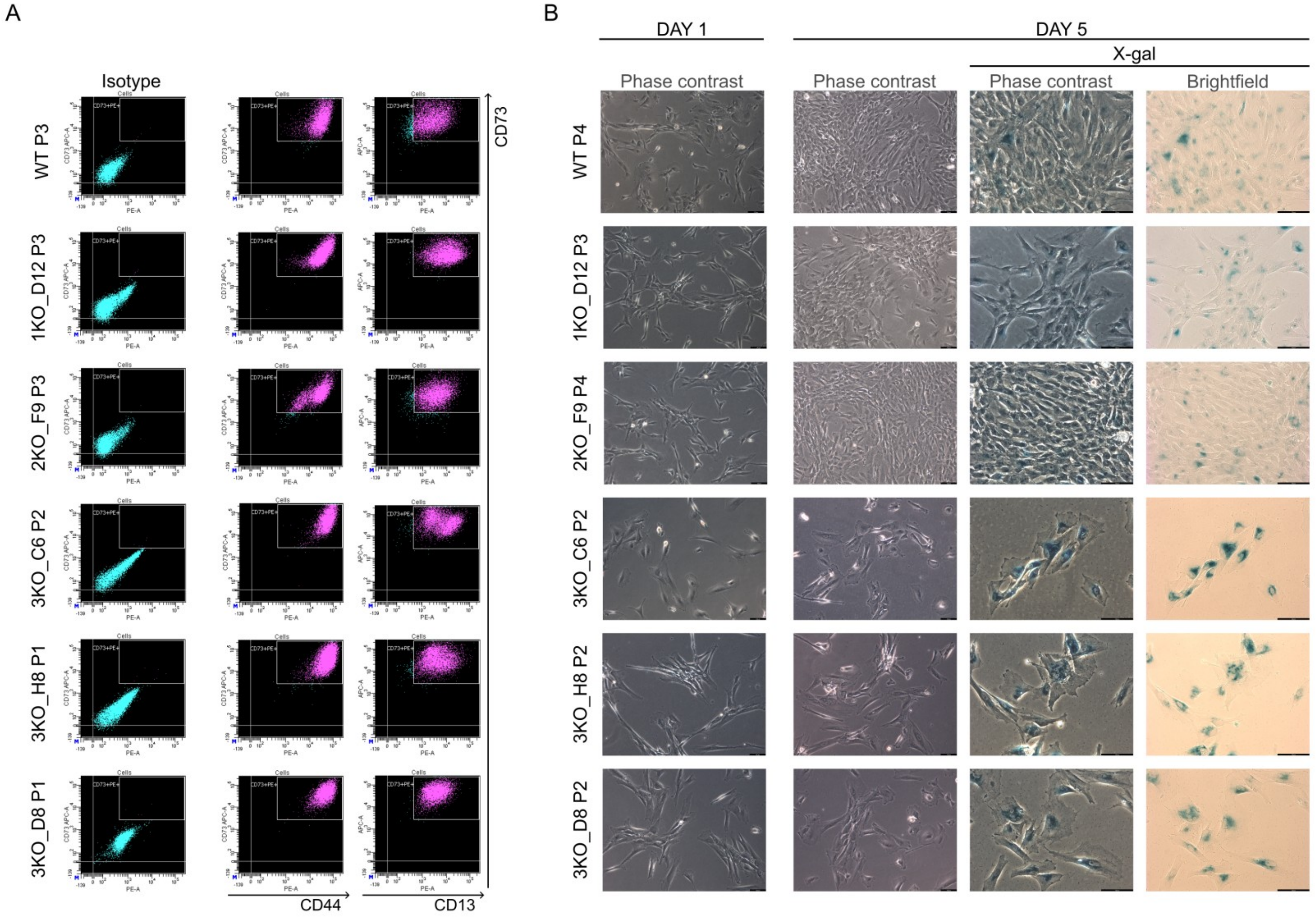


**Figure S4. 3KO NC cells undergo senescence after mesenchymal differentiation, related to Figure 5. A)** Flow cytometry analysis for MSC markers (CD13, CD44 and CD73) in 3KO cell lines and parental lines (WT, 1KO_D12, 2KO_F9) after differentiating them from NC towards MSCs. P: passage. **B)** Day 1: Micrographs of all cell lines at MSC stage. Day 5: Micrographs showing the growth progression and senescence detection by X-Gal staining (blue) in each cell line. Scale bar: 100 µm. P: passage.

Figure S5

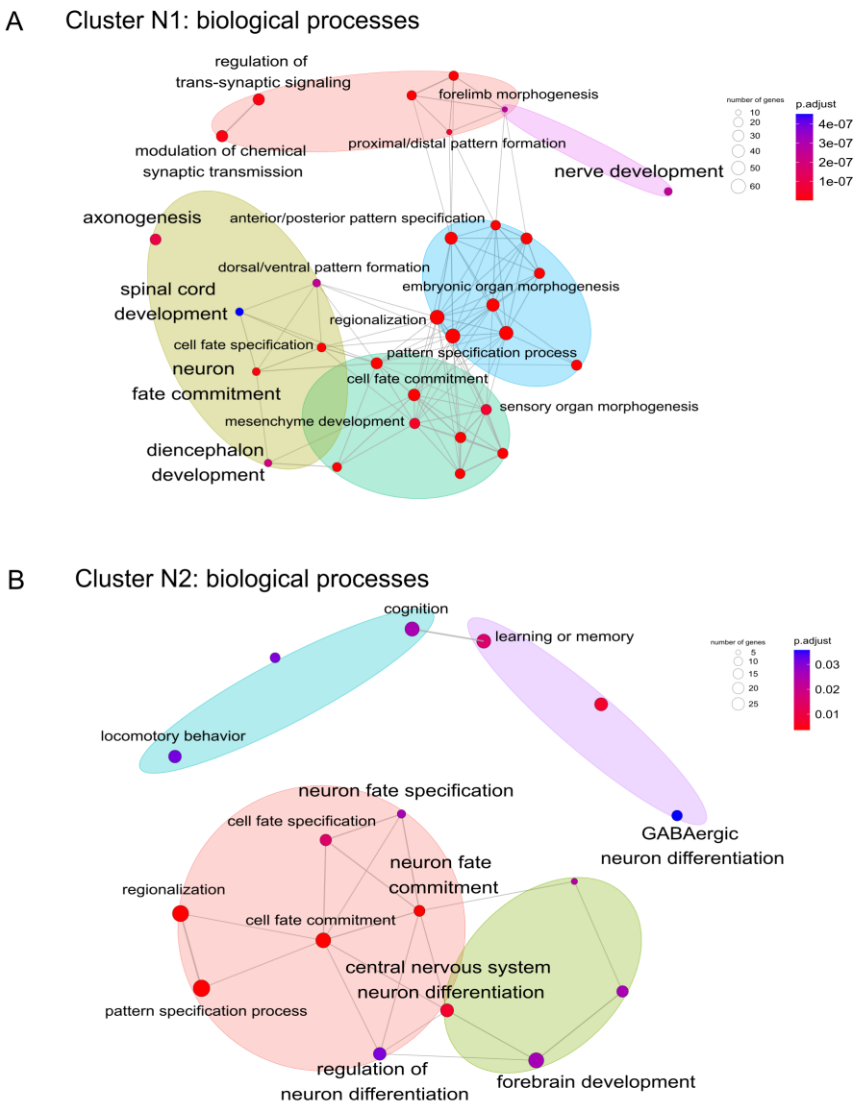


**Figure S5. Biological processes of Cluster N1 and N2, related to Figure 5. A)** Enrichment analysis of genes in Cluster N1 from (Figure 5D). Ellipses represent clusters of related biological processes. See also File S3. **B)** Enrichment analysis of Cluster N2 from (Figure 5G). Ellipses represent clusters of related biological processes. See also File S5.

Figure S6

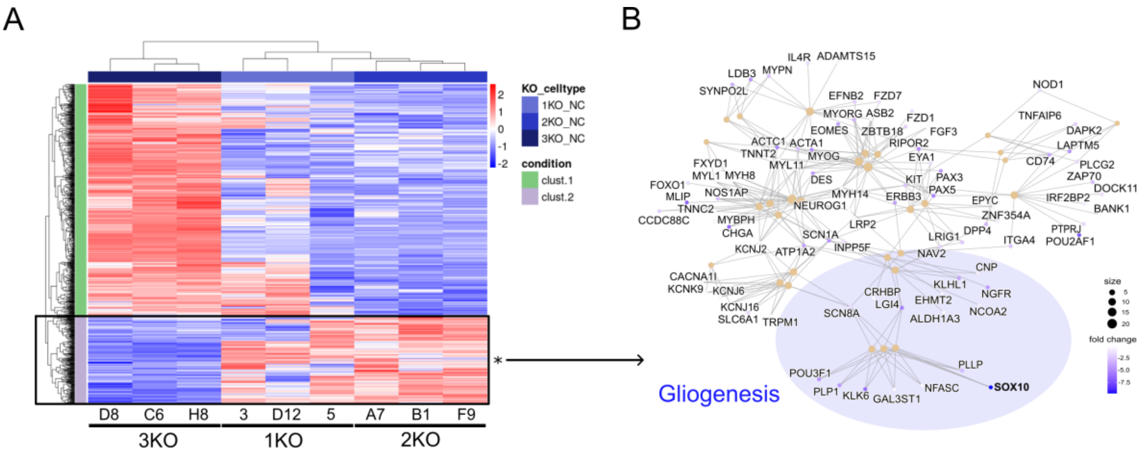


**Figure S6. Gliogenesis downregulation, related to Figure 5. A)** Heatmap showing the differentially expressed genes of 2KO NC vs 3KO NC cells (three independent lines from each genotye are shown). See also File S4. **B)** Enrichment analysis of downregulated genes in 3KO cell lines displayed in (A), highlighting gliogenesis-related genes, like *SOX10*.

Figure S7

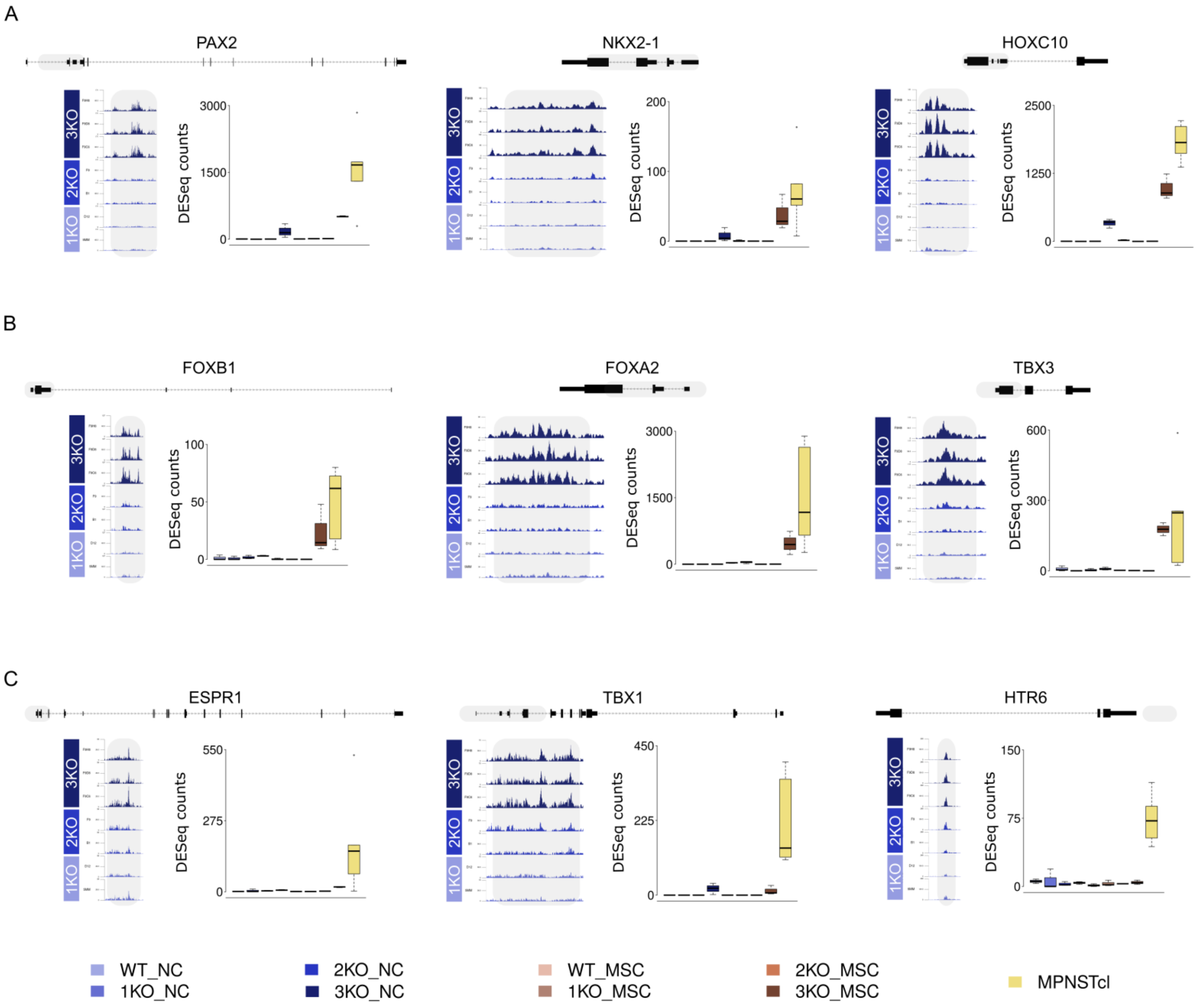


**Figure S7. Gene expression and chromatin accessibility of selected Cluster N2 genes, related to Figure 6. A-C)** Chromatin accessibility profiles in 1KO, 2KO, and 3KO NC cell lines and boxplots illustrating gene expression across all cell lines. Gene schematics are provided, with shaded regions indicating accessible chromatin domains. **A)** Genes expressed in 3KO NCs and further increased in MPNST lines (*PAX2*, *NKX2-1* and *HOXC10)*. **B)** Genes exclusively expressed in 3KO MSCs and MPNST cell lines (*FOXB1*, *FOXA2* and *TBX3*). **C)** Genes expressed only in MPNST cell lines (*ESPR1*, *TBX1* and *HTR*6)

Figure S8

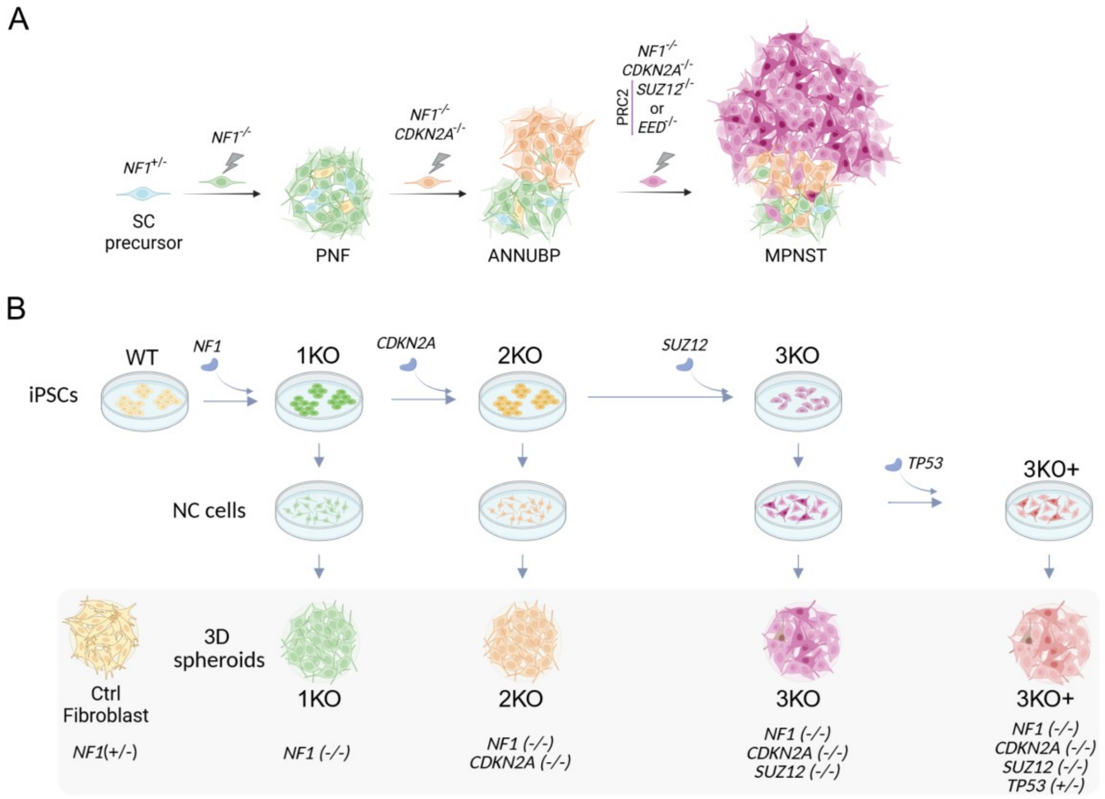


**Figure S8. Representation of the PNF-ANNUBP-MPNST progression and the iPSC-derived 3D NC models used for HTS. A)** Schematic representation of the PNF-ANNUBP-MPNST progression in the context of NF1, highlighting TSG losses. **B)** Schematic representation of the iPSC-derived 3D NC cellular models used for the HTS. *NF1* (+/–) fibroblasts (Ctrl Fibroblast) are used as a control cell line.

Figure S9

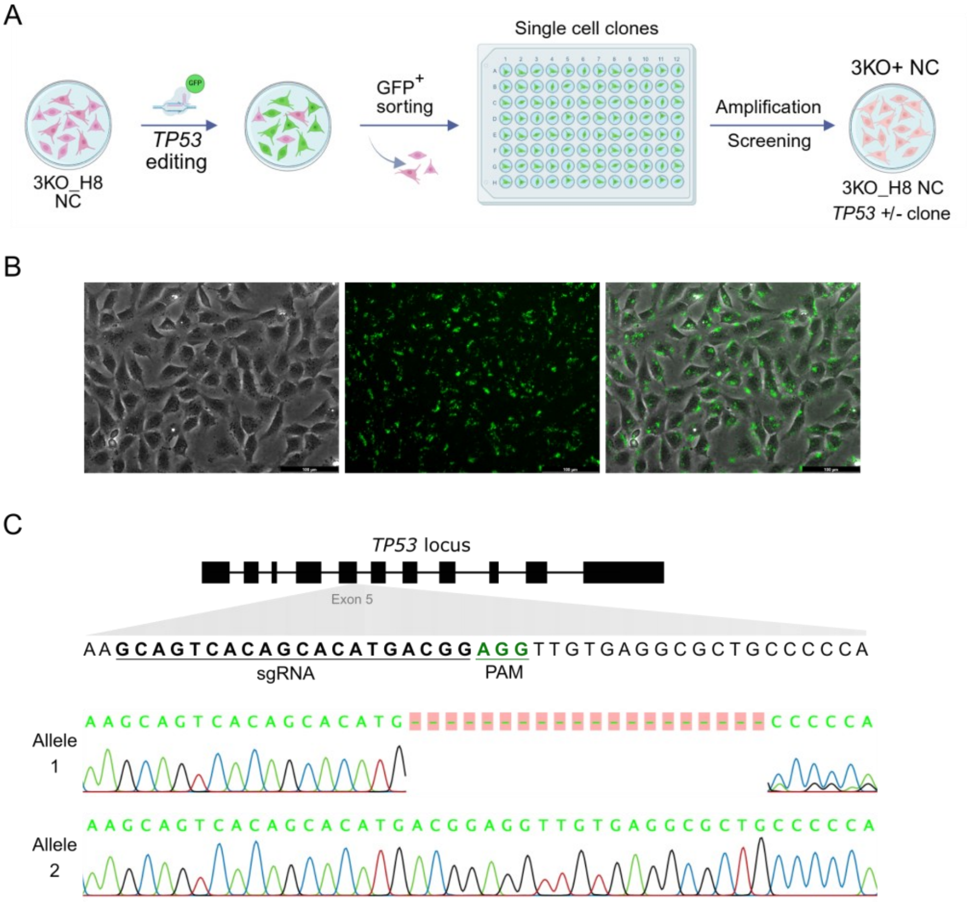


| Clone | Allele | gDNA mutation | cDNA mutation | Protein |
|---|---|---|---|---|
| 3KO+_F3 | Allele 1 | g. 17447_17466del | c. 508_527del | p. T170_Pfs*4 |
| | Allele 2 | WT | WT | WT |

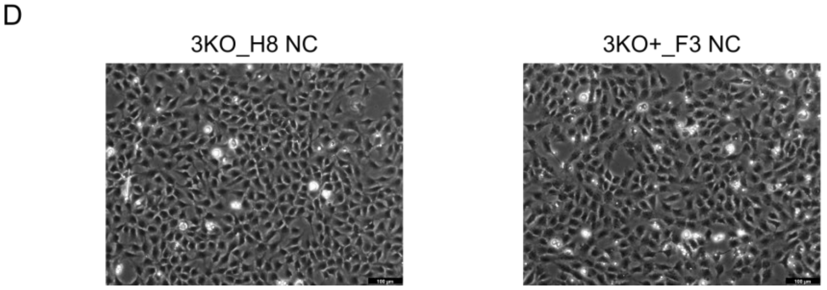


**Figure S9. CRISPR-Cas9-mediated *TP53* gene editing in 3KO cell lines. A)** Schematic representation of the editing strategy. The *TP53* gene was edited in the 3KO-H8 NC cell line using the Architect Ribonucleic Protein (RNP) complex system (StemCell technologies). **B)** Representative micrograph showing RNP-GFP transfection in the 3KO_H8 NC cell line. Scale bar: 100 µm. **C)** Schematic representation of the *TP53* gene showing the location of the designed gRNA in exon 5. Below, Sanger sequencing analysis showing edited mutations in one *TP53* allele. The table below indicates the exact genomic and cDNA location of the mutation and the resulting protein of this mutation. The reference sequences used for alignment and analysis were NG_017013.2 (genomic), NM_000546.5 (mRNA) and NP_000537.3 (protein). **D)** Representative micrographs showing the morphology of the 3KO NC parental line and the newly generated 3KO+ NC cell line. Scale bar: 100 µm.

Figure S10

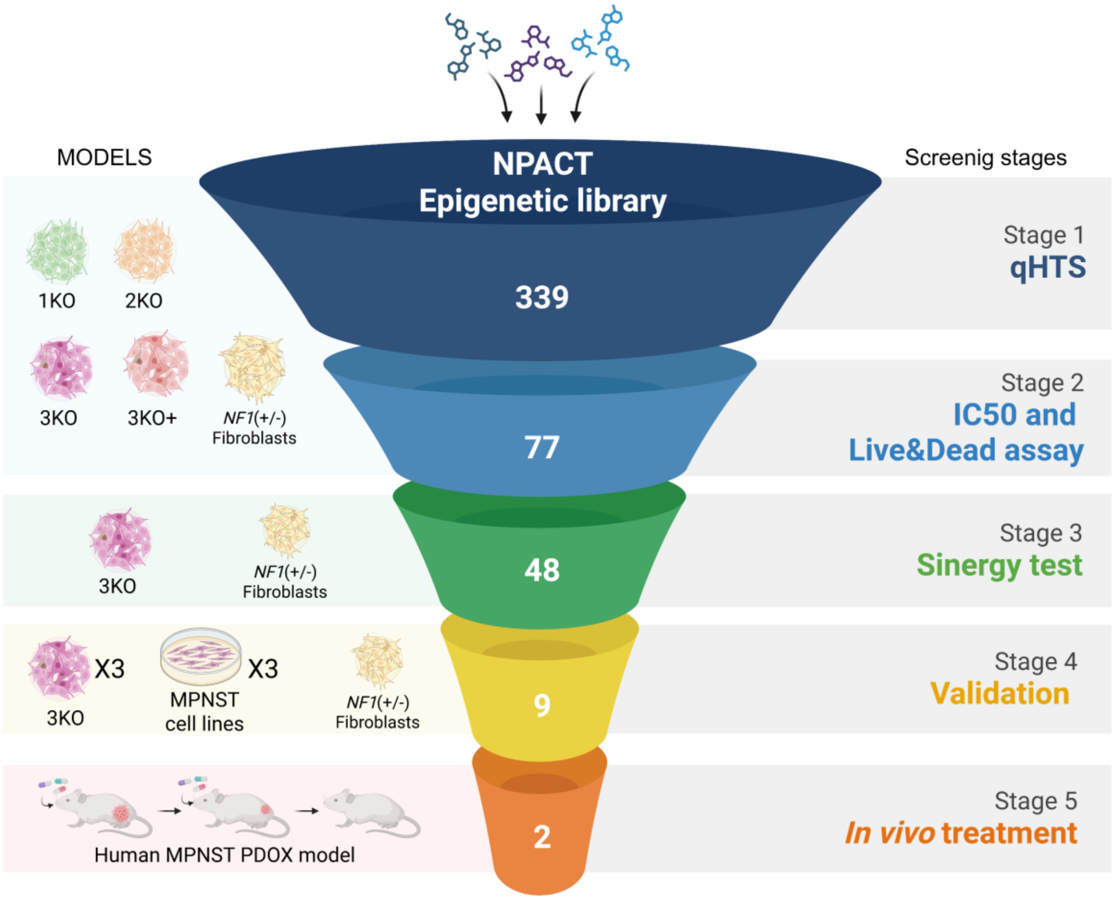


**Figure S10. Screening pipeline and cell lines used at each step.** Left section: schematic representation of the cellular models, including the four isogenic NC lines (1KO, 2KO, 3KO, and 3KO+) grown in 3D as spheroids and fibroblast control spheroids, as well as MPNST cell lines, grown in 2D. Middle section: selected compounds in each HTS step. Right section: screening steps.

Figure S11

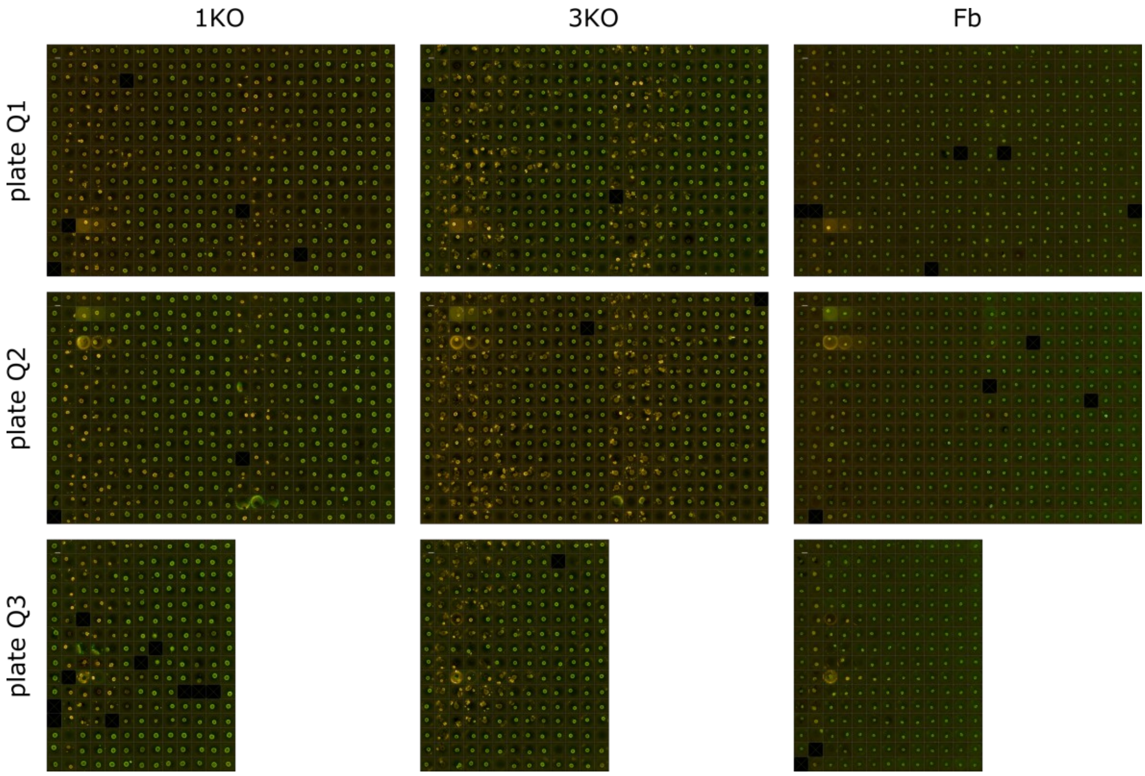


**Figure S11. Live imaging from the compound selection assay across all tested doses, related to Figure 8.** Representative fluorescence micrographs from 384-well live/dead assays obtained after 48 h of treatment of 1KO, 3KO, and control fibroblast (Fb). Green: live cells; yellow: dead cells. Plate maps indicating compounds and corresponding doses are provided in File S5. Scale bar: 500 µm.

Figure S12

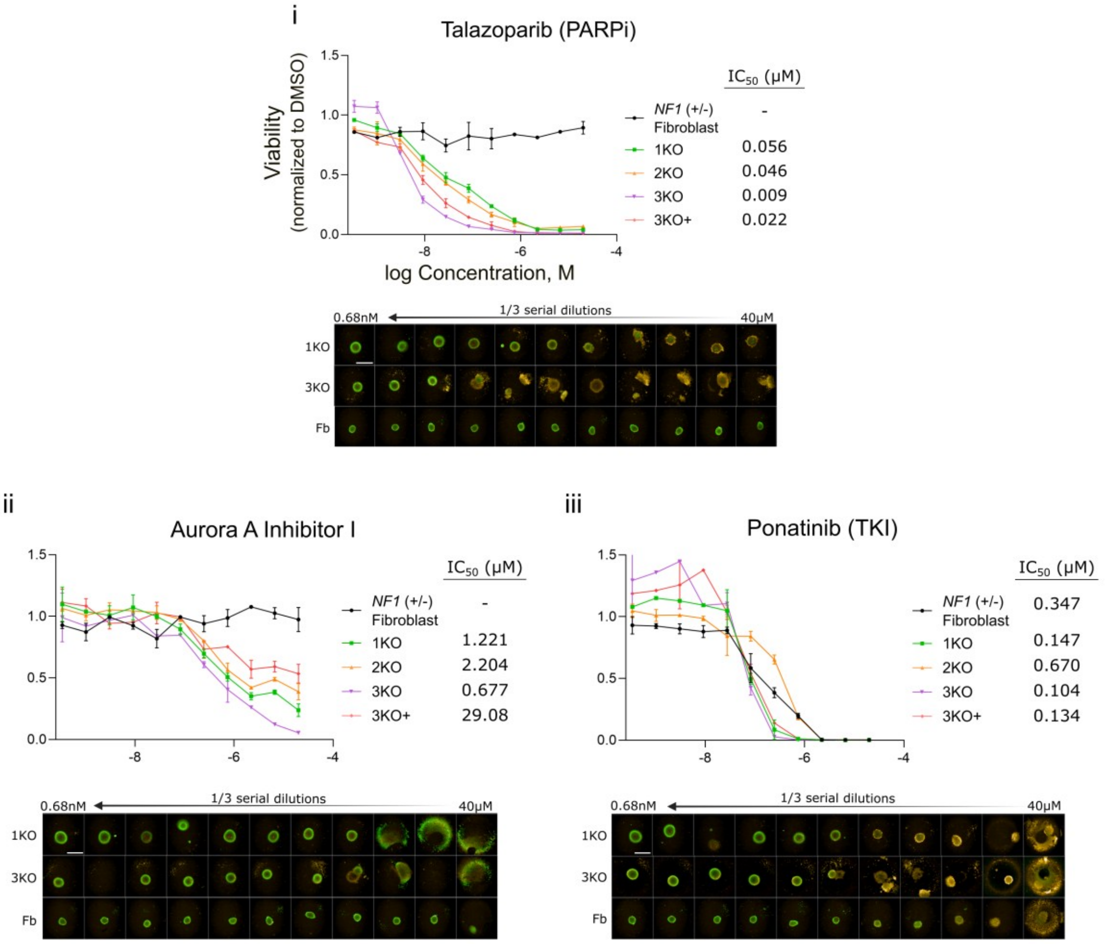


**Figure S12. Dose–response analysis and spheroid imaging of selected compounds from the HTS across all cell lines, related to Figure 8.** Upper panel: Dose–response CellTiter-Glo 3D (CTG) cell viability graphs of the four isogenic NC cell lines (1KO, 2KO, 3KO, and 3KO+) and control fibroblasts, grown in 3D, along with corresponding $IC_{50}$ values. Data correspond to eleven dose–response concentrations; each point shows the mean ± SD from two independent replicates. Bottom panel: Live/dead micrographs for each dose in 1KO, 3KO, and control fibroblasts (Fb). Green: live cells; yellow: dead cells. Scale bar: 500 µm. (**i**) Talazoparib (PARPi), (**ii**) Aurora A inhibitor I (AURKAi), and (**iii**) Ponatinib (TKI).

Figure S13

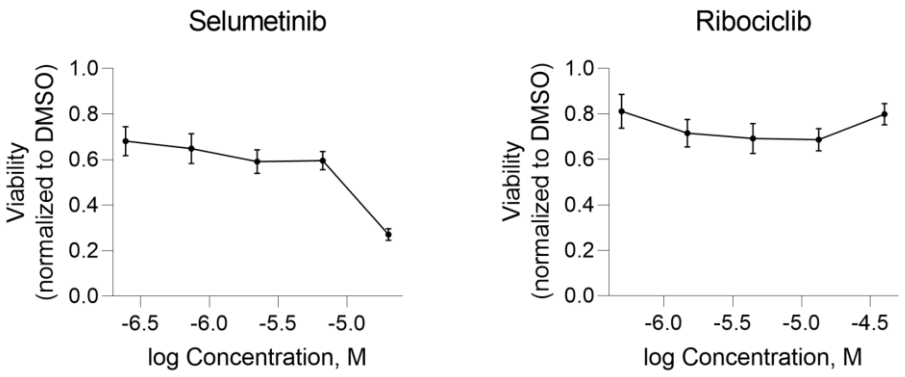


**Figure S13. Dose–response CTG cell viability curves for Selumetinib (MEKi) and Ribociclib (CDKi) in 3KO NC spheroids, related to Figure 8.** Data correspond to five dose–response concentrations from the single-treatment condition in the 6 X 6 blocks of dose-response matrices; each point shows the mean ± SD from fifteen independent replicates.

Figure S14

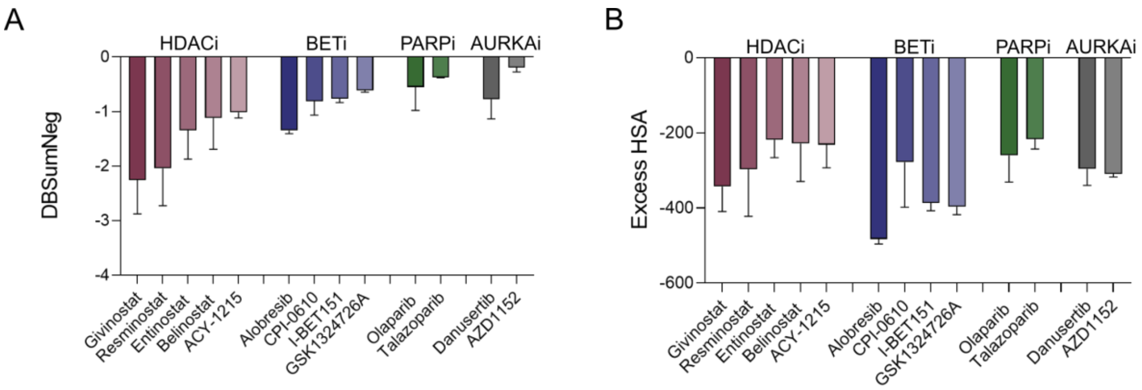


**Figure S14. Synergy graphs of candidate compounds in combination with Selumetinib (MEKi). A)** Delta Bliss Sum Negative (DBSumNeg) values for cell viability of the 13 selected compounds combined with Selumetinib (MEKi) in 3KO spheroids (n=3). **B)** Excess Highest Single Agent (HSA) values for cell viability of the 13 selected compounds combined with Selumetinib (MEKi) in 3KO spheroids (n=3). See also File S10.

Figure S15

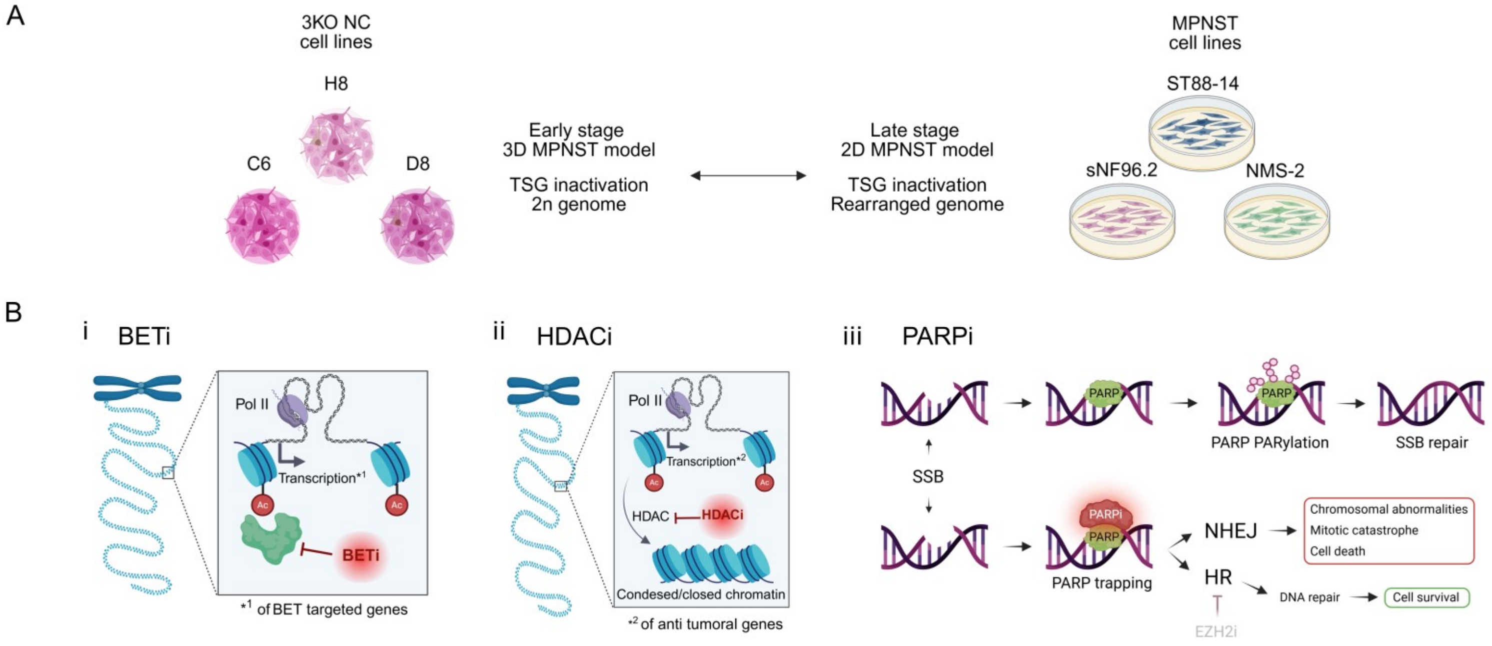


**Figure S15. Schematic representation of the cellular models and mechanisms of action of compound classes used for the HTS validation, related to Figure 8. A)** Schematic overview of the MPNST models used for validation, including their genomic characteristics: early-stage MPNST model (3D 3KO cell lines) have a diploid (2n) genome, and late-stage MPNST model (2D MPNST cell lines) have rearranged genomes. **B)** Schematic representation of the mechanism of action of BETi (**i**), HDACi (**ii**) and PARPi (**iii**).

Figure S16

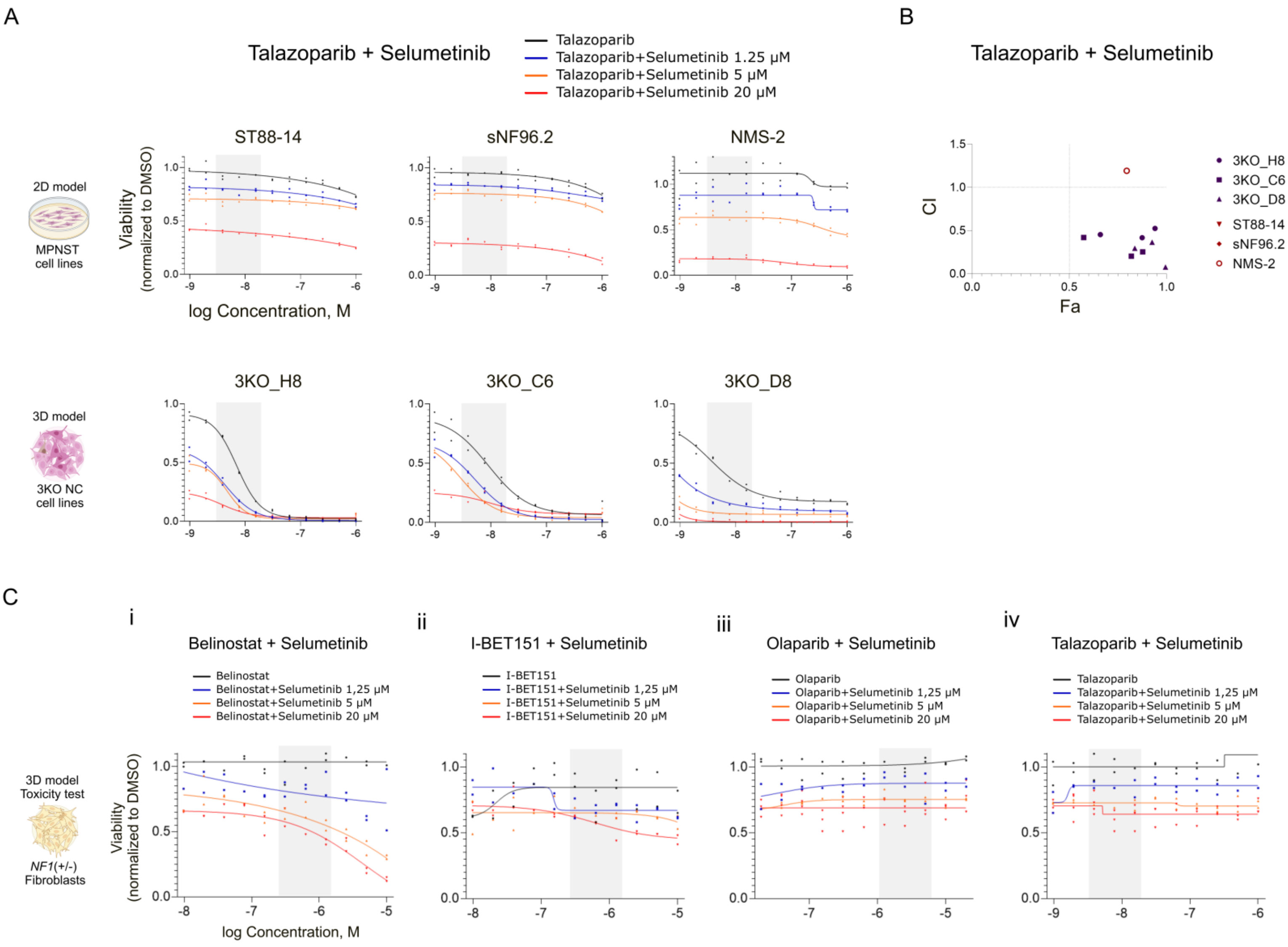


**Figure S16. Validation of candidate compound combinations in 3KO NC cell lines (early-stage MPNST 3D model), compared to MPNST cell lines (late-stage MPNST 2D model), related to Figure 8. A)** Dose–response CTG cell viability curves of Talazoparib (PARPi) in combination with Selumetinib (MEKi) for each 3KO cell line (3KO_H8, 3KO_C6, 3KO_D8; upper panel) and each MPNST cell line (ST88-14, sNF96.2, NMS-2; lower panel). Shaded areas indicate three concentrations above and below the $IC_{50}$ of the compounds in 3KO_H8 cell line. Data correspond to eleven dose–response concentrations; each point shows the mean ± SD from two independent replicates. **B)** Combination index (CI) plots of the shaded doses from panel (A), showing the fraction affected (Fa) versus CI for the 3KO NC (purple) and MPNST (red) cell lines. Fa represents the fraction of cell death induced by drug treatment, ranging from 0 (no cell death) to 1 (complete cell killing). CI values <0.9 indicate a synergistic interaction, values between 0.9 and 1.1 indicate an additive effect, and values >1.1 indicate antagonism. **C)** Dose–response CTG cell viability curves of candidate compounds in combination with Selumetinib (MEKi) in control fibroblasts grown in 3D, showing compound toxicity. Shaded areas indicate three concentrations above and below the $IC_{50}$ of the compounds in 3KO_H8 cell line. Data correspond to eleven dose–response concentrations; each point shows the mean ± SD from two independent replicates. Compounds: **(i)** I-BET151 (BETi), **(ii)** Belinostat (HDACi), **(iii)** Olaparib (PARPi), and **(iv)** Talazoparib (PARPi).

Figure S17

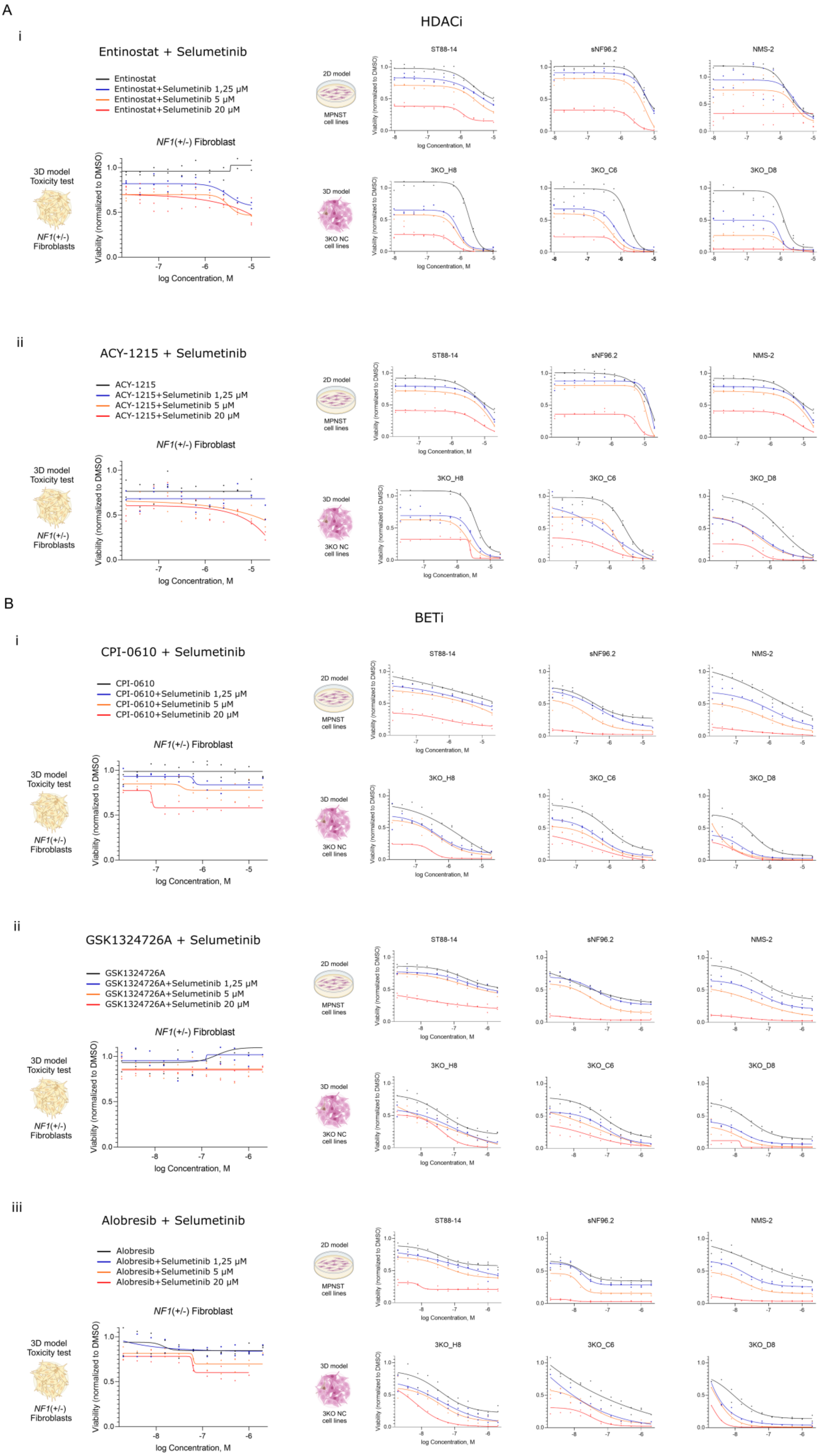
A
i
HDACi
Entinostat + Selumetinib
Entinostat
Entinostat+Selumetinib 1,25 µM
Entinostat+Selumetinib 5 µM
Entinostat+Selumetinib 20 µM
NF1(+/-) Fibroblast
3D model
Toxicity test
NF1(+/-)
Fibroblasts
Viability (normalized to DMSO)
log Concentration, M
2D model
MPNST
cell lines
3D model
3KO NC
cell lines
ST88-14
sNF96.2
NMS-2
3KO_H8
3KO_C6
3KO_D8
ii
ACY-1215 + Selumetinib
ACY-1215
ACY-1215+Selumetinib 1,25 µM
ACY-1215+Selumetinib 5 µM
ACY-1215+Selumetinib 20 µM
B
BETi
i
CPI-0610 + Selumetinib
CPI-0610
CPI-0610+Selumetinib 1,25 µM
CPI-0610+Selumetinib 5 µM
CPI-0610+Selumetinib 20 µM
ii
GSK1324726A + Selumetinib
GSK1324726A
GSK1324726A+Selumetinib 1,25 µM
GSK1324726A+Selumetinib 5 µM
GSK1324726A+Selumetinib 20 µM
iii
Alobresib + Selumetinib
Alobresib
Alobresib+Selumetinib 1,25 µM
Alobresib+Selumetinib 5 µM
Alobresib+Selumetinib 20 µM

Figure S17

**Figure S17. Validation of remaining compound combinations in 3KO cell lines (early-stage MPNST model), compared to MPNST cell lines (late-stage MPNST model), and control fibroblasts, related to Figure 8. A)** Dose response CTG cell viability curves of the remaining HDACi candidate compounds in combination with Selumetinib (MEKi) in control fibroblast (left panel), 3KO NC spheroids (3KO_H8, 3KO_C6, 3KO_D8; upper panel) and MPNST cell lines (ST88-14, sNF96.2, NMS-2; lower panel). (**i**) Entinostat; (**ii**) ACY-1215. Data correspond to eleven dose–response concentrations; each point shows the mean ± SD from two independent replicates. **B)** Dose response CTG cell viability curves of the remaining BETi candidate compounds in combination with Selumetinib (MEKi) in control fibroblast (left panel), 3KO NC spheroids (3KO_H8, 3KO_C6, 3KO_D8; upper panel) and MPNST cell lines (ST88-14, sNF96.2, NMS-2; lower panel). (**i**) CPI-0610; (**ii**) GSK1324726A; (**iii**) Alobresib. Data correspond to eleven dose–response concentrations; each point shows the mean ± SD from two independent replicates.

Figure S18

A

| Compound | Type | Dose (mg/kg) | Schedule (days/week)* | Solvent | Administration route |
|---|---|---|---|---|---|
| Olaparib | PARPi | 45 | 5 | 5% Tween80 + 5% DMSO 40% PEG300 + 50% Salinel | Oral |
| Selumetinib | MEKi | 55 | 5 | 5% Tween80 + 1% DMSO 30% PEG300 + 64% Salinel | Oral |
| I-BET151 | BETi | 22 | 3 | 5% Tween80 + 5% DMSO + 90% Salinel | Intraperitoneal |
| Ribociclib | CDK4/6i | 85 | 5 | 0.5% Methylcellulose + 99.5% Salinel | Oral |

*Single dose per administration day

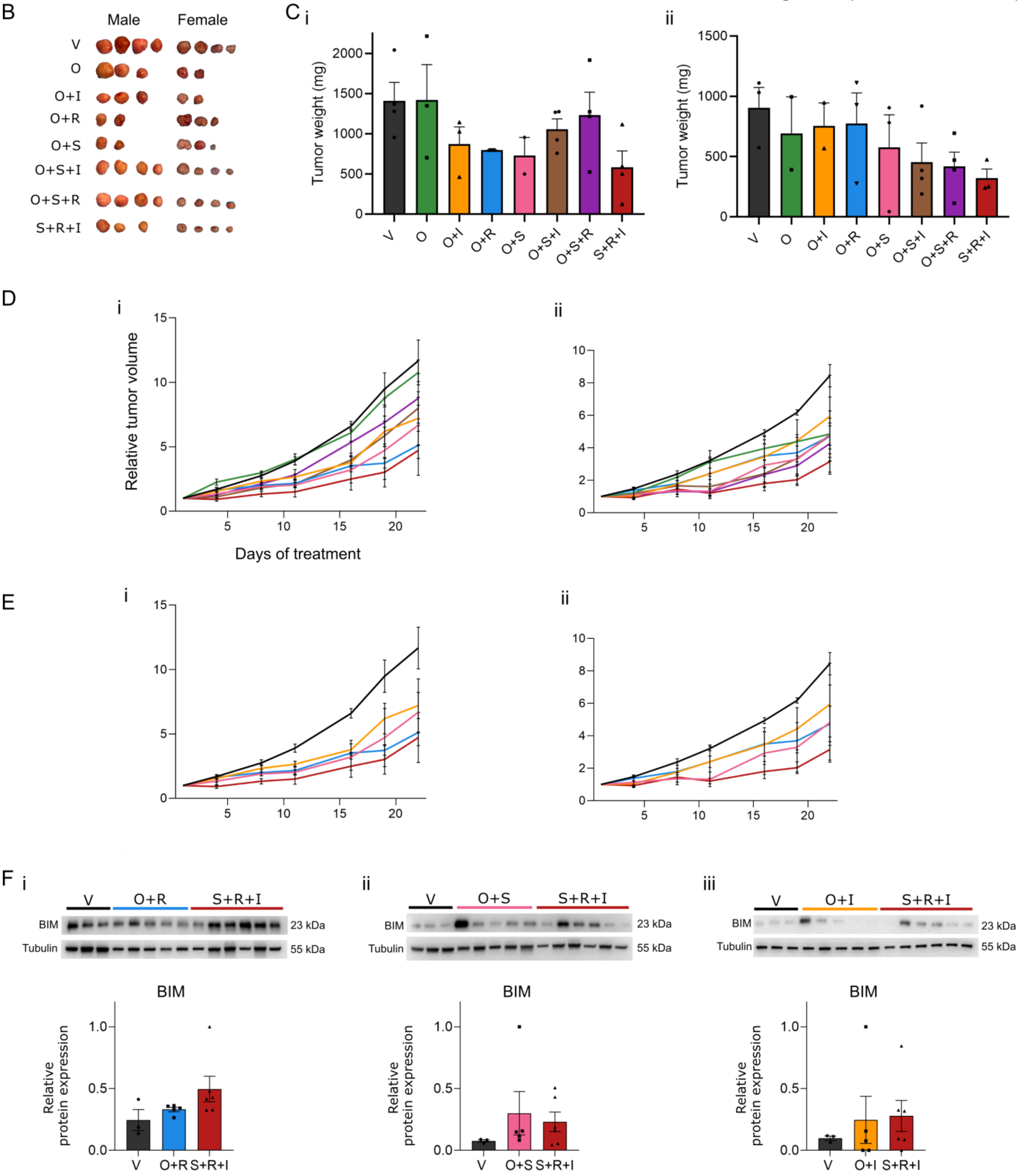

Figure S18

**Figure S18. *In vivo* evaluation of Olaparib (PARPi) alone and in double or triple combination with Selumetinib (MEKi), Ribociclib (CDKi), and I-BET151 (BETi) in the NF1-associated NF1-18B MPNST PDOX mouse model, related to Figure 9. A)** Summary of dosages and administration schedule tested for the three compounds used in the PDOX model. *Single dose per administration day. **B)** Photographs of tumors collected at the end of the experiment. Left panel: Males. Right panel: Females. **C)** Tumor weight at the end of the experiment (Mann–Whitney test). Each black dot indicates one single value. (**i**) Males; (**ii**) Females. **D)** Relative tumor volume growth of the NF1-18B PDOX mouse model in each treatment group over the course of the study. (**i**) Males; (**ii**) Females. **E)** Relative tumor volume growth for each double combination, vehicle control (negative control) and triple Selumetinib + Ribociclib + I-BET151 combination (positive control) throughout the experiment. (**i**) Males; (**ii**) Females. **F)** Western blot analyses and quantification of BIM protein expression in tumors from the NF1-18B PDOX model after treatment with each double combination, vehicle control (negative control) and triple Selumetinib + Ribociclib + I-BET151 combination (positive control).

### Table S1: List of the antibodies.

| Protein Target | Assay | RRID | Source | Reference |
|---|---|---|---|---|
| Phospo-Rb Ser780 | WB-Chemiluminiscence | RRID:AB_330015 | Cell Signaling Technology | Cat# 9307 |
| Rb 4H1 | WB-Chemiluminiscence | RRID:AB_823629 | Cell Signaling Technology | Cat# 9309 |
| c-Myc (D84C12) | WB-Chemiluminiscence | RRID:AB_1903938 | Cell Signaling Technology | Cat# 5605 |
| BIM (C34C5) | WB-Chemiluminiscence | RRID:AB_1030947 | Cell Signaling Technology | Cat# 2933 |
| Vinculin | WB-Chemiluminiscence | RRID:AB_477582 | Sigma-Aldrich | Cat#T60474 |
| Tubulin | WB-Chemiluminiscence/Fluorescence | RRID:AB_10603627 | Sigma-Aldrich | |
| Goat anti-Rabit IgG(H+L) Secondary Antibody, HRP | WB-Chemiluminiscence | RRID:AB_1185567 | Thermo Fisher Scientific | Cat# 32460 |
| Goat anti-Mouse IgG(H+L) Secondary Antibody, HRP | WB-Chemiluminiscence | RRID: AB_1185566 | Thermo Fisher Scientific | Cat# 32430 |
| Poly/Mono-ADP Ribose (D9PTZ) | WB-Fluorescence | RRID:AB_2749858 | Cell Signaling Technology | Cat# 89190 |
| Poly(ADP-Ribose) Polymerase-1 antibody \| A6.4.12 | WB-Fluorescence | RRID:AB_2236751 | BIO-RAD | Cat# MCA1522G |
| Phospho-p44/42 MAPK (Erk1/2) (Thr202/Tyr204) (20G11) | WB-Fluorescence | RRID:AB_331772 | Cell Signaling Technology | Cat# 4376 |
| Purified Mouse Anti-ERK (pan ERK) | WB-Fluorescence | RRID:AB_397529 | BD Transduction Laboratories | Cat#610123 |
| IRDye® 680RD Goat anti-Rabbit IgG | WB-Fluorescence | RRID:AB_10956166 | LICORbio | Cat# 926-68071 |
| IRDye® 800CW Goat Anti-Mouse IgG | WB-Fluorescence | RRID:AB_621842 | LICORbio | Cat# 926-32210 |
| Mouse IgG anti-Nerve growth factor (p75) receptor | Flow cytometry | RRID: AB_303531 | Abcam | Cat# ab3125 |
| OCT3/4 | Immunocytochemistry | RRID: AB_2801346 | StemCell Technologies | Cat# 60093 |
| Mouse IgG anti-AP2 | Immunocytochemistry | RRID: AB_2199412 | Thermo Scientific | Cat# MA1-872 |
| Anti-SOX10 antibody [EPR4007] | Immunocytochemistry | RRID: AB_2650603 | Abcam | Cat# ab155279 |
| Mouse Anti-human SOX9 Monoclonal antibody [3C10] | Immunocytochemistry | RRID: AB_2194156 | Abcam | Cat# ab76997 |
| Rabbit IgG anti-NF1 | WB-Fluorescence | RRID: AB_2149790 | Bethyl laboratories | Cat# A300-140a |
| CDKN2A/p16INK4a antibody | WB-Fluorescence | RRID: AB_10858268 | Abcam | Cat# ab108349 |
| Rabbit IgG anti-Ku80 | Immunohistochemistry | RRID: AB_2218736 | Cell Signaling Technology | Cat# 2180 |
| Rabbit IgG anti-S100B | Histochemistry | RRID: AB_10013383 | DAKO | Cat# Z0311 |
| Mouse monoclonal S100B | Histochemistry | RRID: AB_301508 | Abcam | ab14849 |
| Mouse IgG anti-HNK1 | Flow cytometry | RRID: AB_1078474 | Sigma | Cat# C6680 |
| Tri-Methyl-Histone H3 (Lys27) (C36B11) Rabbit mAb | Immunocytochemistry | RRID: AB_2616029 | Cell Signaling Technology | Cat# 9733 |
| PE Mouse Anti-CD44 | Flow cytometry | RRID: AB_10898347 | BD Pharmigen | Cat# 561858 |
| PE Mouse Anti-CD13 | Flow cytometry | RRID: AB_10563611 | BD Pharmigen | Cat# 560998 |
| APC Mouse Anti-CD73 | Flow cytometry | RRID: AB_10612019 | BD Pharmigen | Cat# 560847 |
| VIMENTIN Monoclonal Antibody, Clone V9 | Histochemistry | RRID: AB_138999 | Life Technologies | Cat# 180052 |
| Rabbit monoclonal Ki67 (clone 30-9) | Histochemistry | RRID: AB_2631262 | Ventana Medical Systems | Cat# 790-4286 |
| Goat anti-Mouse IgG (H+L) Alexa fluor 488 | Immunocytochemistry/ Immunohistochemistry/ Flow cytometry | RRID: AB_2534069 | Thermo Scientific | Cat# A-11001 |
| Goat anti-rabbit IgG (H+L) Alexa fluor 568 | Immunocytochemistry/ Immunohistochemistry/ Flow cytometry | RRID: AB_143157 | Thermo Scientific | Cat# A-11011 |

**Table S2: Z' values across all cell lines in the qHTS and in the secondary screening**. Each Z' has been calculated using Panobinostat as a positive control (n=16) and DMSO as a negative control (n=16). Q1-Q8 represents each point in the qHTS. Q1-Q3 represents each compound plate of the secondary screening.

| | | qHTS | | | | | | | | Secondary screening | | | | | | |
|---|---|---|---|---|---|---|---|---|---|---|---|---|---|---|---|---|
| | | **Q1** | **Q2** | **Q3** | **Q4** | **Q5** | **Q6** | **Q7** | **Q8** | **Q1 rep1** | **Q1 rep2** | **Q2 rep1** | **Q2 rep2** | **Q3 rep1** | **Q3 rep2** | **Mean Z'** |
| **Cell line** | **1KO** | 0.79 | 0.79 | 0.81 | 0.75 | 0.82 | 0.86 | 0.83 | 0.83 | 0.88 | 0.83 | 0.85 | 0.75 | 0.85 | 0.75 | **0.755** |
| | **2KO** | 0.61 | 0.75 | 0.78 | 0.79 | 0.50 | 0.75 | 0.78 | 0.77 | 0.84 | 0.80 | 0.85 | 0.85 | 0.89 | 0.79 | **0.768** |
| | **3KO** | 0.86 | 0.72 | 0.72 | 0.87 | 0.82 | 0.86 | 0.74 | 0.81 | 0.86 | 0.85 | 0.85 | 0.77 | 0.78 | 0.82 | **0.802** |
| | **3KO+** | 0.77 | 0.86 | 0.75 | 0.71 | 0.79 | 0.75 | 0.68 | 0.75 | 0.85 | 0.76 | 0.72 | 0.84 | 0.83 | 0.82 | **0.777** |
| | **Fb** | 0.86 | 0.82 | 0.82 | 0.85 | 0.77 | 0.86 | 0.85 | 0.80 | 0.81 | 0.86 | 0.67 | 0.84 | 0.82 | 0.80 | **0.816** |

**Table S3: qHTS results summarizing CRC and $IC_{50}$ values across different compound types in all cell lines.**

**1KO cell line**

| | CRC | | | | | $IC_{50}$ (µM) | | |
|---|---|---|---|---|---|---|---|---|
| | -1.1 | -1.2 | -2.1 | -2.2 | Inactive | $IC_{50} \geq 13$ | $IC_{50} < 13$ | TOTAL |
| BETi | 13 | 0 | 0 | 0 | 5 | 6 | 12 | 18 |
| HDACi | 16 | 0 | 4 | 0 | 10 | 13 | 17 | 30 |
| AURKA/Bi | 8 | 2 | 3 | 1 | 11 | 10 | 15 | 25 |
| PARPi | 1 | 0 | 2 | 1 | 6 | 8 | 2 | 10 |
| JAKi | 5 | 0 | 0 | 2 | 8 | 8 | 7 | 15 |
| EZH2i | 0 | 0 | 1 | 0 | 12 | 13 | 0 | 13 |
| DNMTi | 1 | 0 | 1 | 0 | 8 | 8 | 2 | 10 |
| EHMT2 | 0 | 0 | 2 | 0 | 4 | 5 | 1 | 6 |
| KDMi | 1 | 0 | 0 | 0 | 13 | 13 | 1 | 14 |
| PRMTi | 0 | 0 | 0 | 0 | 6 | 6 | 0 | 6 |
| SIRTi | 0 | 0 | 0 | 0 | 4 | 4 | 0 | 4 |
| PIK3CA/B | 0 | 0 | 0 | 0 | 4 | 4 | 0 | 4 |
| Others | 23 | 1 | 19 | 2 | 139 | 151 | 33 | 182 |
| TOTAL | 68 | 3 | 32 | 6 | 230 | 249 | 90 | 339 |

**2KO cell line**

| | CRC | | | | | $IC_{50}$ (µM) | | |
|---|---|---|---|---|---|---|---|---|
| | -1.1 | -1.2 | -2.1 | -2.2 | Inactive | $IC_{50} \geq 13$ | $IC_{50} < 13$ | TOTAL |
| BETi | 10 | 0 | 1 | 0 | 7 | 7 | 11 | 18 |
| HDACi | 10 | 0 | 2 | 0 | 18 | 9 | 21 | 30 |
| AURKA/Bi | 6 | 3 | 2 | 0 | 14 | 12 | 13 | 25 |
| PARPi | 1 | 0 | 1 | 1 | 7 | 7 | 3 | 10 |
| JAKi | 5 | 0 | 0 | 1 | 9 | 7 | 8 | 15 |
| EZH2i | 0 | 0 | 0 | 0 | 13 | 12 | 1 | 13 |
| DNMTi | 1 | 0 | 3 | 0 | 6 | 7 | 3 | 10 |
| EHMT2 | 0 | 0 | 0 | 0 | 6 | 6 | 0 | 6 |
| KDMi | 1 | 0 | 1 | 0 | 12 | 11 | 3 | 14 |
| PRMTi | 0 | 0 | 0 | 0 | 6 | 5 | 1 | 6 |
| SIRTi | 0 | 0 | 0 | 0 | 4 | 3 | 1 | 4 |
| PIK3CA/B | 0 | 0 | 0 | 1 | 3 | 4 | 0 | 4 |
| Others | 18 | 0 | 14 | 0 | 152 | 142 | 42 | 182 |
| TOTAL | 52 | 3 | 24 | 3 | 257 | 232 | 107 | 339 |

### 3KO cell line

| | CRC | | | | | IC50 (µM) | | |
|---|---|---|---|---|---|---|---|---|
| | -1.1 | -1.2 | -2.1 | -2.2 | Inactive | $IC_{50} \geq 13$ | $IC_{50} < 13$ | TOTAL |
| **BETi** | 9 | 0 | 3 | 0 | 6 | 5 | 13 | 18 |
| **HDACi** | 13 | 0 | 8 | 0 | 9 | 13 | 17 | 30 |
| **AURKA/Bi** | 10 | 2 | 2 | 2 | 9 | 7 | 18 | 25 |
| **PARPi** | 2 | 0 | 4 | 0 | 4 | 7 | 3 | 10 |
| **JAKi** | 5 | 0 | 1 | 0 | 9 | 9 | 6 | 15 |
| **EZH2i** | 0 | 0 | 1 | 1 | 11 | 13 | 0 | 13 |
| **DNMTi** | 1 | 0 | 0 | 0 | 9 | 8 | 2 | 10 |
| **EHMT2** | 0 | 0 | 0 | 0 | 6 | 6 | 0 | 6 |
| **KDMi** | 1 | 0 | 0 | 0 | 13 | 13 | 1 | 14 |
| **PRMTi** | 0 | 0 | 0 | 0 | 6 | 6 | 0 | 6 |
| **SIRTi** | 0 | 0 | 0 | 0 | 4 | 4 | 0 | 4 |
| **PIK3CA/B** | 0 | 0 | 0 | 0 | 4 | 4 | 0 | 4 |
| **Others** | 28 | 2 | 19 | 1 | 134 | 155 | 29 | 182 |
| **TOTAL** | 69 | 4 | 38 | 4 | 224 | 250 | 89 | 339 |

### 3KO+ cell line

| | CRC | | | | | IC50 (µM) | | |
|---|---|---|---|---|---|---|---|---|
| | -1.1 | -1.2 | -2.1 | -2.2 | Inactive | $IC_{50} \geq 13$ | $IC_{50} < 13$ | TOTAL |
| **BETi** | 8 | 0 | 4 | 0 | 6 | 5 | 13 | 18 |
| **HDACi** | 15 | 0 | 5 | 0 | 10 | 17 | 13 | 30 |
| **AURKA/Bi** | 10 | 2 | 1 | 1 | 11 | 8 | 17 | 25 |
| **PARPi** | 1 | 0 | 4 | 0 | 5 | 7 | 3 | 10 |
| **JAKi** | 6 | 0 | 0 | 0 | 9 | 9 | 6 | 15 |
| **EZH2i** | 0 | 1 | 0 | 0 | 12 | 13 | 0 | 13 |
| **DNMTi** | 0 | 0 | 0 | 0 | 10 | 8 | 2 | 10 |
| **EHMT2** | 0 | 0 | 0 | 0 | 6 | 6 | 0 | 6 |
| **KDMi** | 1 | 0 | 0 | 0 | 13 | 13 | 1 | 14 |
| **PRMTi** | 0 | 0 | 0 | 0 | 6 | 6 | 0 | 6 |
| **SIRTi** | 0 | 0 | 0 | 0 | 4 | 4 | 0 | 4 |
| **PIK3CA/B** | 0 | 0 | 1 | 0 | 3 | 3 | 1 | 4 |
| **Others** | 21 | 0 | 10 | 2 | 150 | 154 | 30 | 182 |
| **TOTAL** | 62 | 3 | 26 | 3 | 245 | 253 | 86 | 339 |

***NF1*(+/-) fibroblast control cell line**

| | CRC | | | | | IC50 (µM) | | |
|---|---|---|---|---|---|---|---|---|
| | **-1.1** | **-1.2** | **-2.1** | **-2.2** | **Inactive** | **$IC_{50} \geq 13$** | **$IC_{50} < 13$** | **TOTAL** |
| **BETi** | 0 | 0 | 1 | 0 | 17 | 17 | 1 | 18 |
| **HDACi** | 0 | 1 | 2 | 1 | 26 | 28 | 2 | 30 |
| **AURKA/Bi** | 5 | 3 | 4 | 0 | 13 | 16 | 9 | 25 |
| **PARPi** | 0 | 0 | 0 | 0 | 10 | 10 | 0 | 10 |
| **JAKi** | 2 | 1 | 6 | 0 | 6 | 9 | 6 | 15 |
| **EZH2i** | 0 | 0 | 1 | 0 | 12 | 13 | 0 | 13 |
| **DNMTi** | 1 | 0 | 2 | 0 | 7 | 9 | 1 | 10 |
| **EHMT2** | 1 | 0 | 2 | 0 | 3 | 4 | 2 | 6 |
| **KDMi** | 0 | 0 | 0 | 0 | 14 | 14 | 0 | 14 |
| **PRMTi** | 0 | 0 | 0 | 0 | 6 | 6 | 0 | 6 |
| **SIRTi** | 0 | 0 | 0 | 0 | 4 | 4 | 0 | 4 |
| **PIK3CA/B** | 0 | 1 | 0 | 0 | 3 | 4 | 0 | 4 |
| **Others** | 0 | 1 | 17 | 7 | 160 | 182 | 2 | 182 |
| **TOTAL** | 9 | 7 | 34 | 8 | 284 | 316 | 23 | 339 |

Inactive compounds: CRC -1.3, -1.4 and positive values. Aurora kinase A/B inhibitor (AURKA/Bi); Tyrosine-protein kinase inhibitor (JAKi); Bromodomain-Containing Protein inhibitor (BETi); Histone deacetylase inhibitor (HDACi); histone–lysine N-methyltransferase inhibitors (EZH2i); protein arginine N-methyltransferase inhibitors (PRMTi); DNA methyltransferase inhibitors (DNMTi); Histone-lysine N-methyltransferase inhibitor (EHMT2i); lysine-specific demethylase inhibitors (LSDi/KDMi).

**Additional material and methods**

The following additional sections of materials and methos are available as supplementary:

DNA extraction; RNA extraction; MPNST cell lines and PNF-derived Schwann cells RNAseq data; Plexiform neurofibromas (PNFs), ANNUBPS, MPNSTs and nerve samples; Neural Crest (NC) differentiation towards Mesenchymal Stem Cells (MSC); Neural Crest (NC) differentiation towards Schwann cells (SCs); Immunohistochemical analysis of formalin-fixed paraffin-embedded (FFPE) samples; Fluorescent immunohistochemistry of FFPE samples; Senescence assay; Ploidy assay; Proliferation assay; Lentiviral vectors; Viral production and viral transduction; Pemetrexed treatment; Extended qHTS; Live/dead cell viability assay; Compound Combination Matrix Screening; DNA methylation; Single-cell RNA-Seq and analysis; WGS.

**DNA extraction**

Genomic DNA from cells was extracted using Promega Maxwell 16 system following the manufacturer's instructions (Promega).

**RNA extraction**

Total RNA from cells was extracted using the 16 LEV simplyRNA Purification Kit (Promega) following manufacturer's instructions in the Maxwell 16 Instrument (Promega). RNA was quantified with a Nanodrop 1000 spectrophotometer (Thermo Scientific).

**MPNST cell lines and PNF-derived Schwann cells RNAseq data**

In this study, we used the RNA-seq data of five NF1-associated MPNST cell lines:[1] S462 (RRID:CVCL_1Y70), ST88-14 (RRID:CVCL_8916), F90-8 (RRID:CVCL_1B47), sNF96.2 (RRID:CVCL_K281) and NMS-2 (RRID:CVCL_4662). This MPNST cell lines contains the most frequently inactivated tumor suppressor genes (TSGs): *NF1*, *CDKN2A* and PRC2.

In this study, we used the RNA-seq data of 3 PNF-derived SC lines. [2]

**Plexiform neurofibromas (PNFs), ANNUBPS, MPNSTs and nerve samples**

NF1 patients diagnosed according to standard diagnostic criteria[3] and control patients kindly provided samples after giving written informed consent, with the approval by the Clinical Research Ethics Committee of Germans Trias i Pujol Hospital, Badalona, Spain. Tumor and nerve specimens were obtained after surgery, placed in DMEM medium (Gibco) containing 10% FBS (Gibco) + 1x GlutaMax (Gibco) + 1x Normocin antibiotic cocktail (InvivoGene) and shipped at room temperature to our laboratory. Samples were cut into 1-mm pieces and cryopreserved in 10% DMSO (Sigma) + 90% FBS (Gibco) until used.

**Neural Crest (NC) differentiation towards Mesenchymal Stem Cells (MSC)**

Neural crest (NC) cells were plated at a density of $6.5x10^4$ cells per $cm^2$ in DMEM + 10% FBS + 500 U/mL penicillin/500 mg/mL streptomycin (Gibco) + 1x Glutamax-I 200mM + 90µm 2-bmercaptoethanol onto noncoated tissue culture plates following previously reported protocols.[4] Cells were passaged every 4-5 days using trypsin-EDTA.

**Neural Crest (NC) differentiation towards Schwann cells (SCs)**

Schwann cell (SC) differentiation was performed as reported previously.[5] Briefly, 4x $10^4$ NC cells per $cm^2$ were plated onto 0.1 mg/mL Poly-Llysine (Sigma) and 4 mg/mL Laminin (Gibco)-coated plates and cultured in SC differentiation medium: DMEM:F12 (3:1); 500 U/mL penicillin/500 mg/mL streptomycin (Gibco); 1% FBS (Gibco); 5 µM Forskolin (Sigma); 50 ng/mL Heregulin β1; and 2% N2 supplement (Gibco). The medium was replaced twice a week.

**Immunohistochemical analysis of formalin-fixed paraffin-embedded (FFPE) samples**

Hematoxylin & Eosin (H&E) and S100B, SOX10 and Ki67 immunostaining was performed in the IGTP-Germans Trias i Pujol University Hospital (HUGTP) Biobank Facility and Anatomy Pathology Department. Immunohistochemical staining was performed on glass slides after deparaffination and rehydration of formalin-fixed paraffine-embedded tissue. The automatic processing was performed in a Benchmark Ultra (Ventana). Briefly, rehydrated slides were subjected to heat antigen retrieval in CC1 buffer, incubated with primary antibodies and developed with the Ultraview DAB detection system (Ventana).

For H3K27me and vimentin immunostaining endogenous peroxidases were blocked with hydrogen peroxide ($H_2O_2$ 3% for 20 min), and antigen retrieval was performed by heating tissue sections for 20 min in citrate buffer (pH = 6). Blocking was performed with 10% goat serum for 20 min and primary antibodies, vimentin (1:200) and H3K27me3 (1:200) were incubated overnight at 4 °C. The secondary HPRT-conjugated antibody (EnVision; DAKO) was incubated at RT for 30 min. Finally, staining was conducted using diaminobenzidine (DAB; DAKO) for 10 min; nuclei were counterstained with hematoxylin. Images were taken using a Nikon Eclipse 80i vertical microscope. See Table S1 for antibody details.

**Fluorescent immunohistochemistry of FFPE samples**

FFPE Sections were rehydrated using standard procedures, and antigen retrieval was performed by heating tissue sections for 20 min in citrate buffer (pH = 6). Samples were permeabilized in 0.3% Triton-X in PBS for 30 min, blocked in 1% bovine serum albumin, 10% goat serum (Gibco), 0.1% Triton-X in PBS for 1 hour, and incubated with primary antibodies (S100B, Ku80) diluted in incubation buffer (1% bovine serum albumin with 10% goat serum in PBS and 0.3% TritonX100) overnight at 4ºC. Then, slides were washed 3 times for 10 minutes each in PBS, incubated with secondary antibodies (Alexa Fluor 488 goat anti-mouse, and Alexa Fluor goat 568 anti-rabbit,

1:1000 each) diluted in incubation buffer for 45 minutes, washed again and incubated with DAPI (Stem Cell Technologies, 1:1000) diluted in PBS for 10 minutes. Sections were mounted with Vectashield (Vector laboratories) and secured with polish nail. Images were captured using the DMI 6000B microscope (Leica) and LAS X software (Leica). See Table S1 for antibody details.

**Senescence assay**

Cells were cultured until they reached 70% confluency and senescent cells were stained using the Senescence detection kit (Sigma CS0030) following manufacturer's instructions. Phase contrast images were captured using the DMI 6000B microscope (Leica) and LAS X software (Leica).

**Ploidy assay**

Five hundred thousand NC cells were seeded on matrigel-coated 6-well plates. After 48h 1 million cells were fixed with 70% cold ethanol overnight at -20°C. Cells were incubated 30min at 37°C with 50µg/ml propidium iodide in PBS-FBS 1% with 100µg/ml RNase and analyzed by flow cytometry using BD FACSCanto II and BD FACSDiva 6.2 software.

**Proliferation assay**

One hundred fifty thousand NC cells were seeded on matrigel-coated 12-well plates. After 48h cells were treated with 20µM EdU for 1 hour, fixed, permeabilized, and click labeled with Alexa Fluor 488 using Click-iT Plus EdU Flow Cytometry Assay Kits (Thermo Fisher) according to the manufacturer instructions. Cells were also stained with propidium iodide to detect DNA content. Data was collected and analyzed using BD FASCCanto II and BD FACSDiva 6.2 software.

**Lentiviral vectors**

We generated a Tet Inducible Gene Expression Systems to re-introduce the expression of p14 and p16. We first introduced the helper lentiviral vector pLV-EF1A>Tet3G(ns):T2A:Neo (VB230505-1096yze); and then the inducible lentiviral vectors coding for either p16 (pLV-TRE3G>hCDKN2A[NM_000077.5]:IRES:mCherry-hPGK>Puro (VB230505-1095kwb)) or p14 (pLV-TRE3G>hCDKN2A[NM_058195.4]:IRES:EGFP-hPGK>Hygro (VB230505-1076bgt)). These three vectors were purchased from VectorBuilder.

**Viral production and viral transduction**

Lentiviral production was performed upon transfection of HEK293T cells using $CaCl_2$ transfection reagent. Packaging vectors used were psPAX2 (Addgene, 12260) and the envelope expression plasmid pCMV-VSV-G (Addgene, #8454). Briefly, 1.4 million of HEK293T cells were seeded in 100mm plates in DMEM 10 %FBS media. 24h later media was changed to NC media and cells were co-transfected with lentiviral vectors. 72h post-transfection, viral supernatant was collected and filtered through a 0.45 µm filter.

Lentiviral transduction was performed in a medium containing viral particles adding 10 mg/mL polybrene for 6 hours. 48h post-transduction cells were selected with antibiotics: G418 (150 µg/mL), Puromycin (0.2-0.4 µg/mL), and Hygromycin (50 µg/mL).

For enrichment of transduced cells, we FACS sorted GFP+ or mCherry+ cells after doxycycline induction (1 µg/mL) for 24 hours in the Invitrogen *Bigfoot* Spectral Cell *Sorter.* Sorted cells were seeded in NC medium without doxycycline.

For induction experiments, 250.000 cells/well in 12 well plates were seeded and the next day doxycycline (1µg/mL) was added to induce expression of p14 and p16. GFP and mCherry expression were analyzed at 24 hours, 5 days, and 8 days post-induction by flow cytometry using BD LSR Fortessa SORP and BD FACSDiva 6.2 software.

**Pemetrexed treatment**

1KO and 2KO NC cell lines (1750 cells/well) and GM00662 *NF1*(+/−) control fibroblasts (750 cells/well) were seeded into 384-well plates (Greiner 781091). 24 hours later, Pemetrexed (LY231514; MedChemExpress) was added in triplicate at seven concentrations using a 1:3 serial dilution starting from 30 µM. After 96 hours of treatment, cell viability was measured using the CellTiter-Glo Luminescent Cell Viability Assay (Promega), and luminescence was quantified with a GloMax Explorer plate reader (Promega). For morphological and senescence assessment, 1KO and 2KO NC cells (15,000 cells/well) were seeded onto chamber slides and incubated overnight. Cells were treated with 450 nM Pemetrexed for 96 hours, and then senescence was evaluated using the Senescence Detection Kit (Sigma-Aldrich, CS0030) according to the manufacturer's protocol. Phase-contrast images were captured using a DMI 6000B microscope (Leica) and LAS X imaging software (Leica).

**Extended qHTS and CRC**

Four concentrations of each compound were plated on different quadrant of one 1536-well plate, leaving column 1-4 available for control compounds. A compound transfer station equipped with a 384-pin tool was used to perform two transfers of 60 nL of compound from each quadrant to a 384-well plate containing spheroid cultures with 30 µL medium, such that columns 3–23 of the 384-well cell plate contained library compounds. Column 1 was reserved for vehicle control (DMSO), column 2 included positive controls—either Panobinostat or UCN-01 (final concentration: 40 µM), and column 24 (rows B, D, F, H, J, L, N, P) contained an 8-point, threefold serial dilution of JQ1 (highest final concentration: 40 µM).

For the cell viability assessed by the CellTiter-Glo® 3D Cell Viability Assay (Promega), 30 µL (equal volume as media in the well) of CellTiter-Glo 3D reagent was added to each well, plates were vortexed for 5 minutes, incubated for 25 minutes at room temperature in the dark. Luminescence was recorded using PHERAstar FSX plate reader (BMG LABTECH).

**Live/dead cell viability assay**

To assess cell viability and morphological changes at 48 hours post-treatment in the HTS, a Live/dead staining assay was performed. A 2X staining solution was prepared by mixing DMEM/F12 (1:1) basal medium supplemented with Hoechst 33342 (1:1000; Thermo Fisher Scientific, cat. no. 62249), propidium iodide (1:500; Thermo Fisher Scientific, cat. no. P3566), and Calcein-AM (1:1000; Thermo Fisher Scientific, cat. no. C1430). This solution was added to each well containing treated spheroids at a 1:1 volume ratio. Spheroids were incubated for 30 minutes at 37°C and 5% $CO_2$. Following incubation, 10X confocal images were acquired using an Opera Phenix™ High-Content Screening System (PerkinElmer).

**Compound Combination Matrix Screening**

Forty-eight candidate compounds from the second single agent were evaluated in combination with Selumetinib and Ribociclib in 3KO NC spheroids (n = 1). The 13 most promising candidates in combination with Selumetinib were selected for further validation in 3KO NC spheroids (n = 3). To assess potential off-target cytotoxicity, these 13 combinations were also tested in control fibroblast (GM00622) 3D spheroids (n = 1).

For matrix screening, cells were seeded into 384-well ULA round bottom plates with 30 µL medium per well. After 24 hours post-seeding, two compounds (120 nL each) were acoustically dispensed using Echo 655 Liquid Handling System (Beckman Coulter), into one well with spheroids, in a threefold dilution 6 X 6 blocks of dose-response matrices. Spheroids were treated with compound combinations for 48 hours, after which cell viability was assessed using the CellTiter-Glo® 3D Cell Viability Assay (Promega), following the protocol described above.

**DNA methylation**

DNA methylation was produced at Diagenode (Ougrée, Belgium). In short, DNA profiles were generated using the Infinium MethylationEPIC (850k) BeadChip array (Illumina, San Diego, USA) according to the manufacturer's instructions. The data was processed, normalized and filtered using minfi Bioconductor package.[6] We then represent the distribution of the samples according its processed beta values using a PCA.

**Single-cell RNA-Seq and analysis**

scRNA-seq analysis was performed at CNAG (Barcelona, Spain). Cryopreserved samples (4 nerves, 4 PNFs, 4 ANNUBPS and 4 MPNSTs) were thawed and digested with 160 U/mL Collagenase Type 1 (Worthington,Lakewood, NJ) and 0.8 U/mL Dispase (Worthington, Lakewood, NJ) for 16 hours at 37ºC 5%$CO_2$. For PNFs, ANNUBPS and MPNSTs the Dead Cell Removal MicroBeads kit (Miltenyi Biotec) was used to eliminate dead cells. For nerve samples the Myelin Removal kit (Milteny Biotec) was used to eliminate myelin and all procedure from this point was conducted at 4ºC. Cells were then resuspended in DMEM + 10% FBS (Gibco) + 1x GlutaMAX (Gibco), filtered with a 40 mm filter and cell viability was calculated with the TC20TM Automated Cell Counter (Bio-Rad). scRNA-seq libraries were built following standard protocols and sequenced using an Illumina platform. FASTQ files were processed with cellranger-arc version 2.0.1 (10X Genomics) using the 2020-A-2.0.0 reference based on GRCh38 and Ensembl genes and multiple samples were combined using the cellranger-arc aggr command with the option "–normalize=depth". Merged UMIs per gene per cell matrix was then loaded into R. We performed quality check, filtering (low library size, high mitochondrial content, low number of features) and normalization using scater.[7] We removed highly stressed cells from one sample (scYoga) and doublets (scDblFinder).[8] Data was batch corrected with fastMNN from package batchelor.[9] Cell types were assigned with SingleR[10] based on the HumanPrimaryCellAtlas reference.[11] For the graphical representation we performed a dimensionality reduction step with PCA and T-distributed Stochastic Neighbor Embedding (TSNE).

**WGS**

The WGS was produced at BGI (Shenzhen, China). In short, the libraries were prepared following standard DNBseq protocols, sequenced in a BGISEQ-500 to a median of 881 million 150-bp paired-end reads per sample, and mapped with BWA-MEM[12] against the GRCh38 genome. WGS data was processed as described elsewhere.[1] In summary, small nucleotide variants were called with Strelka2 and annotated with annovar (Wang et al., 2010; Kim et al., 2018). We called copy-number variation (CNV) using CNVkit[15] with the recommended settings for WGS. As recommended by CNVkit developers, we created a panel of normals and we compared our samples with it. To obtain the exact copy number profile of each sample we used the threshold method, and we provided the Strelka2 germline results for the detection of loss of heterozygosity (LOH) regions. To adjust the thresholds, we calculated the purity of the samples considering the 2n calling and a pseudo-B Allele Frequency (BAF) obtained from the Strelka2 germline results using the loadSNPDataFromVCF function from CopyNumberPlots R package (10.18129/B9.bioc.CopyNumberPlots). The purity calculation was performed using an in-house function that also considers copy-neutral LOH regions or heterozygous loss regions and pseudo-BAF output. Once the purity was calculated, we estimated the ploidy and manually selected the most accurate CNV calling for each sample. Copy number profiles were plotted using the CopyNumberPlots (10.18129/B9.bioc.CopyNumberPlots) and karyoploteR R packages.[16]
